# Polygenic adaptation after a sudden change in environment

**DOI:** 10.1101/792952

**Authors:** Laura K. Hayward, Guy Sella

## Abstract

Polygenic adaptation is thought to be ubiquitous, yet remains poorly understood. Here, we model this process analytically, in the plausible setting of a highly polygenic, quantitative trait that experiences a sudden shift in the fitness optimum. We show how the mean phenotype changes over time, depending on the effect sizes of loci that contribute to variance in the trait, and characterize the allele dynamics at these loci. Importantly, we describe the two phases of the allele dynamics: a rapid phase in which directional selection introduces small frequency differences between alleles whose effects are aligned with or opposed to the shift, which ultimately lead to small differences in their probability of fixation during a second, longer phase, governed by stabilizing selection. As we discuss, our key results should hold in more general settings, and have important implications for efforts to identify the genetic basis of adaptation in humans and other species.

## Introduction

Many traits under natural selection are quantitative, highly heritable, and genetically complex, meaning that they take on continuous values, that a substantial fraction of the population variation in their values arises from genetic differences among individuals, and that this variation arises from small contributions at many segregating loci. It therefore stands to reason that the responses to changing selective pressures often involve adaptive changes in such traits, accomplished through changes to allele frequencies at the many loci that affect them. In other words, we should expect polygenic adaptation in complex, quantitative traits to be ubiquitous. This view traces back to the dawn of population and quantitative genetics (Wright, 1931; Fisher, 1958) and is supported by many lines of evidence (Walsh and Lynch, 2018; Sella and Barton, 2019).

Notably, it is supported by studies of the response to directional, artificial selection on many traits in plants and animals in agriculture and in evolution experiments (Walsh and Lynch, 2018; Sella and Barton, 2019). In these settings, selected traits typically exhibit amazingly rapid and sustained adaptive changes (Weber and Diggins, 1990; Barton and Keightley, 2002; Hill, 2016), which are readily explained by models in which the change is driven by small shifts in allele frequencies at numerous loci (Weber and Diggins, 1990; Hill and Kirkpatrick, 2010), and inconsistent with models with few alleles of large effects (Barton and Keightley, 2002; Zhang and Hill, 2005). The potential importance of polygenic adaptation has also been highlighted by more recent efforts to elucidate the genetic basis of adaptation in humans. In the first decade after genome-wide polymorphism datasets became available, this quest was largely predicated on the monogenic model of a hard selective sweep (Smith and Haigh, 1974; Kaplan et al., 1989), in which adaptation proceeds by the fixation of new or initially rare beneficial mutations of large effects (e.g., Voight et al., 2006). Subsequent analyses, however, echoed studies of artificial selection in indicating that hard sweeps were rare, at least over the past ~500, 000 years of human evolution (Coop et al., 2009; Hernandez et al., 2011). Yet humans plausibly adapted in myriad ways during this time period, and they definitely experienced substantial changes in selection pressures, notably during more recent expansions across the globe. These considerations refocused the quest for the genetic basis of human adaptation on polygenic adaptation (Pritchard et al., 2010; Pritchard and Di Rienzo, 2010).

Findings from genome wide association studies (GWASs) in humans have been central to this research program. Statistical analyses of GWASs indicate that in humans, heritable variation in complex traits is highly polygenic (Loh et al., 2015; Shi et al., 2016; Boyle et al., 2017). For example, for many traits, estimates of the heritability contributed by chromosomes are approximately proportional to their length (Shi et al., 2016), suggesting that the contributing variants are numerous and roughly uniformly distributed across the genome. Such findings reinforced the view that adaptive changes to quantitative traits are likely to often be highly polygenic, but also implied that their identification would be difficult, as the changes to allele frequencies at individual loci may be minute. To overcome this limitation, recent studies pooled signatures of frequency changes over the hundreds to thousands of alleles that were found to be associated with an increase (or decrease) in a given trait (Turchin et al., 2012; Berg and Coop, 2014; Robinson et al., 2015; Field et al., 2016; Berg et al., 2017; Edge and Coop, 2019; Speidel et al., 2019). Initial studies suggest that polygenic adaptation has affected multiple human traits, but these conclusions have been called into question with the realization that the results are highly sensitive to systematic biases in GWASs, most notably due to residual population structure (Berg et al., 2019; Sohail et al., 2019).

Given that polygenic adaptation is plausibly ubiquitous, yet likely hard to identify, there is a clear need for a deep understanding of its behavior in populations and footprints in data. To date, theoretical work has primarily focused on two scenarios. The first is motivated by the observed responses to sustained artificial selection, modeled either as truncation selection (Robertson, 1960) or as stabilizing selection, with the optimal phenotype moving at a constant rate in a given direction (e.g., Bürger and Lynch, 1995; Bürger, 1999; Kopp and Hermisson, 2009; Matuszewski et al., 2015; Jain and Devi, 2018). In natural populations, however, quantitative traits are unlikely to be subject to long-term continuous change in one direction. Instead, considerable evidence indicates that they are often subject to long-term stabilizing selection (Sella and Barton, 2019), with intermittent shifts of the optimum in different directions. The second scenario therefore assumes that a sudden change in the environment induces an instantaneous shift in the optimum of a trait under stabilizing selection (Lande, 1976; Barton et al., 2009; de Vladar and Barton, 2014; Jain and Stephan, 2015; Bod’ová et al., 2016; Jain and Stephan, 2017b; Stetter et al., 2018; Thornton, 2019). Although more elaborate scenarios (where, for example, the optimum and/or strength of stabilizing selection vary frequently) are also possible, this simple scenario provides a sensible starting point for thinking about polygenic adaptation in nature, and is our focus here.

Although there has been considerable work on the adaptive response to an instantaneous change in optimal phenotype, our understanding of this process is still limited. Seminal work by Lande (1976) described the change in the phenotypic mean assuming that phenotypes are normally distributed in the population and that the phenotypic variance remains constant over time. Barton and Turelli (1987) derived recursions for the expected change to higher moments of the phenotypic distribution, and showed that when phenotypic variation arises from alleles with large effect sizes, which are strongly selected and rare, the response to selection introduces skewness in the phenotypic distribution that can substantially affect the change in the phenotypic mean. Their recursions, however, are not generally tractable, and their analyses do not extend to the phenotypic response in more realistic cases, in which phenotypic variation arises from alleles with a wide range of effect sizes. Also, with GWASs now enabling us, at least in principle, to learn about the genetic basis of the phenotypic response, we would like to understand the allele dynamics that underlie it.

Several studies have tackled this problem using simulations (e.g., Stetter et al., 2018; Thornton, 2019). Although illustrative of the dynamics, it is unclear how to generalize their results, given (necessarily) arbitrary choices about multiple parameters and the complexity of these dynamics. In turn, elegant analytical work by de Vladar and Barton (2014) and extensions by Jain and Stephan (2017a,b) afford a general understanding of the allele dynamics in models with an infinite population size. These dynamics, however, are shaped by features of mutation-selection balance that are specific to infinite populations. Notably, they strongly depend on the frequency of alleles prior to the shift in optimum following deterministically from their effect size, and on the critical effect size at which this frequency transitions from being dominated by selection to being dominated by mutation. But in real (finite) populations (including humans), the frequencies of alleles whose selection effects are sufficiently small to be dominated by mutation will be shaped by genetic drift; more generally, variation in allele frequencies due to genetic drift will crucially affect the allele response to selection (see below). Thus, we still lack a solid understanding of the allele dynamic underlying polygenic adaptation in natural populations.

Here, we follow previous work in considering the phenotypic and allelic responses of highly polygenic traits after a sudden change in optimal phenotype. But we do so in finite populations and employ a combination of analytic and simulation approaches to characterize how the responses varies across a broad range of evolutionary parameters.

## The model

We build upon the standard model for the evolution of a highly polygenic, quantitative trait subject to stabilizing selection (Wright, 1935a; Robertson, 1956; Turelli, 1984; Keightley and Hill, 1988; Johnson and Barton, 2005; Simons et al., 2018; Sella and Barton, 2019). An individual’s phenotype is represented by the value of a continuous trait, which follows from its genotype by the standard additive model (Falconer, 1996; Lynch and Walsh, 1998).

Namely, we assume that the number of genomic sites affecting the trait (i.e., the target size) is very large, *L* >> 1, and that an individual’s phenotype is given by

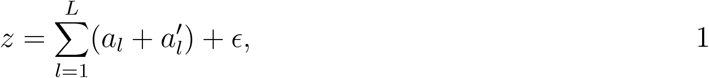

where the first term is the genetic contribution, with *a*_*l*_ and 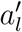 denoting the phenotypic effects of the parents’ alleles at site *l*, and *ϵ* ~*N* (0, *V*_*E*_) is the environmental contribution.

Stabilizing selection is introduced by assuming that fitness declines with distance from the optimal trait value positioned at the origin (*z* = 0). Specifically, we assume a Gaussian (absolute) fitness function:

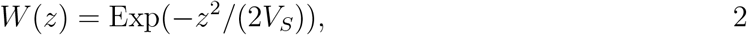

where 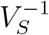 measures the strength of selection. The specific form of the fitness function is unlikely to affect our results under parameter ranges of interest (see below), however. Additionally, since the additive environmental contribution to the phenotype can be absorbed into *V*_*S*_ (by replacing it by 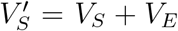; e.g., Turelli (1984); Bürger (2000)), we consider only the genetic contribution.

The population dynamics follow the standard model of a diploid, panmictic population of constant size *N*, with non-overlapping generations. In each generation, parents are randomly chosen to reproduce with probabilities proportional to their fitness (i.e., Wright-Fisher sampling with fertility selection), followed by mutation, free recombination (i.e., no linkage) and Mendelian segregation. We assume that the mutational input per site per generation is sufficiently small such that segregating sites are rarely more than bi-allelic (i.e., that *θ* = 4*Nu* ≪ 1, where *u* is the mutation rate per site per generation). We therefore employ the infinite sites approximation, in which the number of mutations per gamete per generation follows a Poisson distribution with mean *U* = *Lu*. The effect sizes of mutations, *a*, are drawn from a symmetric distribution, i.e., with equal probability of increasing or decreasing the trait value; further assumptions about this distribution are specified below. ***App. A*** Table 1 provides a summary of our notation.

**Table 1.**
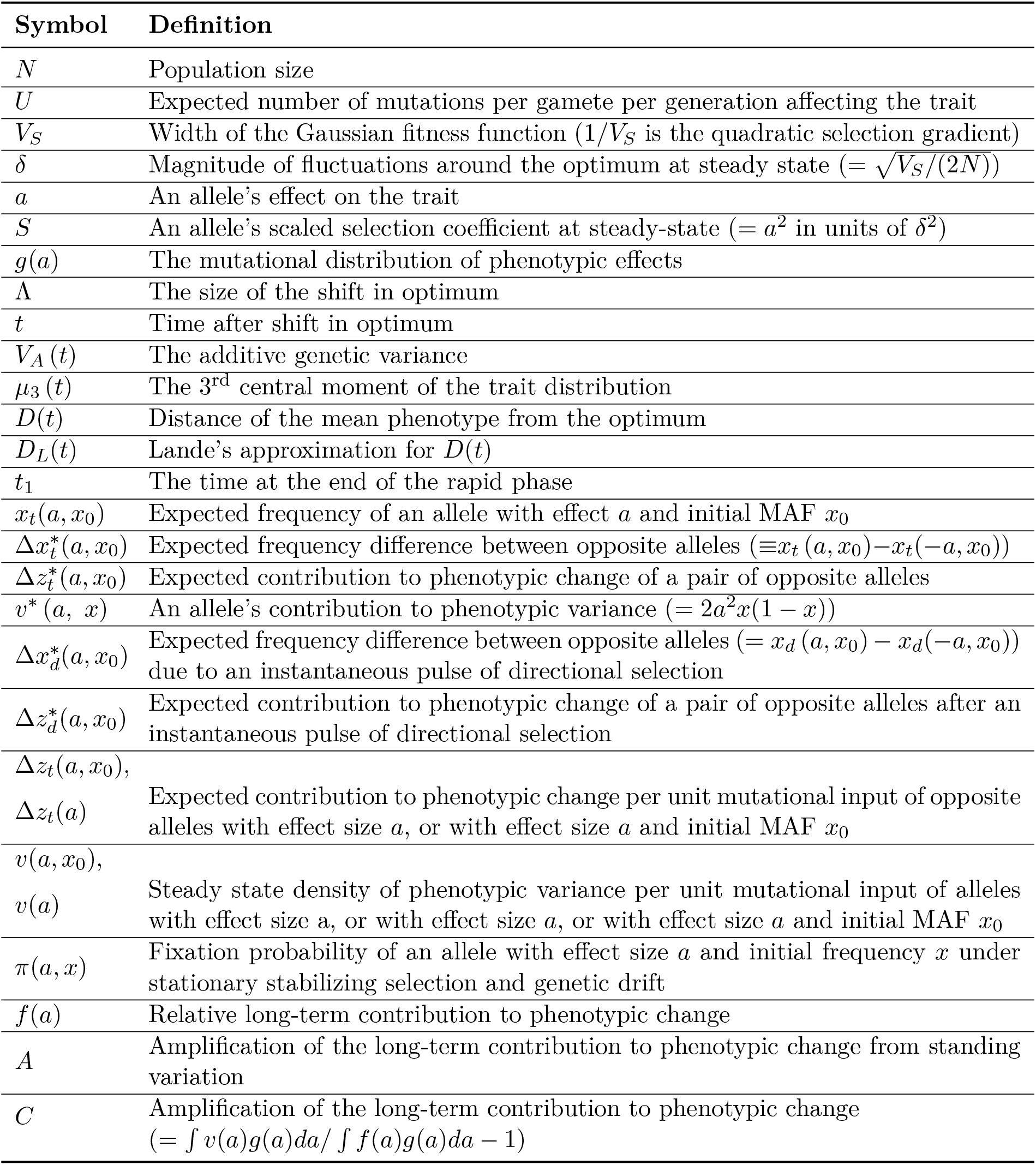
Summary of notation.

### Evolutionary scenario and parameter ranges

We assume that at the outset—before the shift in optimal phenotype—the population has attained mutation-selection-drift balance. We follow previous work on this balance in making several plausible assumptions about parameter ranges (e.g., Simons et al., 2018). First, we assume that the per generation, population scaled mutational input is sufficiently large for variation in the trait to be highly polygenic (specifically, that 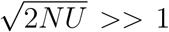). Second, we assume that the expected number of mutations affecting the trait per generation, per gamete, is small (specifically, that *U* = *Lu* ≪ 0.2). For this assumption to be violated in humans, for example, the mutational target size, *L*, would have to exceed ~1.5 Mb (assuming that *u* ≈ 1.25 · 10^*−*8^ per bp per generation; Kong et al. (2012); Besenbacher et al. (2016)). We believe that, when the model is extended to account for effects of pleiotropy, our results should still hold qualitatively for substantially greater values of *U*, but this extension is beyond the scope of this paper. Third, we make the standard assumption that selection coefficients of all alleles satisfy *s* ≪ 1, which implies that the equilibrium selection coefficient *s*_*e*_ = *a*^2^*/V*_*S*_ ≪ 1 (see below and Wright, 1931, 1935b; Turelli, 1984). Fourth, we assume that a substantial proportion of mutations are not effectively neutral, i.e., have 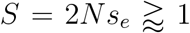. This assumption is supported by empirically based estimates of persistence time for a variety of traits and taxa (Walsh and Lynch, 2018; Sella and Barton, 2019) and by inferences based on human GWASs (Simons et al., 2018; Zeng et al., 2018), indicating that quantitative genetic variance is not predominantly neutral. Under these assumptions, the phenotypic distribution at mutation-selection-drift balance is symmetric and tightly centered on the optimal phenotype (***Fig. 1***). Specifically, the mean phenotype exhibits tiny, rapid fluctuations around the optimal phenotype with variance *δ*^2^ = *V*_*S*_/(2*N*) (Simons et al., 2018); the phenotypic standard deviation is considerably greater than these fluctuations, i.e., 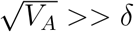 (*SI Section 2.2*); but the phenotypic variance is much smaller than the curvature of the fitness function, i.e., *V*_*S*_ >> *V*_*A*_ (Simons et al., 2018).

**Figure 1.**
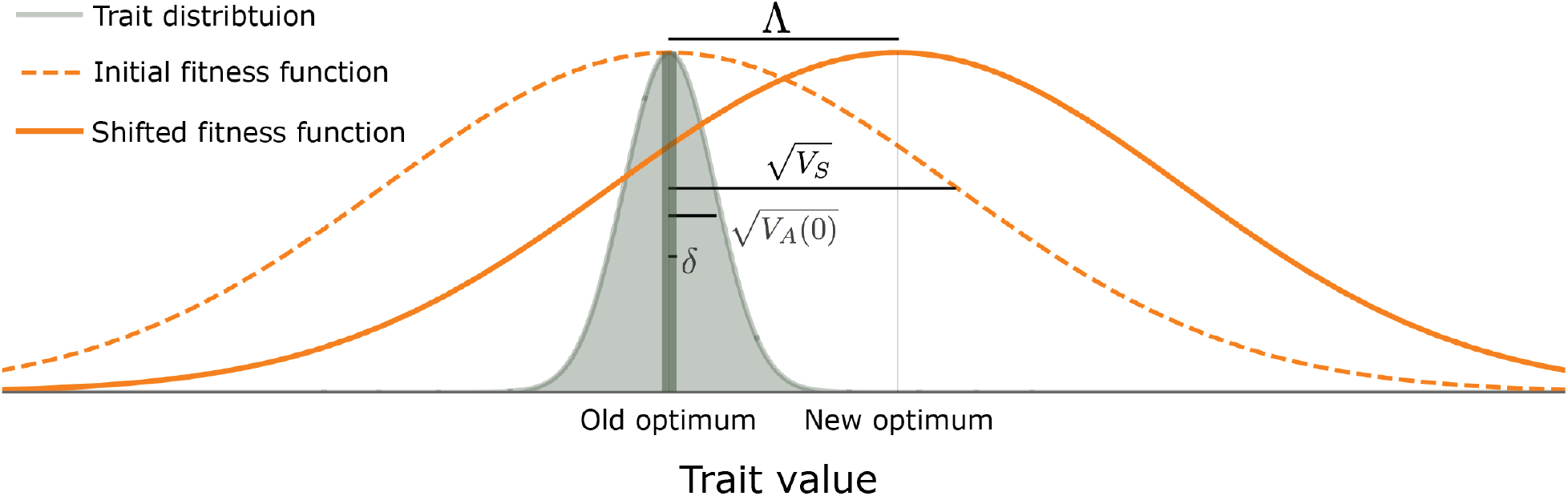
The evolutionary scenario. Before the shift in optimum, phenotypes are distributed symmetrically, with a mean that is very close to the old optimum and a variance that is much smaller than the curvature of the fitness function (*V*_*A*_(0) ≪ *V*_*S*_). We consider the response to an instantaneous shift in optimum, for the case where the magnitude of the shift is smaller than the width of the fitness function 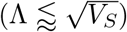. See text for further details.

We consider the response to an instantaneous shift of Λ in optimal phenotype at time *t* = 0 (***Fig. 1***). We assume that the shift in optimum is greater than the equilibrium fluctuations in mean phenotype, i.e., that Λ > *δ*. We further assume that 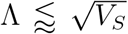 and 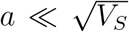. The latter requirements ensure that the maximal directional selection coefficients of alleles, which are attained immediately after the shift, satisfy *s*_*d*_ = 2 · (Λ · *a*)*/ V*_*S*_ ≪ **1** (see below and Wright, 1931; Barton and Turelli, 1987). The requirement that 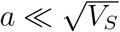 is not particularly restrictive, as it allows for the selection coefficients of alleles at equilibrium, *s*_*e*_ = *a*^2^*/V*_*S*_, to be as large as 1%. Neither is the assumption that 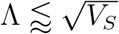, because, given that the genetic variance before the shift satisfies *V*_*A*_(0) *V*_*S*_, it allows for shifts of several equilibrium phenotypic standard deviations (*SI Section 2.2* and ***Fig. S2.2***). Lastly, we assume that the shift is not massive relative to the equilibrium phenotypic standard deviation, specifically that 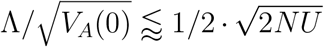. This condition ensures that adaptation to the new optimum requires only a small average frequency change per segregating site (see *SI Section 2.2*). Given that we assume that the trait is highly polygenic, specifically that 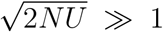, this condition allows for shifts of several equilibrium phenotypic standard deviations. Our assumptions on parameter values and their rational are summarized in ***App. A Table 2***

**Table 2.** Summary of assumptions on parameters.

| Assumption | Interpretation |
| --- | --- |
| $\sqrt{2N\bar{U}} \gg 1$ | High polygenicity |
| $U = Lu \ll 0.2$ | Mutation rate per gamete per generation is substantially smaller than 0.2 |
| $a/\sqrt{V_S} \ll 1$ | Allele effect sizes are substantially smaller than the width of the fitness function |
| $g(a)$ | A substantial proportion of incoming mutations are not effectively neutral |
| $g(a)$ with a substantial portion satisfying $a^2 \gtrsim 1$ | A substantial proportion of incoming mutations are not effectively neutral |
| $g(a) = g(-a)$ | The number of trait-increasing mutations is equal to the number of trait-decreasing mutations |
| $\Lambda > \delta$ | The shift is larger than the expected steady-state fluctuations in mean phenotype |
| $\Lambda \lesssim \sqrt{V_S}$ | The shift is not much larger than the width of the fitness function |
| $\Lambda/\sqrt{V_A(0)} \lesssim 1/2 \cdot \sqrt{2N\bar{U}}$ | The shift is not massive relative to the initial phenotypic standard deviation |

### Choice of units

Our analysis allows us to choose the units in which we measure the trait. When we study the allelic response, we use units based on the dynamics at mutation-selection-drift balance (before the shift in optimum). The population-scaled selection coefficient at steady-state is 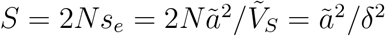; here, the effect size *ã* is measured in arbitrary units, and 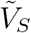 is measured in these units-squared, such that the scaled selection coefficient has no units. We will measure the trait in units of *δ*. In these units, the effect size 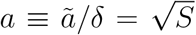, the stabilizing selection parameter 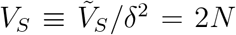, and an allele’s contribution to variance is *v*^*∗*^(*a, x*) = 2*a*^2^*x*(1 − *x*) (and has units of *δ*^2^). We also measure the distance between the mean and optimal phenotype, *D*, and shift in optimum, Λ, in units of *δ*. As shown below, stating our results in these terms makes their form invariant with respect to the population size, *N*, and the strength of stabilizing selection, 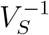.

### Simulations and resources

We compare our analytical results to three layers of simulations (see *SI Section 5* for further detail). The first realizes the full model described above. It is run for a burn-in period of 10*N* generations before the shift and for a period of 12*N* generations after, to attain steady-state both before and after. The second traces all alleles (AA) rather than individuals. It assumes linkage equilibrium (rather than free recombination), and changes to allele frequencies every generation are modeled according to the diffusion approximation detailed below. It is also run with a burn-in period of 10N generations before the shift and for 12*N* generations after. The third kind of simulation traces the dynamic of one allele at the time (OA). To that end: i) when we simulate the selection response from standing variation (as opposed to new mutations), we sample initial minor allele frequencies from the closed form, equilibrium distributions (***Eq. S2.11***), using importance sampling based on the density of variance contributed by different minor allele frequencies (*SI Section 2.1*); and ii) the change in the population’s mean phenotype over time, on which the allele dynamics depend, is given as input, based on either an analytical approximation (see below) or on an average over simulations of the second layer. The OA simulation is run until the focal allele fixes or goes extinct. The last two layers allow for greater computational tractability, and we validate our main results against simulations from the first layer (***Figs. S5.1*** and ***S5.2*** and *SI Section 5*). Documented code for simulations, numerical analysis, and graphs can be found at https://github.com/sellalab/PolygenicAdaptation1D.

## Results

### Phenotypic response

We first consider how the population’s mean phenotype approaches the new optimum. In *SI Section 1.2*, we express the mean distance from the new optimum, *D*(*t*), as a sum over alleles’ contributions. We show that under our assumptions, the expected, per generation change in this distance is well approximated by

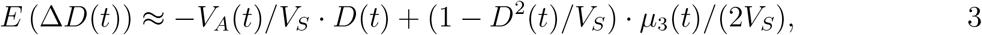

where *V*_*A*_(*t*) and *µ*_3_(*t*) denote the 2^nd^ and 3^rd^ central moments of the phenotypic distribution. Similar expressions were derived by Barton and Turelli (1987) under the rare-alleles approximation and by Bürger (1991) under the assumption of a parabolic fitness function.

We rely on ***Eq. 3*** to describe the phenotypic response to selection. This response takes a simple form in the infinitesimal limit, in which genetic variation at equilibrium arises from infinitely many segregating alleles with infinitesimal effect sizes (Fisher, 1919; Bulmer, 1980; Barton et al., 2017). In this limit, the change in mean phenotype is achieved by infinitesimal changes to allele frequencies at infinitely many loci, without changing the frequency distribution. Consequently, the phenotypic distribution remains normal and the phenotypic variance remains constant. Under these assumptions, ***Eq. 3*** reduces to

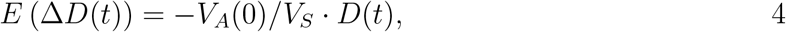

which (in continuous time) is solved by

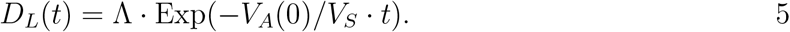

This solution was first derived by Lande (1976), and we refer to it henceforth as Lande’s solution or approximation. When genetic variance is dominated by loci with small and intermediate effect sizes (as defined below), the trait is highly polygenic, and the shift in optimum is not too large (relative to the phenotypic standard deviation), changes to the 2^nd^ and 3^rd^ moments of the phenotypic distribution are small and the expected phenotypic response is well approximated by Lande’s solution (***Figs. 2A, S7.2*** and ***S7.3***).

**Figure 2.**
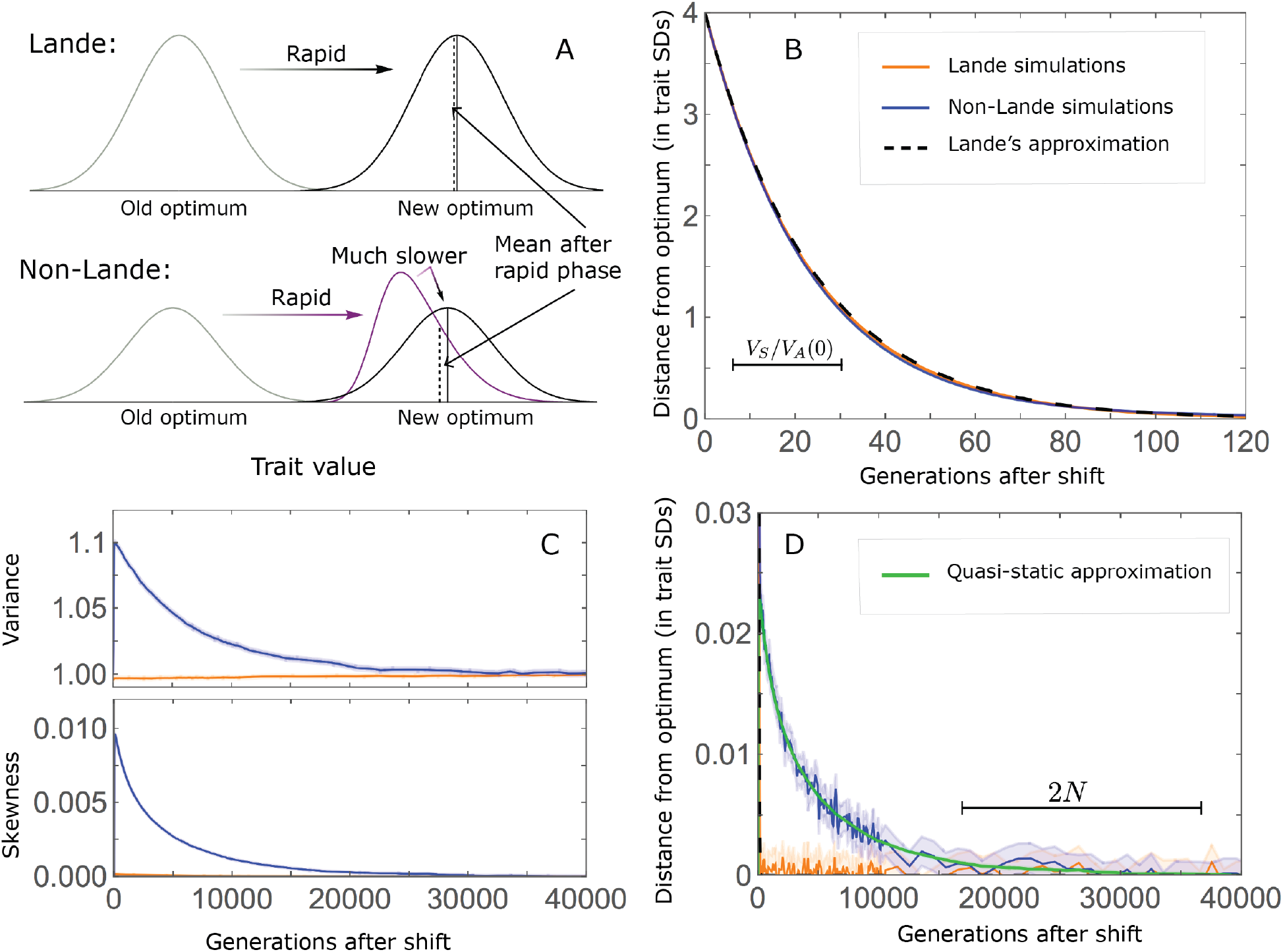
The phenotypic response to a shift in optimal phenotype. A) Cartoon of the two kinds of phenotypic response: i) the Lande approximation, in which the mean approaches the new optimum exponentially with time and the phenotypic distribution maintains its shape; ii) substantial deviations from Lande’s approximation, in which the mean approaches the new optimum rapidly at first, but during this time the phenotypic distribution becomes skewed, causing the mean’s approach to slow down dramatically, to a rate that is dictated by the decay of the 3^rd^ moment. B) In both the Lande and non-Lande cases, the mean phenotype initially approaches the new optimum rapidly. This approach is described by Lande’s approximation, and thus almost identical in the two cases (which is why only the Lande curve is visible). C) In the non-Lande case, the phenotypic variance and skewness increase during the rapid phase and then take a very long time to decay to their values at steady state. D) Over the longer-term, the approach to the optimum in the non-Lande case almost grids to a halt, where its rate can be described by the quasi-static approximation (***Eq. 6***). The simulation results in B-D were averaged over 2500 runs of our allele simulations (AA) (see section on *simulations and resources* and *SI Section 5*), with 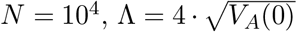, and 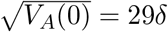. In the Lande case, squared effect sizes were Gamma distributed with *E (a*^2^)= *V (a*^2^)= 1 (*S* = *a*^2^ ~Γ(1, 1)) and *U* ≈ 0.03 (to match *V* (0) in the non-non-Lande case,) in the non-Lande case, squared effect sizes were Gamma distributed with *E (a)*^2^ = 16 and *V (a)*^2^ = 256 (*a*^2^~Γ(1, 16)) and *U* = 0.01.

When alleles with large effects contribute markedly to genetic variance, the trait is not highly polygenic or the shift in optimum is large, changes to the 2^nd^ and 3^rd^ moments of the phenotypic distribution and consequently deviations from Lande’s approximation become more substantial (***Figs. 2C***, ***S7.2*** and ***S7.3***; Barton and Turelli, 1987). In *SI Section 6* we investigate the relationship between model parameters and deviations from Lande’s approximation in more depth. For intuition, consider a pair of minor alleles with the same initial frequency and magnitude of effect, where the effect of one is aligned with the shift and the effect of the other opposes it. After the shift, directional selection increases the frequency of the aligned allele relative to that of the opposing one. The frequency increase of the aligned allele increases variance more than the frequency decrease of the opposing allele decreases it, resulting in a net increase to variance (***Fig. 2C***; Barton and Turelli, 1987; de Vladar and Barton, 2014; Jain and Stephan, 2017a). The relative changes in frequency and thus the net increase in variance are greater for alleles with larger effects. Next consider the 3^rd^ moment. At steady-state, the contributions of alleles with opposing effects to the 3^rd^ moment cancel out. After the shift, the frequency increase of aligned alleles relative to opposing ones introduces a non-zero 3^rd^ moment (***Fig. 2C***). Large effect alleles contribute substantially more to this 3^rd^ moment, plausibly because their individual, steady-state contribution to the 3^rd^ moment is greater to begin with (see *SI Section 2.3* and ***Fig. S2.1B***) and because they exhibit larger relative changes in frequency after the shift (see section on *the allelic response in the rapid phase* and *SI Section 3*). By the same token, with larger shifts and lower polygenicity, directional selection causes greater relative frequency differences between alleles with opposing effects, and thus greater increases to variance and a greater skew of the phenotypic distribution.

The increase in 2^nd^ and 3^rd^ moments after the shift result in a phenotypic dynamic with two distinct phases. First, immediately after the shift, the mean phenotype rapidly approaches the new optimum, akin to the exponential approach in Lande’s approximation. In this case, however, genetic variance increases and thus the exponential rate of approach may increase, making the expected approach even faster (***Eq. 3***). Shortly thereafter, when the mean phenotype nears the optimum, the decreasing distance and increasing 3^rd^ moment reach the point at which

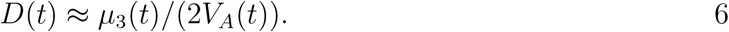

The two terms on the right-hand side of ***Eq. 3*** then approximately cancel out, and the dynamic enters a second, prolonged phase, in which the approach to the optimum nearly grinds to a halt (***Fig. 2D***). During this phase, the expected change in mean phenotype can be described in terms of a quasi-static approximation given by ***Eq. 6*** (***Figs. 2D*** and ***S7.1***). The rate of approaching the optimum is then largely determined by the rate at which the 3^rd^ moment decays. This roughly corresponds to the rate at which the allele frequency distribution equilibrates and steady-state around the new optimum is restored (see section on *other properties of the equilibration process*).

### Allele dynamics

We now turn to the allele dynamics that underlie the phenotypic response. These dynamics can be described in terms of the first two moments of change in frequency in a single generation (Crow and Motoo, 1970; Ewens, 2012). For an allele with effect size *a* and frequency *x*, we calculate the moments by averaging the fitness of the three genotypes over genetic backgrounds (*SI Section 1*). Under our assumptions, the moments are well approximated by

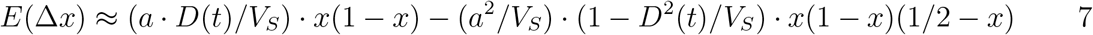

and

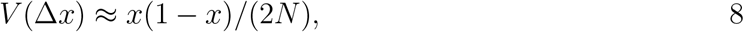

which is the standard drift term. Similar expressions for the first moment trace back to Wright (1935a) and have been used previously to study the response to selection on quantitative traits (Barton, 1986; Bürger, 1991; Charlesworth, 2013; de Vladar and Barton, 2014). The two terms in the first moment reflect different modes of selection: directional and stabilizing, respectively. The first term arises from directional selection on the trait and takes a semi-dominant form with selection coefficient *s*_*d*_ = 2*a D*(*t*)*/V*_*S*_. Its effect is to increase the frequency of alleles whose effects are aligned with the shift (and vice versa) and its strength weakens as the distance to the new optimum, *D*, decreases. The second term arises from stabilizing selection on the trait and takes an under-dominant form with selection coefficient *s*_*e*_ = *a*^2^*/V*_*S*_. (1 − *D*^2^(*t*)*/V*_*S*_). Its effect is to decrease an allele’s contribution to phenotypic variance, 2*a*^2^*x*(1 − *x*), by reducing minor allele frequency (MAF); it becomes weaker as the MAF approaches 1/2.

The relative importance of the two modes of selection varies as the mean distance to the new optimum, *D*, decreases. We therefore divide the allelic response into two phases: a *rapid phase*, immediately after the shift, in which the mean distance to the new optimum is substantial and changes rapidly, and a subsequent, prolonged *equilibration phase*, in which the mean distance is small and changes slowly (Jain and Stephan, 2017a). We define the end of the rapid phase as the time, *t*_1_, at which Lande’s approximation for the distance to the optimum *D*_*L*_ (*t*_1_), equals the standard deviation of the distance from the optimum at steady-state 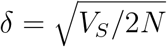, i.e.,

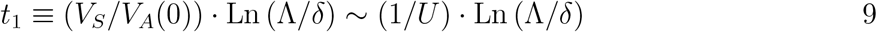

(in *SI Section 2.2* we show that *V*_*S*_*/V*_*A*_(0) ~1*/U*). This definition is somewhat arbitrary, as the transition between phases is gradual, but it roughly captures the change in allele dynamics (***Fig. S7.1***).). Moreover, our analysis is insensitive to this particular choice (we only use in comparing analytic and simulation results for the rapid phase).

The change in mean phenotype during the rapid phase is driven by the differential effect of directional selection on minor alleles whose effects are aligned and opposed to the shift in optimum (***Fig. 3***). Considering a pair of minor alleles with opposing effects of the same size and the same initial frequency, selection increases the frequency of the aligned allele relative to the opposing one. By the end of the rapid phase, the frequency differences across all aligned and opposing alleles drive the mean phenotype close to the new optimum (***Fig. 2A***). Deviations from Lande’s approximation manifest as prolonged, weak directional selection during the equilibration phase, which further increases the expected frequency difference between aligned and opposing alleles. However, given that we are considering a highly polygenic trait, the expected frequency difference between a pair of opposing alleles will be small. This small difference causes aligned alleles to have a slightly greater probability of eventually fixing during the equilibration phase (***Fig. 3***). Over a period on the order of 2*N* generations (see below), the frequency differences between aligned and opposing alleles are replaced by a slight excess of fixed differences between them, and the steady-state genetic architecture is restored around the new optimum. In the following sections, we describe these processes quantitatively. Specifically, we ask how the relative contribution of alleles to phenotypic change during the two phases depends on their effect size and initial frequency.

**Figure 3.**
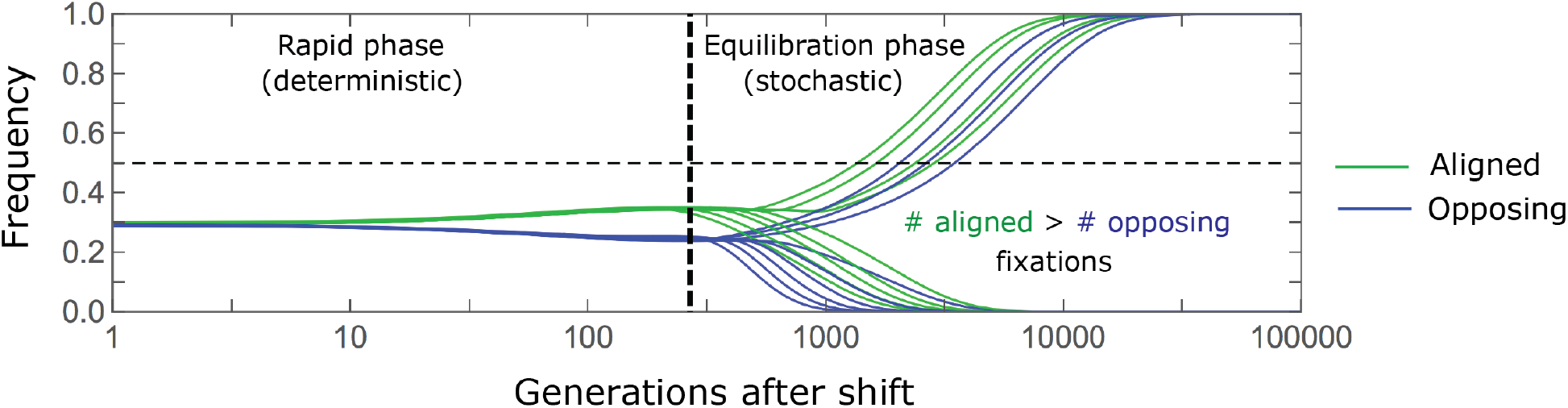
A cartoon of allele dynamics. We divide the allele dynamics into rapid and equilibration phases, based on the rate of phenotypic change, and consider the trajectories of alleles with opposing effects of the same size, which start at the same initial minor frequency. During the rapid phase, alleles whose effects align with the shift slightly increase in frequency relative to those with opposing effects. During the equilibration phase, this frequency difference can increase further and eventually leads aligned alleles to fix with slightly greater probabilities than opposing ones.

### The allelic response in the rapid phase

We can describe changes to allele frequencies during the rapid phase with a simple deterministic approximation. The duration of the rapid phase is much shorter than the time scale over which genetic drift has a substantial effect (*t*_1_ ~1*/U* ≪ 2*N* generations; see ***Eq. 9***), allowing us to rely only on the first moment of change in allele frequency ((***Eq. 7***). Additionally, deviations of the distance *D*(*t*) from Lande’s approximation during this phase have negligible effects (***Figs. 2B*** and ***S3.1D-F***), allowing us to assume that *D*(*t*) = *D*_*L*_(*t*) (***Eq. 5***). Lastly, when relative frequency changes are small, we can use a *linear approximation*in which we substitute the frequency in the first moment by its initial value. With these simplifications, we can integrate the first moment over time to obtain an explicit approximation for frequency changes.

Consider a pair of minor alleles with opposing effects of size *a* and initial frequency *x*_0_ before the shift in optimum. Using our linear approximation, we find that the frequency difference between them at the end of the rapid phase is

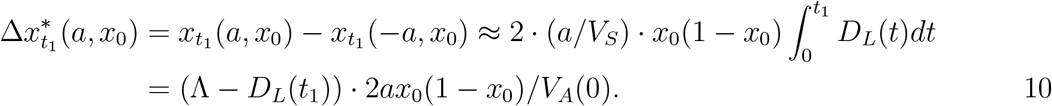

The contribution of the pair to the change in mean phenotype is

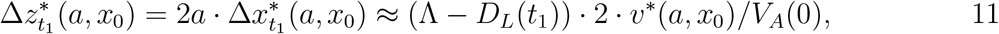

where *v*^*\**^(*a, x*_0_) = 2*a*^2^*x*_0_(1−*x*_0_) is the contribution to variance of an allele with effect size *a* and frequency *x*_0_. Thus, the pair’s contribution to phenotypic change is proportional to its contribution to phenotypic variance before the shift in optimum.

The expected relative contribution of *all* alleles with a given effect size and initial frequency is therefore proportional to their expected initial, steady-state contribution to phenotypic variance. We will focus on the contribution per unit mutational input of alleles with a given effect size. To this end, we measure the trait value in units of 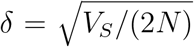 and express allele effect sizes in terms of the scaled selection coefficients at steady-state (when *D* = 0), 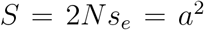 in these units (see *the model* section). Expressing our results in this form makes them invariant with respect to changing the population size, *N*, stabilizing selection parameter, *V*_*S*_, mutational input per generation, 2*NU*, and distribution of effect sizes, *g*(*a*). In these terms, the expected contribution of alleles with given effect size and initial MAF to phenotypic change is

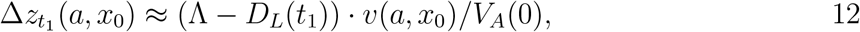

and the marginal contribution of alleles with a given effect size is

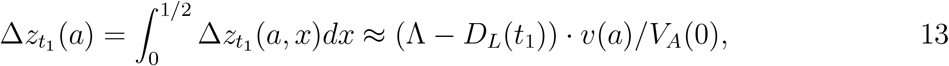

where *v*(*a, x*_0_)≈4*a*^2^ Exp (-*a*^2^*x*_0_(1 -*x*_0_)) and 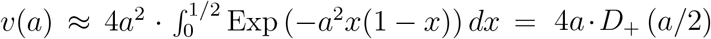 are the corresponding densities of variance per unit mutational input at steady-state, and *D*_+_ is the Dawson function (*SI Section 2.2*). The absolute expected contributions follow from multiplying these expressions by the mutational input per generation, 2*NU ·g*(*a*). Specifically, as we would expect, the total change in mean phenotype during the rapid phase is 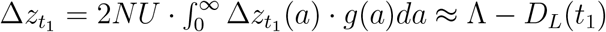 (as 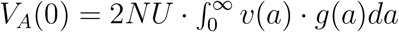).

The relative contribution of alleles with given effect size and initial MAF to phenotypic change follows from their expected contribution to variance at steady-state ((***Eqs. 12*** and ***13***, and ***Fig. 4***). The properties of *v*(*a*) imply that (***Fig. 4A***):i) the relative contribution of alleles with small effect sizes (*a*^2^≪1) scale linearly with *S* = *a*^2^ (*v*(*a*) ≈ 2*a*^2^, measured in units of *δ*^2^); ii) the contribution of alleles with moderate and large effect sizes (roughly *S* = *a*^2^> 3) are much greater, and fairly insensitive to the effect size (with *v* (*a*) ~4); and iii) the contribution is maximized for *a*^2^ ≈ 10 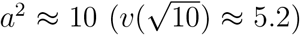 (see Simons et al., 2018, for intuition about these properties). While large and moderate effect alleles make similar contributions to phenotypic change, MAFs of large effect alleles before the shift are much lower than the MAFs of moderate ones (***Fig. 4B***), because they are subject to stronger stabilizing selection. The expected frequency difference between pairs of opposing alleles is greatest for moderate effect sizes (***Fig. 4C***), because it is proportional to *E* (2*ax*_0_(1 −*x*_0_)) (***Eq. 10***), which increases with *v*(*a*)*/a* and *v*(*a*) is similar for moderate and large effect sizes. Additional properties of the allelic response during the rapid phase are presented in *SI Section 3*.

**Figure 4.**
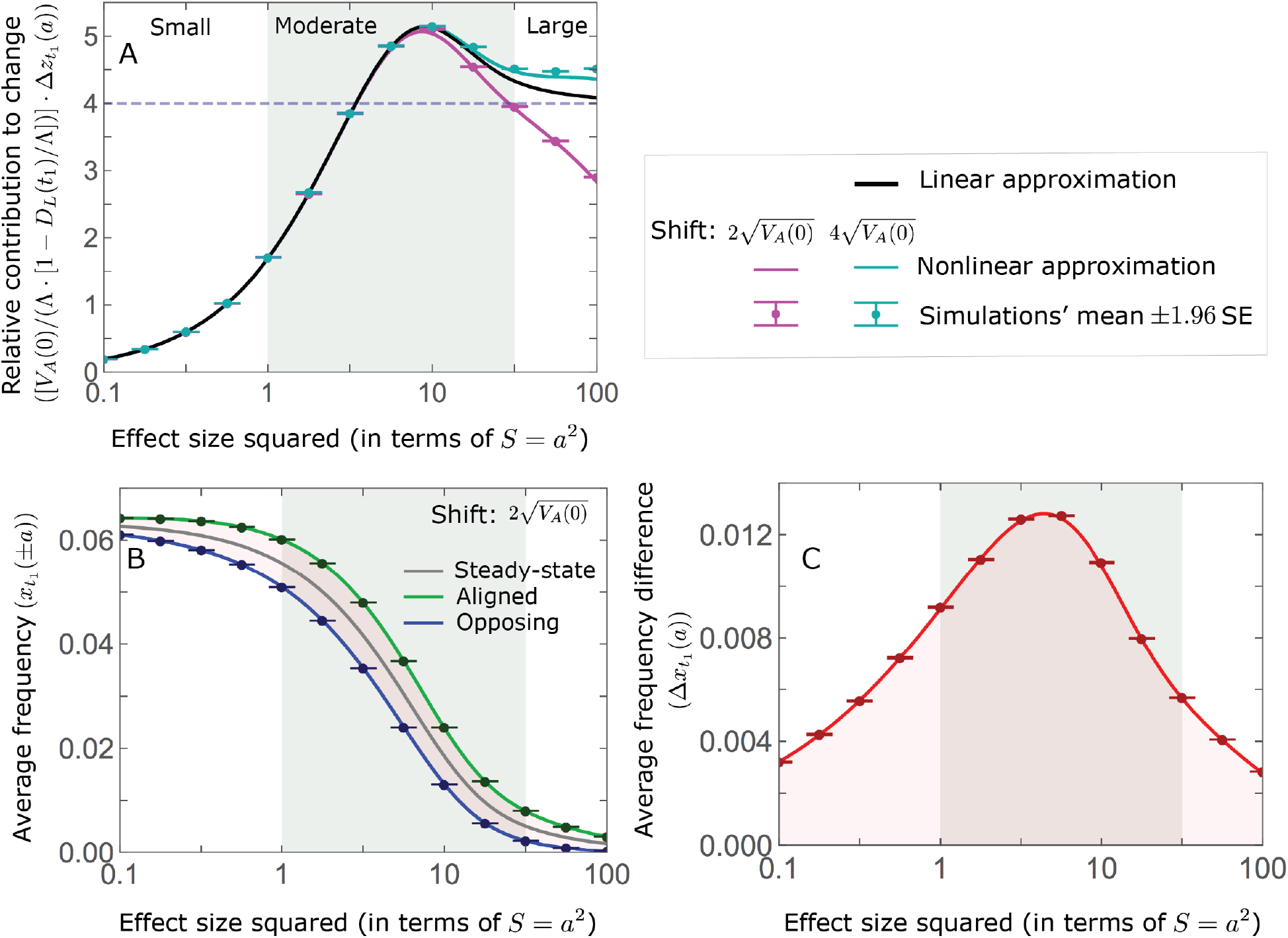
The allelic response during the rapid phase. A) Alleles with moderate and large effects make the greatest contribution to phenotypic change (per unit mutational input). The results of our linear approximation are compared with a more accurate nonlinear one (see *SI Section 3.1.2*) and with simulations. B) The average MAF of aligned and opposing alleles at the end of the rapid phase decreases with effect size. C) The expected frequency difference between pairs of opposing alleles is greatest for moderate effect sizes. Simulation results for each point were averaged over 2 5 10^5^ runs of our one allele (OA) simulation, assuming = 10^4^, Lande’s approximation with 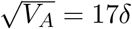, and the shifts specified in the legend.

When the polygenicity is low and/or the shift in optimum or effect sizes are large our linear approximation becomes less accurate (***Fig. 4A***) Specifically, minor alleles exhibit large relative changes in frequency such that substituting the initial MAF for the frequency in ***Eq. 7*** for the 1^st^ moment is inaccurate. In *SI Section 3.1.2* we derive a nonlinear approximation that is more accurate in these cases (***Figs. 4A, S3.2*** and ***S3.3***). Nonetheless, the qualitative behaviors we outlined remain intact.

### The allelic response in the equilibration phase

Over the long run, the small frequency differences between opposite alleles that accrue during the rapid phase translate into small differences in their fixation probabilities (***Fig. 3***).). In the non-Lande case, prolonged weak directional selection during the equilibration phase amplifies these differences in fixation probabilities. We approximate fixation probabilities in two steps. First, we model the effect of directional selection on frequency as an instantaneous, deterministic pulse. Second, we apply the diffusion approximation for the fixation probability, assuming stationary stabilizing selection (*D* = 0), genetic drift, and the initial frequency after the pulse. We further assume that the relative changes in allele frequencies due to directional selection are small, such that we can use approximations that are linear in this change; but in *SI Section 4*, we derive nonlinear approximations that relax this assumption.

### The Lande case

When Lande’s approximation is accurate, directional selection is non-negligible only briefly after the shift. This justifies approximating its effects as if they were caused by an instantaneous pulse. It also suggests that mutations that arise after the shift in optimum contribute negligibly to phenotypic change, because fairly few of them arise when directional selection is non-negligible and their fixation probabilities and thus the difference in fixation probabilities between mutations with opposite effects are tiny (given that they start from an initial frequency of 1/2*N*).

Consider a pair of opposite minor alleles, with effect size *a* and initial frequency *x*_0_. Modeling the effects of directional selection on their frequencies as an instantaneous pulse, and assuming that these effects are small, we find that the resulting frequency differences between them is approximated by

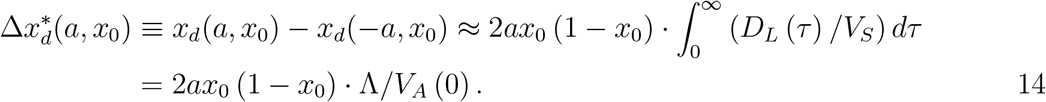

Consequently, the pair’s expected contribution to phenotypic change is approximated by

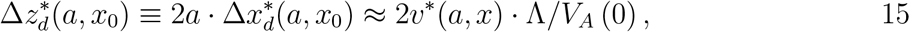

and the contribution of such pairs per unit mutational input is approximated by

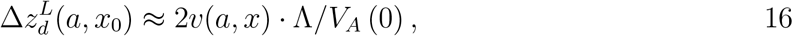

where, as before, *v*^*\**^(*a, x*) = 2*a*^2^*x*(1 − *x*) is an allele’s contribution to genetic variance and *v*(*a, x*) is the steady-state (initial) density of variance per unit mutational input, and we use the superscript *L* to denote that this applies to the Lande case.

We approximate a pair’s expected long-term, fixed contribution to phenotypic change by calculating the difference in fixation probabilities of the opposite alleles given their frequency after the pulse, again assuming that the effects of the pulse are small. Namely,

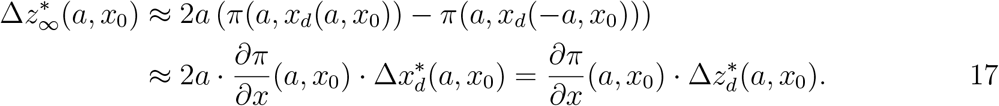

where *π*(*a, x*) denotes the fixation probability of an allele with effect size *a* and initial frequency *x* under stationary stabilizing selection and drift. In *SI Section 2* we derive the diffusion approximation for *π*(*a, x*) and show that

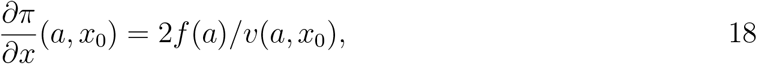

where *f* (*a*) 2*a*^3^ Exp 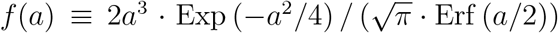. From ***Eqs. 14–18***, we find that the expected fixed contribution per unit mutational input of pairs of alleles is

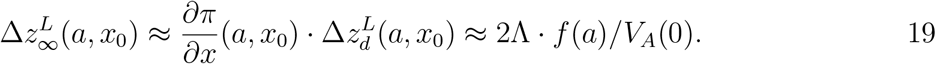

Note that this expression does not depend on the initial frequency! The expected marginal contribution of alleles with a given effect size follows and is

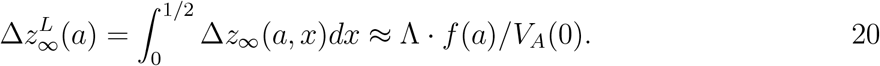

Hence, the function *f* approximates how the relative long-term contribution of alleles depends on their effect sizes (***Fig. 5A***).

**Figure 5.**
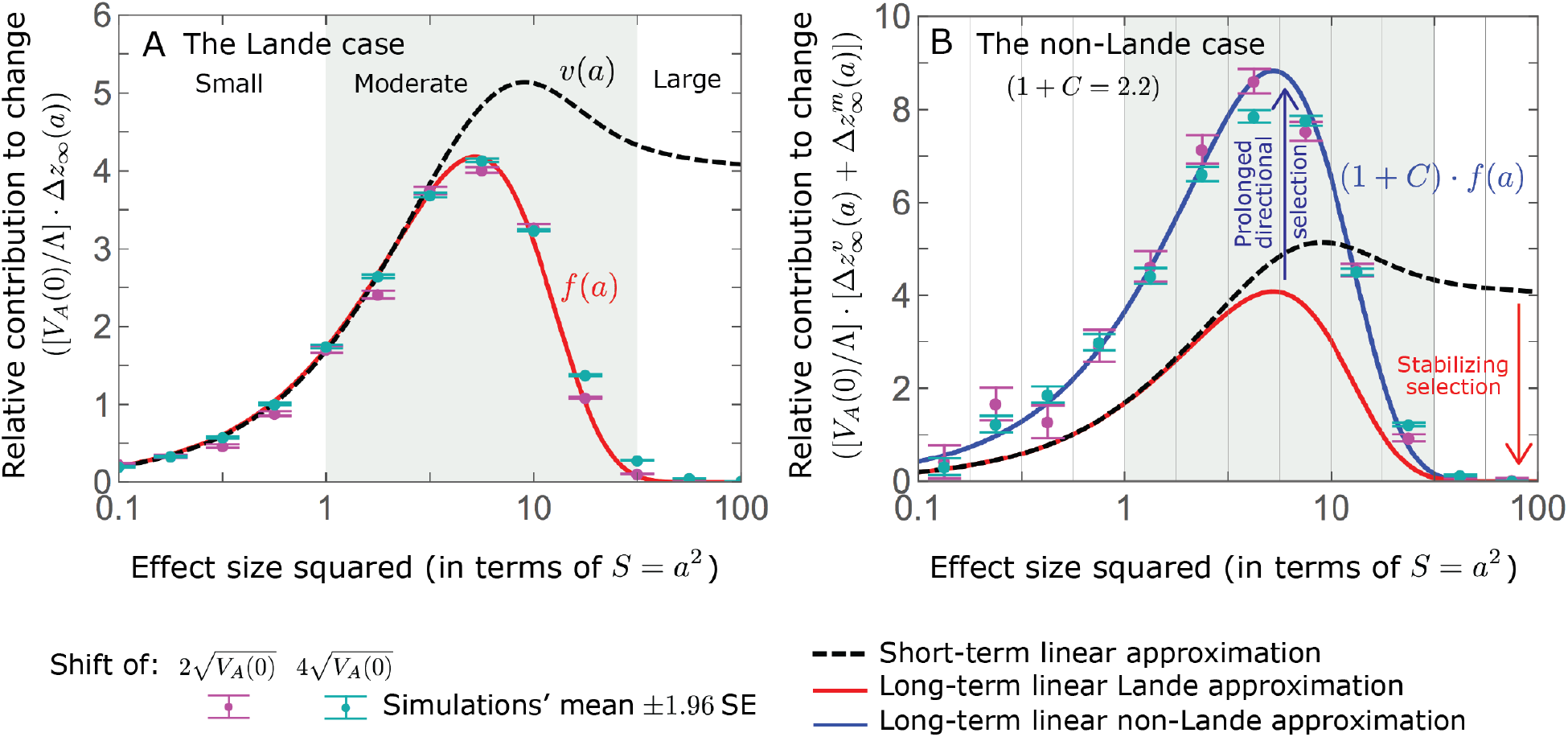
The long-term (fixed) allelic contribution to phenotypic adaptation. We show the relative contribution of alleles as a function of effect size, based on our linear approximations and on simulations with the two shift sizes specified in the caption. A) The Lande case. Our theoretical prediction is described by the function *f* (*a*) (***Eqs. 13*** and ***20***) and simulation details are the same as in ***Fig. 4***. Our prediction for the long-term contribution (corresponding to *f* (*a*)) is always below the prediction for the short-term (corresponding to *v*(*a*)). The difference becomes substantial for *a*^2^ ⪆ 4, implying that the linear Lande approximation underestimates the fixed contribution when large effect alleles contribute markedly to genetic variance. B) The non-Lande case. Here, we assume the same effect size distribution as in ***Fig. 2***, which yields an amplification factor of 1 + *C* ≈ 2.2 (***Eq. 22***). Our theoretical prediction for the joint contribution of standing variation and new mutations is described by the function (1 + C) · *f*(*a*) · Λ/*V*_*A*_(0) (Eq. 28). We calculate the relative contribution of alleles in each effect size bin (between the gray gridlines), by dividing the contribution of all fixations in the bin by the mutation rate per generation corresponding to that bin. In both cases, long-term stabilizing selection diminishes the contribution of alleles with large effects and amplifies that of alleles with small and moderate effects (red arrows). In the non-Lande case, long-term, weak directional selection amplifies the contribution of alleles with small and moderate effects (blue arrow). See ***Figs. S4.1-S4.8*** for other attributes of the long-term allelic response and for the nonlinear approximations.

We expect the total long-term allelic contribution to equal the shift in optimum, Λ. In our linear Lande approximation, the total contribution is

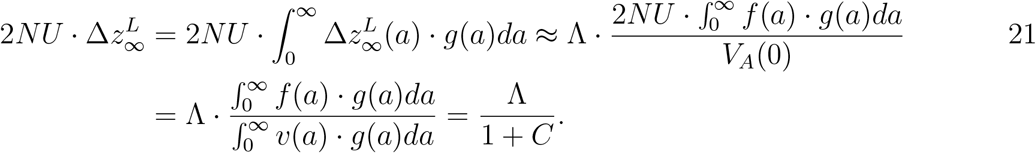

where we note that 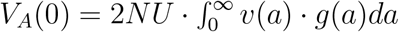and define

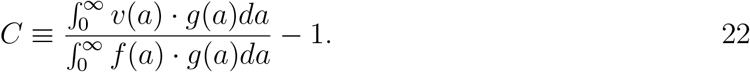

Thus, *C* ≪1 is a necessary condition for our approximation to be accurate. Given that *f* (*a*) *< v*(a) for any *a*, but the difference is substantial only for *a* ⪆ 2 (***Fig. 5A***), this condition implies that the bulk of variance before the shift arises from alleles with *a <* 2. When alleles with larger effects contribute substantially to genetic variance before the shift (and *C* is not negligible) then Lande’s approximation becomes inaccurate. The prevalence of large effect alleles leads to a quasi-static decay of the distance *D* during the equilibration phase (***Fig. 2D*** and the section on *phenotypic response*), and the resulting prolonged, weak directional selection markedly amplifies the difference in fixation probabilities between opposite alleles. Our linear Lande approximation does not account for this amplification, and it therefore underestimates the total long-term allelic contribution.

### The non-Lande case

We can, however, extend our approximation to account for the amplification in the non-Lande case. To this end, we modify our instantaneous pulse approximation for a pair of opposite alleles (***Eq. 14***) to

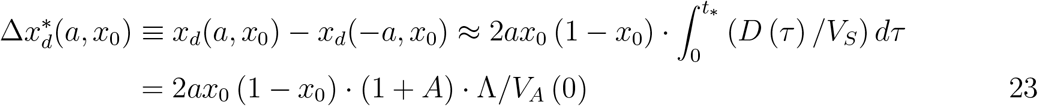

where *t*_*\**_ is the effective number of generations over which an allele is subject to directional selection (because alleles segregate for a finite time) and *A* is defined such that 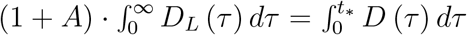 (although the justification for the instantaneous pulse approximation is less obvious in this case, since *t*_*\**_ can be on the order of 2*N* generations; see *SI Section 4.2*). Following the same steps as in the Lande case, we then find that the expected long-term (fixed) contribution per unit mutational input of pairs of alleles with a given effect size and initial MAF is

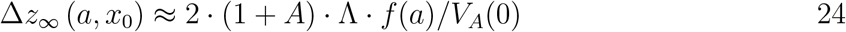

and that their expected marginal contribution for a given effect size is

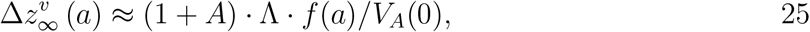

where the superscript *v* denotes that these contributions originate from variation that segregated before the shift in optimum.

In the non-Lande case, the fixation of mutations that arise after the shift in optimum can also contribute substantially (***Fig. S6.2B***), because prolonged, weak directional selection can produce a substantial difference in the numbers of fixations of mutations with opposite effects. In *SI Section 4.2.2*), we follow the same approach that we applied for standing variation to show that the relative long-term contribution of new mutations with a given effect size can be approximated by

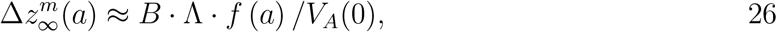

where *B >* 0 (we provide an explicit expression for *B* in *SI Section 4.2.2*). Thus, similar to what we found in the Lande case, the function *f* approximates how the relative long-term contribution of alleles depends on their effect sizes, but here it applies to both standing variation and new mutations (***Fig. 5B***).

To gain further understanding of the non-Lande case, we consider the joint contribution of standing variation and new mutations. Equating the total contribution with the shift in optimum we find that

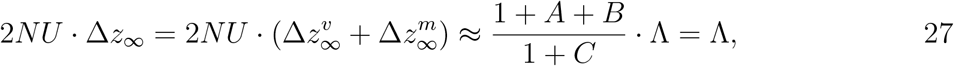

with *C* defined in ***Eq. 22***. This implies that *A* + *B* = *C* and that the proportional contributions of standing variation and new mutations are (1 + *A*)/(1 + C) and B/(1 + C), respectively. It also implies that the contribution per unit mutational input of alleles with a given effect size *a* is

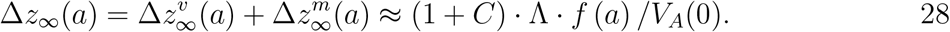

Thus, in the linear approximation, prolonged weak directional selection amplifies the relative contribution of alleles of any given effect size by the same factor of (1 + *C*) (***Figs. 5B*** and ***S4.8***). The amplification *C* is therefore an allelic measure of the deviation from Lande’s approximation (see ***Figs. S6.1*** and ***S6.2***), and, intriguingly, it depends only on the mutational distribution of effect sizes (***Eq. 22***).

When polygenicity is low, the shift in optimum is large or effect sizes are large, our linear approximations become less accurate (*SI Section 4.2* and ***Figs. S4.2*** and ***S4.5***). In these cases, directional selection causes large relative changes in MAFs and using the initial MAF in the instantaneous pulse approximation (i.e., in ***Eqs. 14*** and ***23***) becomes inaccurate. Large changes in frequency also undermine the accuracy of our Taylor approximation of fixation probabilities (***Eq. 17***). In *SI Section 4*, we show that the linear approximations are accurate when *a* (1 + *A*)Λ/*V*_*A*_ (0) ≪ 1 (with *A* = 0 in the Lande case) and we derive nonlinear approximations that are more accurate when this condition is violated (***Figs. 4A, S4.2*** and ***S4.5***). Even in these cases, however, the linear approximations capture the salient features of the long-term allelic contribution to phenotypic adaptation (***Figs. 5*** and ***S4.2-S4.8***).

### Turnover in the genetic basis of adaptation

Notably, our linear approximations capture the dramatic turnover in the genetic basis of adaptation during the equilibration phase (***Fig. 6***). In the long run, the short-term contribution of large effect alleles (*S* = *a*^2^ ⪆30) is almost entirely wiped out, and is supplanted by the contribution of moderate effect alleles (*a*^2^ ≈5) (***Figs. 6A, S4.4*** and ***S4.8***). Moreover, for any given effect size, the proportional long-term contribution of minor alleles that segregated at low frequencies before the shift is diminished relative to their short-term contribution, all the more so for large effect sizes (***Figs. 6B*** and ***S4.3***). For instance, for an effect size *a*^2^ =35, minor alleles with initial frequencies below 0.05 account for more than 99% of the short-term contribution but for only ~ 10% of the (much smaller) long-term contribution (***Figs. 6B*** and ***S4.3***).

**Figure 6.**
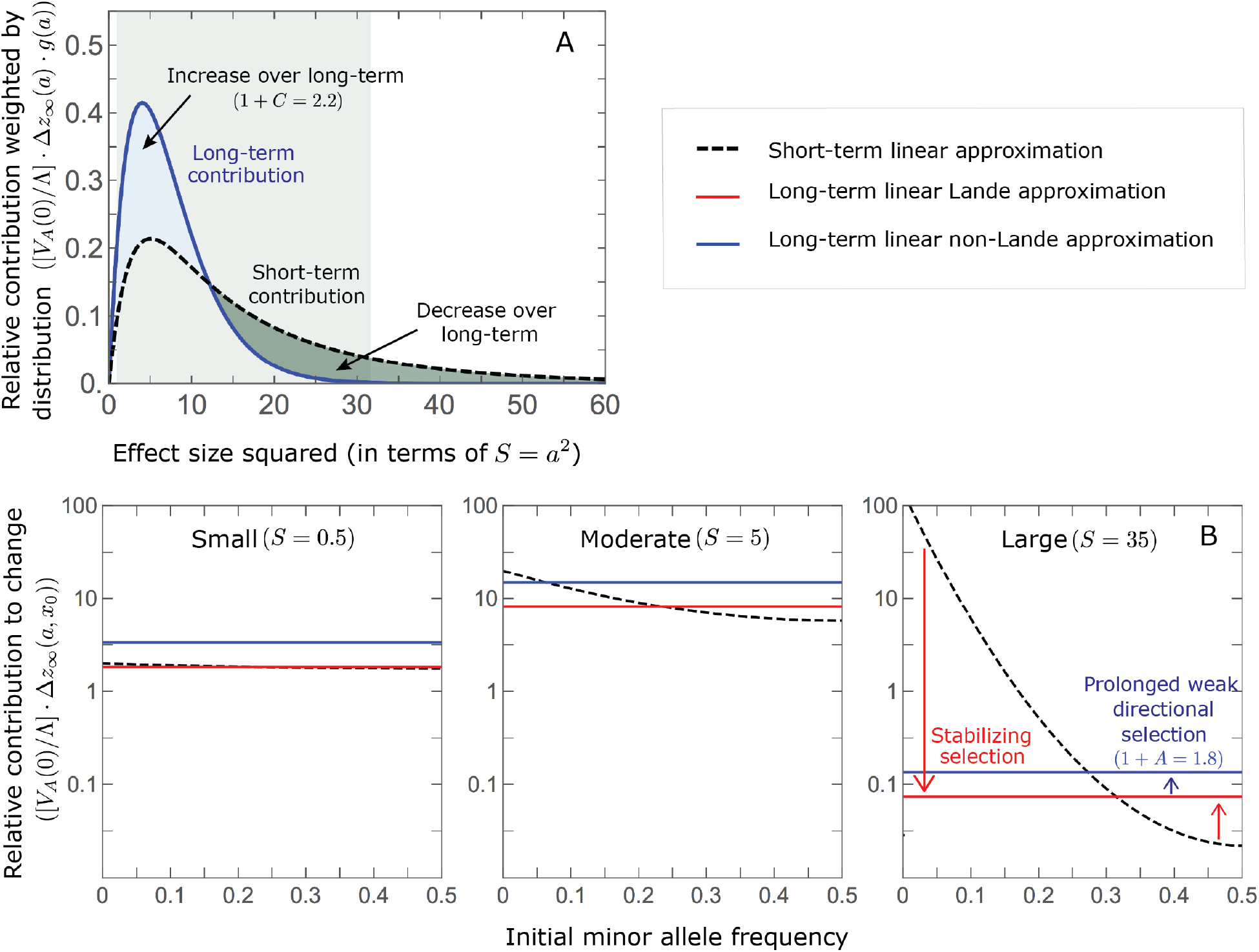
The genetic basis of adaptation turns over during the equilibration phase. A) The short-term contribution of large effect alleles is supplanted by the contribution of moderate effect alleles. As an illustration, we show the results of the linear approximation for the non-Lande case in ***Fig***. 5B (***Eqs. 13*** and ***28***). Specifically, we weight the short- and long-term relative contributions by the mutational input and use a linear (rather than log) scale for the squared effect sizes. This way we can see that the decrease in the contribution of large effect alleles (shaded dark gray area) equals the increase in the contribution of moderate effect alleles (shaded blue area). B) The proportional long-term contribution of alleles that segregated at low MAFs before the shift is diminished relative to their short-term contribution, an effect most pronounced for large effect sizes. As an illustration, we show the linear approximations for the contribution of alleles with a given effect size as a function of their initial MAF (***Eqs. 12, 19*** and ***24***) for the same non-Lande case as in A, with a shift of 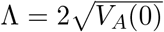 To this end, we estimate the amplification factor for standing variation (1 + *A*) from the allele simulations described in the legend of ***Fig. 5*** (see *SI Section 4.2*). In both the Lande and non-Lande cases, long-term stabilizing selection diminishes the contribution of alleles with lower initial MAF and amplifies the contribution of alleles with higher initial MAF (red arrow). In the non-Lande case, prolonged, weak directional selection amplifies the contribution of alleles, regardless of their initial MAF (blue arrow).

We can understand this turnover by considering the effects of stabilizing selection during the equilibration phase (***App. B Fig. 2***). As noted, stabilizing selection on the trait induces selection against minor alleles, which weakens as MAF increases and vanishes at MAF = 1/2. Now consider how it affects a pair of alleles with opposite, moderate or large effect. If their initial frequencies are very low, both alleles will have low MAFs at the end of the rapid phase. Consequently, they will both be strongly selected against during the equilibration phase and will almost certainly go extinct (***App. B Fig. 2***). In the long-run, their expected contribution to phenotypic adaptation is therefore diminished. In contrast, if the alleles’ initial MAF is sufficiently high, the relative increase in the aligned allele’s frequency by the end of the rapid phase causes it to be subject to substantially weaker selection than is the opposing allele. In the extreme in which the aligned allele has exceeded frequency 1/2, the direction of selection on it is even reversed (***App. B Fig. 2***). In such cases, the pair’s expected contribution to phenotypic adaptation will be amplified. This reasoning suggests that, for a given effect size, there is a critical initial MAF *<* 1/2 such that the long-term contribution of alleles that start above it is amplified and the contribution of those that start below it is diminished (***Figs. 6B*** and ***App. B Fig. 3***).

The turnover among alleles with different effect sizes can be explained in similar terms. Alleles with large effect sizes almost always start from low MAF, because they are subject to strong stabilizing selection before the shift (***Fig. 4B***). Consequently, they are highly unlikely to exceed the critical initial frequency and their expected long-term contribution to phenotypic adaptation is diminished (***Figs. 5A*** and ***6A***). However, changes to the frequencies of these alleles skew the phenotypic distribution, leading to prolonged, weak directional selection that amplifies the long-term contribution of small and moderate effect alleles (***Fig. 6B***). In our linear approximation, this amplification will occur for effect sizes that satisfy (1 + *C*) *f*(*a*) ⪆ *v*(*a*) (***Fig. 5B***). These considerations explain why the contributions of alleles with small and moderate effects supplant those of alleles with large effects (***Figs. 5A*** and ***6A***). They also highlight that deviations from Lande’s approximation are critical to understanding the allelic response, even when they have small phenotypic effects.

### Other properties of the equilibration process

While long-term phenotypic adaptation arises from an excess in fixations of aligned relative to opposing alleles, this excess and the increase in the total number of fixations are typically small relative to the number of fixations at steady-state (***Fig. 7***). The proportional excess of both aligned fixations and total fixations decrease with increased polygenicity and increase with the shift in optimum and allele effect size (***Fig. 7B*** and ***C***). For sufficiently large effect sizes, practically all fixations are caused by the shift and are of aligned alleles (***Fig. 7B*** and ***C***). However, with the exception of extreme cases in which the contribution of alleles of small and moderate effects to genetic variance is negligible, the number of fixations of such large effect alleles and their contribution to phenotypic change will be small (***Figs. 5*** and ***7A***). Typically, most fixations and contribution to phenotypic change will arise from alleles with small and moderate effects (***Figs. 5*** and ***7A***), for which the proportional excess of aligned and total fixations is modest (***Fig. 7B*** and ***C***).

**Figure 7.**
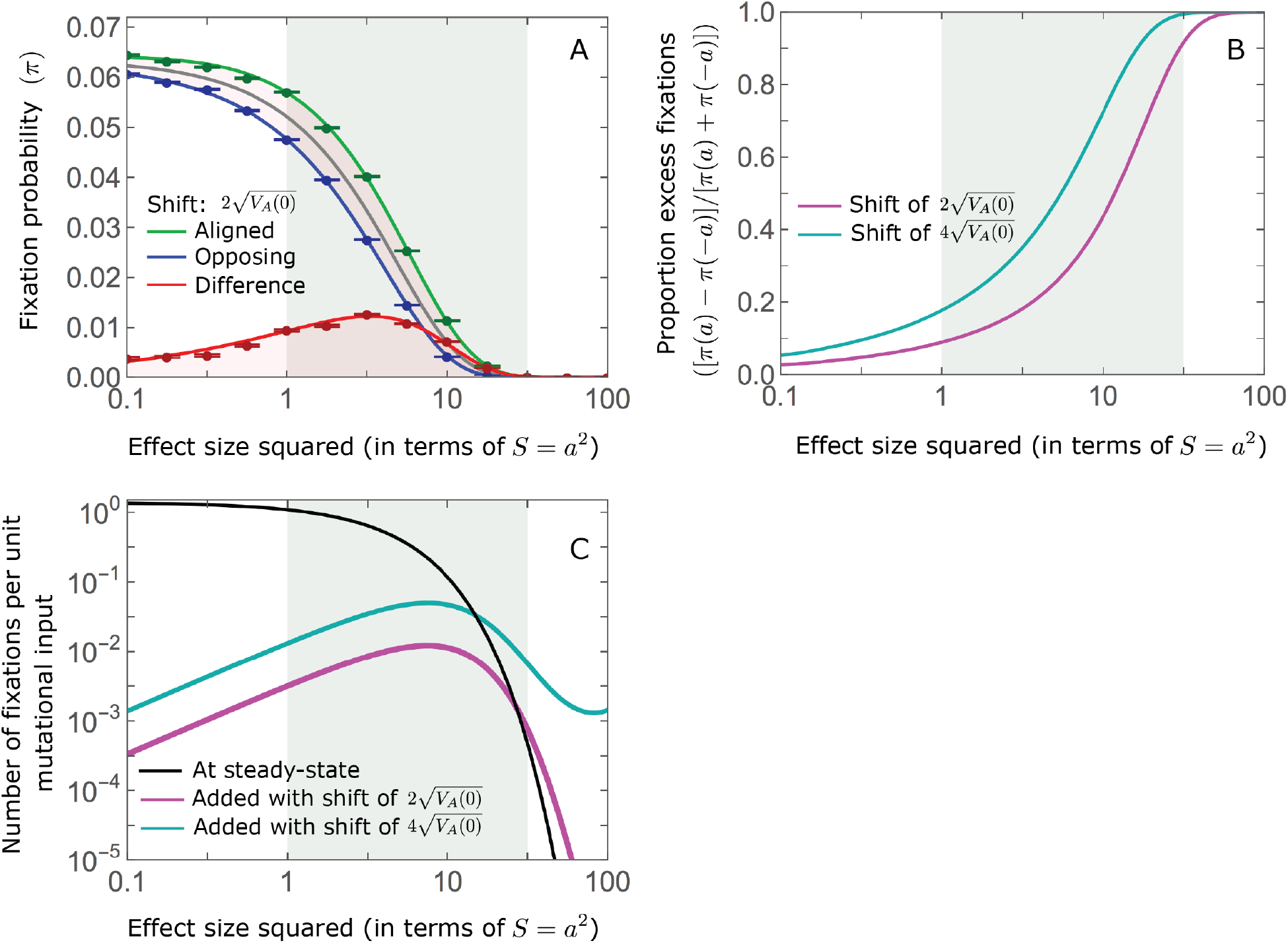
While long-term phenotypic adaptation arises from an excess in fixations of aligned relative to opposing alleles, this excess and its effect on the total number of fixations is typically small. A) The fixation probabilities of aligned and opposing alleles as function of their effect size. For large effect sizes, the fixation probabilities become vanishingly small, whereas for small and moderate effect sizes, the difference in the fixation probabilities of alleles with opposing effects is small. B) The proportional excess of fixations of aligned alleles as a function of effect size. C) Polygenic adaptation typically adds a small number of fixations relative to the number at steady state. For large effect sizes, the proportional increase in number is large, but their absolute number and the corresponding contribution to phenotypic adaptation are extremely small.

In the long run, these fixations move the mean phenotype all the way to the new optimum, and genetic variation around the new optimum returns to steady-state. A proxy for the approach to steady-state is the ‘fixed distance from the optimum’, defined as the phenotypic distance of an individual that is homozygote for the ancestral allele at every segregating site; at steady-state, we expect the fixed distance to be 0. Our simulations suggest that, under a broad set of parameter values, the change in fixed distance after the shift is well approximated by an exponential decay with a rate of 1/(2N) generations (***Figs. 8*** and ***S7.4***). This approximation is remarkably accurate in the Lande case. In the non-Lande case, the decay is initially slower than the approximation suggests, possibly because the long-term contribution of new mutations (as opposed to standing variation) takes longer to amass. In both cases, the return to steady-state occurs on a time scale of 2*N* generations after the shift (***Figs. 8*** and ***S7.4***).

**Figure 8.**
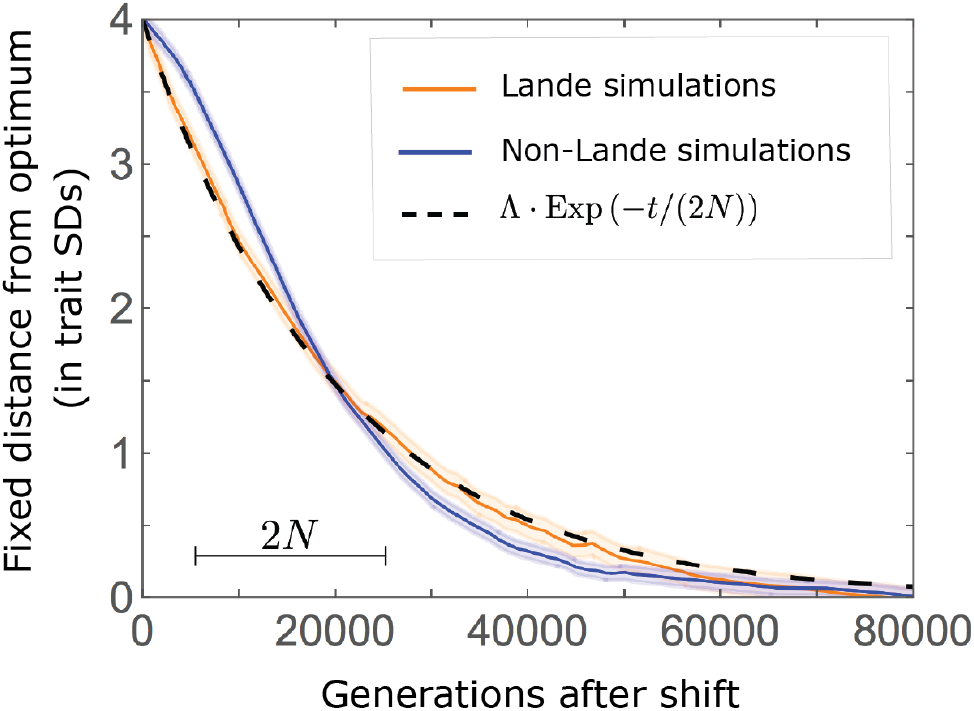
A steady-state around the new optimum is restored on a time-scale of 2*N* generations after the shift. The change in fixed distance after the shift estimated using simulations is compared with an exponential decay with a rate of 1/(2*N*) generations. Model parameters and other details of the simulations are the same in ***Fig. 2***. See ***Fig. S7.4*** for similar comparisons using a broad range of model parameter values.

## Discussion

Here, we investigated the phenotypic and genetic adaptive response to selection on a highly polygenic quantitative trait in a simple yet highly relevant setting, in which a sudden change in environment shifts the trait’s optimal value. The phenotypic response to selection was previously studied by Lande (1976). Assuming that phenotypes are normally distributed in the population, he predicted that after the shift the population’s mean phenotype will approach the new optimum exponentially, at a rate that is proportional to the additive genetic variance in the trait. We found that this prediction is accurate when the trait is sufficiently polygenic, the shift in optimum is not too large (relative to genetic variance in the trait), and the variance is dominated by loci with small and moderate effect sizes, which are defined based on the selection acting on them before the shift. When these conditions are violated, most notably when loci with large effects contribute markedly to genetic variance, the initial, rapid change in mean phenotype is followed by a pronounced quasi-static phase, governed by changes to the 3^rd^ moment of the phenotypic distribution, in which the mean phenotype takes much longer to catch up to the new optimum.

We also characterized the genetic basis of these adaptive phenotypic changes. The closest previous work assumed an infinite population size (de Vladar and Barton, 2014; Jain and Stephan, 2015, 2017a,b), and we found that relaxing this assumption leads to entirely different behavior. Notably, in infinite populations, small effect alleles whose frequencies before the shift are dominated by mutation and equal 1/2 make the greatest contribution to phenotypic change after the shift (see *introduction*). In contrast, in any real (finite) population, the frequencies of such small effect alleles are dominated by genetic drift rather than mutation. More generally, variation in allele frequencies due to genetic drift, which is absent in infinite populations, critically affects the allelic response to selection.

To study the allelic response, we divided it into two periods: a rapid phase, immediately after the shift, and a subsequent, prolonged equilibration phase. During the rapid phase, the population’s mean distance to the optimum is substantial and changes rapidly. Directional selection on the trait increases the frequency of minor alleles whose effects are aligned with the shift relative to minor alleles with opposing effects (given the same effect size and initial frequency). By the end of the rapid phase, the cumulative effect of these frequency differences pushes the mean phenotype close to the new optimum, but because this effect is spread over myriad alleles, the frequency difference between any individual pair of opposing alleles is fairly small. Specifically, we found that an allele’s contribution to phenotypic change is proportional to its contribution to phenotypic variance before the shift, implying that alleles with moderate and large effect sizes make the greatest per site contributions to phenotypic change, while alleles with moderate effect sizes experience the greatest frequency changes. The expected frequency differences between opposing alleles is amplified by prolonged, weak directional selection during the subsequent equilibration phase, and this amplification is pronounced when the phenotypic approach to the new optimum deviates markedly from Lande’s approximation.

Over the long run, stabilizing selection on the trait and genetic drift transform these small frequency differences into a small excess of fixed aligned alleles relative to opposing ones, and cumulatively this excess moves the population mean all the way to the new optimum. This transformation process involves a massive turnover in the properties of the contributing alleles. Notably, the transient contributions of large effect alleles are supplanted by contributions of fixed moderate, and to a lesser extent, small effect alleles. In the non-Lande cases, the fixation of mutations that arise after the shift in optimum can also contribute substantially to long-term phenotypic adaptation. These processes take on the order of 2*N*_*e*_ generations, after which the steady-state architecture of genetic variation around the new optimum is restored.

Our finding that large effect alleles almost never sweep to fixation appears at odds with the results of previous studies of similar models. These discrepancies are largely explained by earlier papers considering settings that violate our assumptions, notably about evolutionary parameter ranges. For instance, some studies assume that large effect alleles segregate at high frequencies before the shift in optimum (e.g., Christodoulaki et al., 2019), which is presumably uncommon in natural populations and in any case, violates our assumption that the trait is at steady-state before the shift. Other models implicitly consider quantitative traits of intermediate genetic complexity; while such traits likely exist, there are to our knowledge few well-established examples. Notably, Thornton (2019) observes sweeps in cases in which the trait is not highly polygenic (violating our assumption that 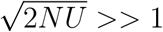). Relatedly, Chevin and Hospital (2008) observe sweeps in cases in which a single newly arising mutation of large effect contributes substantially to genetic variance, which violates our assumptions that genetic variation is highly polygenic and is not predominantly effectively neutral (i.e., that alleles with *S* ⪆ 1 contribute substantially). Although it remains to be seen, we believe that this architecture is much less common, given mounting evidence, reviewed in the *introduction*, which suggests that traits are often highly polygenic, and other considerations, notably estimates of persistence time (Walsh and Lynch, 2018; Sella and Barton, 2019) and inferences based on human GWASs (Simons et al., 2018; Zeng et al., 2018), which indicate that quantitative genetic variation is not predominantly neutral.

Lastly, Stetter et al. (2018) considered a huge shift in the optimal trait value (e.g., of ~90 phenotypic standard deviations), resulting in a massive drop in fitness (violating our assumption that 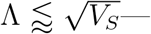 although shifts in optimum need not be that large to result in the fixations of some large effect alleles. While there are many examples of rapid and large environmental fluctuations, e.g., seasonal fluctuations or shifting weather systems, they occur on a much shorter time scale than fixation (although they might have some effect on genetic architecture; see below). In turn, little is known about the magnitude of shifts in optimal trait values over the time scales of large effect, beneficial fixations. While it seems plausible that moderate shifts, which fall within our assumed parameter ranges, are common, we cannot rule out that larger shifts are common as well. The response to such larger shifts is not covered by our analysis and clearly warrants further study.

Other factors that we have not considered may also affect polygenic adaptation. Most notable among them is pleiotropy. Given that quantitative genetic variation affecting one trait often affects many other traits (Bulik-Sullivan et al., 2015; Pickrell et al., 2016; Boyle et al., 2017; Sella and Barton, 2019; Liu et al., 2019), alleles that would have been positively selected because of their effect on the trait under directional selection may be selected against because of their adverse effects on other traits. Moreover, pleiotropy is known to affect the genetic architecture of a given trait at steady-state (Simons et al., 2018), which we have shown to shape the allelic response to selection on that trait. Pleiotropy is therefore likely to affect which alleles contribute to phenotypic change at the different phases of polygenic adaptation (see Otto, 2004, for related considerations for simple traits). Linkage disequilibrium (LD) may have an effect as well, perhaps most notably for minor alleles with large effects, which start at low frequencies and experience strong directional selection during the rapid phase. Before the shift, large effect alleles located in genomic regions with low recombination and high functional density are more likely to be in LD with, for example, alleles with countervailing effects on the focal trait (Lande, 1975) or deleterious effects on other traits. If this were the case, then directional selection during the rapid phase might be effectively weaker, because it would act on extended haplotypes rather than on individual alleles.

In addition, the demography of a population, notably its size, as well as the selection pressures on quantitative traits are likely to change over a shorter time scale than it takes the genetic architecture of complex traits to equilibrate. When these changes occur over the ~2*N*_*e*_ generations preceding a shift in optimal trait value, they could affect the genetic architecture of the trait and consequently its response to selection. Changes in population size influence the number of segregating sites affecting a trait and the distribution of their frequencies and contributions to variance, with more recent population sizes affecting strongly selected variation more than weakly selected variation (Lohmueller, 2014; Simons et al., 2014; Simons and Sella, 2016; Sella and Barton, 2019). The effects of varying selection will depend on the attributes of this variation in ways that await further study.

While the effects of all of these factors on the response to a shift in optimum warrant investigation, we expect the response to follow from the principles we outlined. Notably, we expect the short-term contribution of alleles to phenotypic change to be proportional to their contribution to variance before the shift, and their long-term contribution to arise from differences between the fixation probabilities of alleles with opposite effects, caused by the opposing effects of directional selection on their frequencies. Thus, while all of these factors are likely to affect the response, we expect the main features of the dynamics we portrayed to remain largely intact. These features include the role of the 3^rd^ moment of the phenotypic distribution in slowing down phenotypic adaptation near the new optimum; the transient contribution of large effect alleles to phenotypic adaptation; and the long-term importance of alleles with moderate effects.

As polygenic adaptation in quantitative traits is likely ubiquitous, our conclusions have potentially important implications. One is that, contrary to adaptation mediated by selective sweeps of initially rare, large effect, beneficial alleles (Smith and Haigh, 1974; Kaplan et al., 1989; Braverman et al., 1995; Hermisson and Pennings, 2005; Coop and Ralph, 2012; Berg and Coop, 2015), polygenic adaptation might have minor effects on patterns of neutral diversity at any given point in time (but may affect temporal diversity patterns (Buffalo and Coop, 2019, 2020). The effects of selected alleles on neutral diversity at linked loci follow from their trajectories (Barton, 2000). Our results indicate that directional selection on a highly polygenic trait introduces only small changes to allele frequencies at individual loci, which amount to minor perturbations to the allele trajectories expected under stabilizing selection at steady-state (also see Chevin and Hospital, 2008; Thornton, 2019). Indeed, alleles with large effects exhibit only small, transient changes. For those with more moderate effects, there is a modest, long-term excess of fixations of those alleles whose effects are aligned with the shift relative to those whose effects are opposed, accompanied by a small increase in the total number of fixations(***Fig. 7***). The trajectories of the alleles that fix are largely driven by weak stabilizing selection and tend to be drawn out (***Fig. 8***). Thus, our results indicate that the effects of polygenic adaptation on neutral diversity should be minor (other than perhaps for massive shifts in optimal trait values, as noted above).

In contrast, long-term stabilizing selection on quantitative traits likely has substantial effects on neutral diversity patterns. Specifically, selection against minor alleles induced by stabilizing selection may well be a major source of background selection and is expected to affect neutral diversity patterns in ways that are similar to those of background (purifying) selection from other selective origins (Charlesworth et al., 1993; Hudson and Kaplan, 1995; McVean and Charlesworth, 2000).

Another implication of our results pertains to the search for the genetic basis of human adaptation, as well as adaptation in other species. Efforts to uncover the identity of individual adaptive genetic changes on the human lineage were guided by the notion that their identity would offer insight into what “made us human”. Under the plausible assumption that many adaptive changes on the human lineage arose from selection on complex, quantitative traits, this approach may not be as informative as it appears (Pritchard et al., 2010; Boyle et al., 2017). Our results indicate that after a shift in the optimal trait value, the number of fixations of alleles whose effects are aligned to the shift are typically nearly equal to the number of alleles that are opposed (***Fig. 7A***). Moreover, the alleles that fix are a largely random draw from the vastly greater number of alleles that affect the trait, both in the sense of being those that happened to segregate at high MAFs at the onset of selection and because of the stochasticity of fixation. Thus, in this plausible scenario, it becomes meaningless to say that any given fixation was adaptive, and arguably uninteresting to focus on the particular subset of alleles that happened to reach fixation. In contrast, identifying the traits that experienced adaptive changes promises to provide important insights. Recent efforts to do so pool the signatures of frequency changes over many loci that were found to be associated with a given trait in GWAS (Turchin et al., 2012; Berg and Coop, 2014; Robinson et al., 2015; Field et al., 2016; Berg et al., 2017; Edge and Coop, 2019; Speidel et al., 2019), an exciting approach that has also proven to be technically challenging (Berg et al., 2019; Sohail et al., 2019). A better understanding of the process of polygenic adaptation should help to guide such efforts.

## Acknowledgments

We thank Guy Amster, Jeremy Berg, Nick Barton, Yuval Simons and Molly Przeworski for many helpful discussions, and Jeremy Berg, Joachim Hermisson, Will Milligan, Yuval Simons, Leo Speidel and Molly Przeworski for comments on the manuscript. This work was funded by NIH grant GM115889 to GS and NIH grant GM121372 to Molly Przeworski.

## A. Additional Tables

## B. Additional Figures

**Figure 1a.**
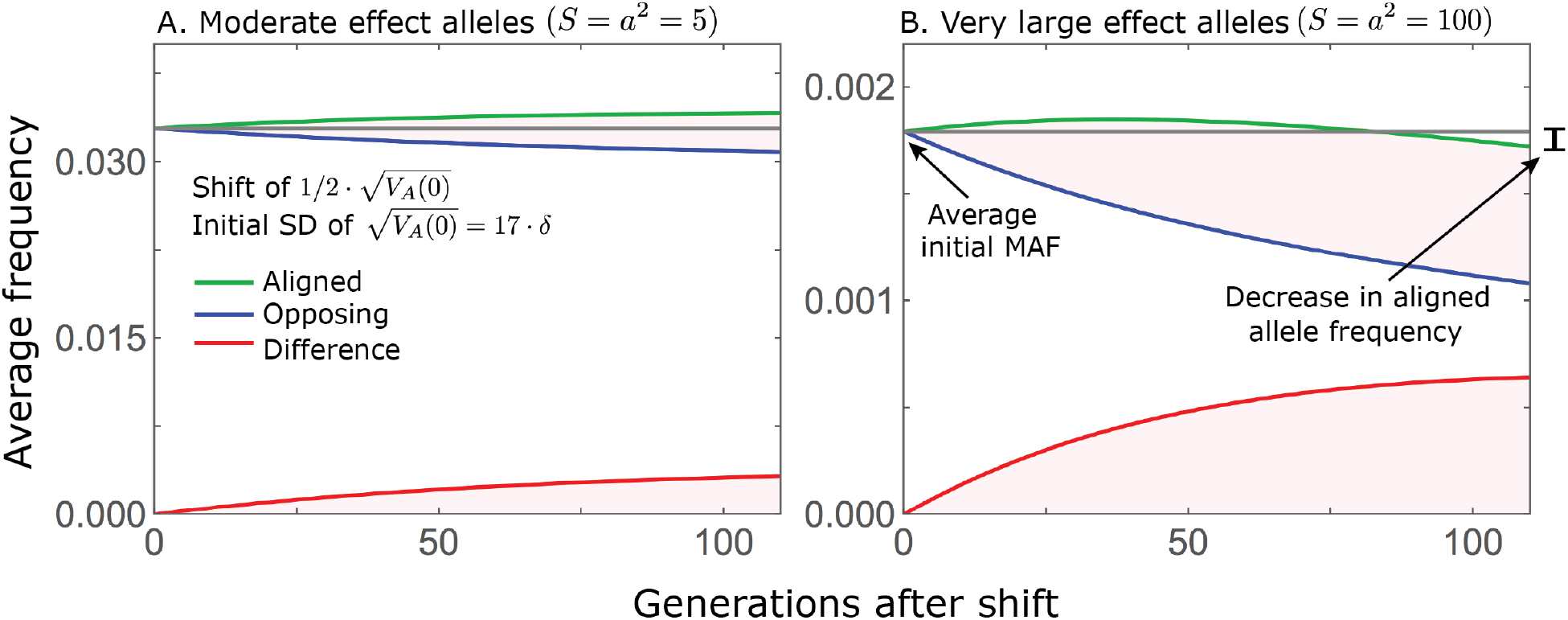
While directional selection during the rapid phase increases the frequency of aligned alleles relative to opposing ones, the frequency of aligned alleles does not necessarily increase. Here we show an example of the trajectories of (A) moderate and (B) large effect alleles in response to a relatively small shift in optimum; the trajectories were calculated using ***Eqs. S3.5***, ***S3.6*** and ***S3.14***. When directional selection is sufficiently weak (the shift is small), the frequency of aligned alleles with sufficiently large effects will decrease (B). However, the frequency of opposing alleles decreases *more*, and the frequency difference (in red) contributes to the change in mean phenotype.

**Figure 2a.**
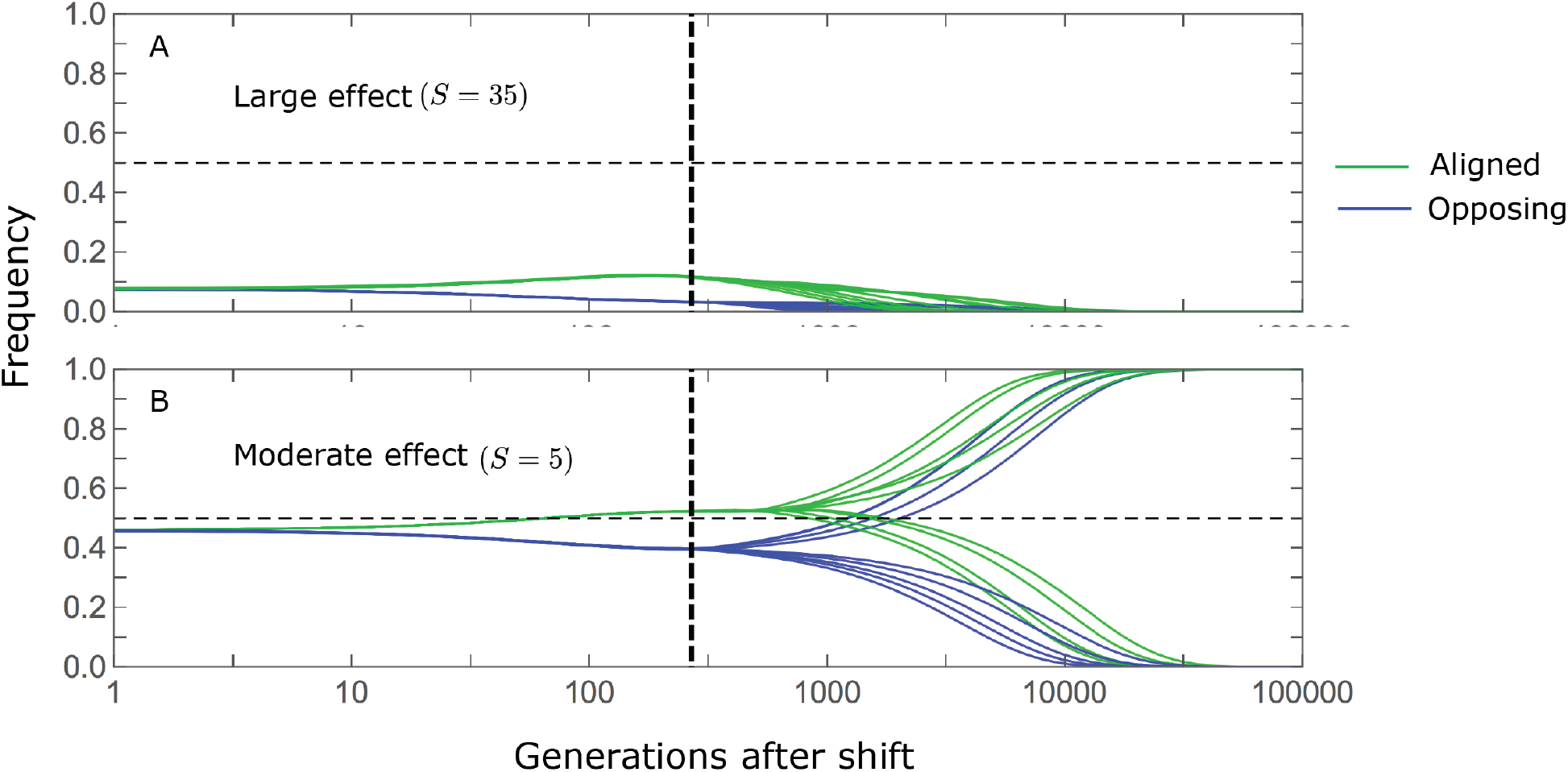
Stabilizing selection during the equilibration phase causes turnover in the genetic basis of adaptation. The cartoons depict the trajectories of alleles with opposing effects of a given magnitude, and a given initial MAF. For the purpose of illustration, we focus on alleles with large (A) and moderate (B) effects, with initial MAFs in the tail of the corresponding steady state MAF distribution (the 99.5^th^ percentile), and a shift size of 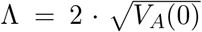with 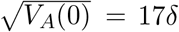. Directional selection during the rapid phase increases the frequency of aligned alleles relative to those with opposing effects, and these frequency differences underlie short-term phenotypic adaptation. A) The initial MAF of large effect alleles, even those in the 99.5^th^ percentile, is sufficiently low such that both aligned and opposing alleles still have low MAFs at the end of the rapid phase. Consequently, they are both strongly selected against during the equilibration phase and almost certainly go extinct, thereby erasing their short-term contribution to phenotypic adaptation. B) Moderate effect alleles start at much higher initial MAFs. In the extreme, this initial frequency is sufficiently high for directional selection during the rapid phase to push aligned alleles above frequency 1/2, thereby reversing the direction of (under-dominant) selection on them, but not on the opposing alleles, during the equilibration phase. Consequently, the expected contribution of moderate effect alleles with sufficiently high initial MAF to phenotypic adaptation is amplified during the equilibration phase.

**Figure 3a.**
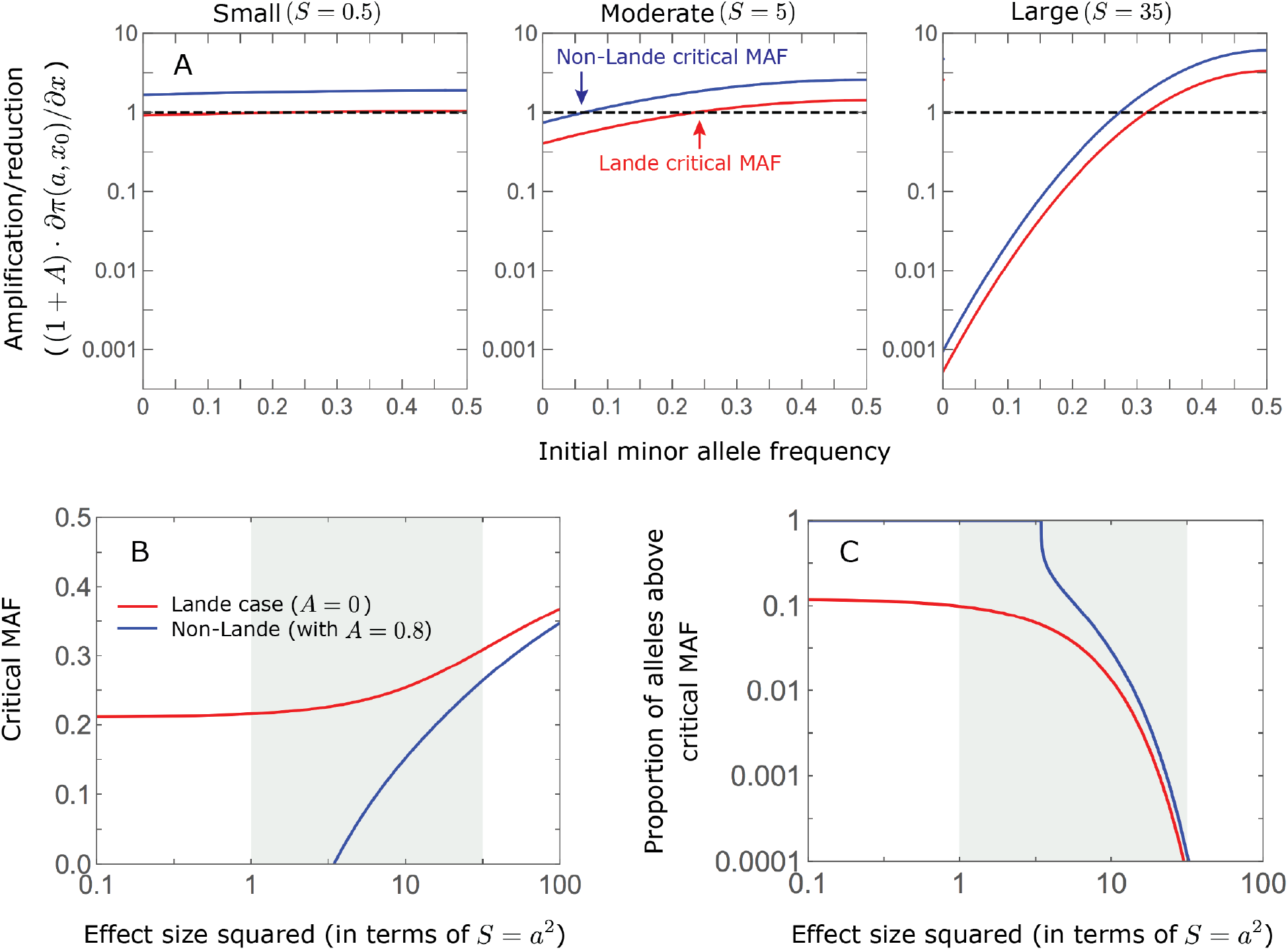
The long-term phenotypic contribution of alleles is amplified if they start above a critical initial MAF, which depends on their effect size, and it is diminished if they start below this critical MAF. In the instantaneous pulse approximation, we found that

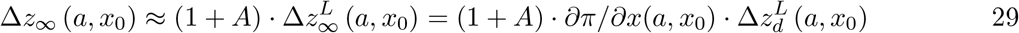 (***Eq. 24***). Further assuming that 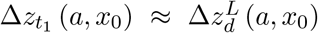 (which holds if the shift is not miniscule, i.e., Λ ≫ *δ*), we find that the (multiplicative) amplification/reduction of the phenotypic contribution is approximated by (1 + *A*) *∂π/∂x*(*a, x*_0_). In this approximation, the critical MAF for alleles with effect size *a* satisfies

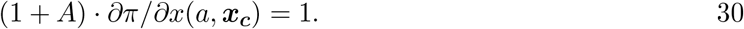 A) The amplification/reduction for alleles with small (left), moderate (middle), and large (right) effects as a function of initial MAF, in the Lande (red) and a non-Lande (blue) case. The curves and critical MAFs are calculated from ***Eqs. 29*** and ***30***, respectively. Given a factor *A >* 0, the contribution of alleles with sufficiently small effect sizes are amplified for any initial MAF *x*_0_ (***Fig. 6***), because *∂π/∂x*(*a, x*_0_) ≈ 1 and thus 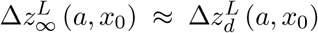 and 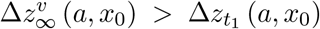 (***Eq. 29***). In turn, for sufficiently large effect sizes, the curves for 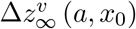 and 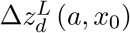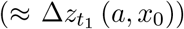 intersect (***Fig. 6B***) and thus a critical MAF exists (i.e., 0 *< x*_*c*_ *<* 1/2). These considerations explain why, for sufficiently large effect sizes, the long-term contribution of alleles with low initial MAFs is diminished relative to their short-term contribution. B) The critical MAF as a function of effect size in the Lande and a non-Lande case (based on ***Eq. 30***). The critical MAF is lower in the non-Lande case, because *A >* 0 (see ***Eq. 30***). This proportion declined with increasing effect size both because initial MAFs at steady state decrease and because the critical MAF increases (panel B). It is greater in the non-Lande case because the critical MAF is lower (panel B).

