## Supplementary Information for "Polygenic adaptation after a sudden change in environment"

### Polygenic adaptation after a sudden change in environment
### Supplementary Information

**Laura K. Hayward**[a,1] **and Guy Sella**[a,b,c,1]

[a]Department of Biological Sciences, Columbia University, New York, NY 10027
[b]Center for Computational Biology and Bioinformatics, Columbia University, New York, NY 10032
[c]Program for Mathematical Genomics, Columbia University, New York, NY 10032

### 1. The basic equations

Our analysis relies on two basic equations:

- **The allelic equation:** describing the expected change in frequency per generation of an allele with effect size $a$ and frequency $x$ (***Eq. 7*** in the main text):

$$E(\Delta x) \approx (a \cdot D(t)/V_S) \cdot x(1-x) - \left(a^2/V_S\right) \cdot (1 - D^2(t)/V_S) \cdot x(1-x)(1/2-x). \qquad \text{S1.1}$$

- **The phenotypic equation:** describing the expected change in the mean distance from the optimum, $D(t)$, per generation (***Eq. 3*** in the main text):

$$E(\Delta D(t)) \approx -(V_A(t)/V_S) \cdot D(t) + (1 - D^2(t))/V_S \cdot \mu_3(t)/(2V_S), \qquad \text{S1.2}$$

  where $V_A(t)$ and $\mu_3(t)$ denote the $2^{\text{nd}}$ and $3^{\text{rd}}$ central moments of the phenotypic distribution.

Previous work relied on similar allelic and phenotypic equations. Assuming a fitness landscape with a single optimum, the expected change in allele frequency per generation has commonly been approximated by

$$E(\Delta x) \approx (a \cdot D/V_S) \cdot x(1-x) - \left(a^2/V_S\right) \cdot x(1-x)(1/2-x) \qquad \text{S1.3}$$

(Barton, 1986; Charlesworth, 2013; de Vladar and Barton, 2014). This equation was used to derive phenotypic equations similar to ours (see below and Barton and Turelli, 1987; Bürger, 1991). The allelic equation has been justified for a Gaussian fitness function, i.e., $W(z) = \text{Exp}\left(-z^2/(2V_S)\right)$ (our ***Eq. 2***), further assuming that: i) phenotypes are normally distributed in the population; ii) the width of the distribution is far smaller than the width of the fitness function (i.e. $V_A \ll V_S$); and iii) the mean distance from the optimum is very small (i.e., $D^2 \ll V_S$) (Barton and Turelli, 1987; Simons et al., 2018). As we show in the main text, the assumption that the phenotype distribution is normally distributed can be violated in ways that affect the course of adaptation qualitatively, giving rise to the non-Lande case. Also, requiring that the mean distance from the optimum is very small can be restrictive.

We derive our basic equations (in *Section 1.1* our allelic equation and in *Section 1.2* our phenotypic equation) without making the assumptions of a normal phenotypic distribution or that the distance to the optimum is very small. Similar to previous work, we do not provide a rigorous treatment of the errors introduced by our approximations (see Hayward (2020) for such treatment). We therefore validate our main results against simulations of the full model (*Section 5*).

#### 1.1. Derivation of the allelic equation

The expected change in frequency of an allele with phenotypic effect $a$ at frequency $x$ in a single generation derives from the standard form (Gillespie, 2004):

$$E(\Delta x) = E(x') - x = \frac{x^2 \cdot \bar{W}_2 + x(1-x) \cdot \bar{W}_1}{\bar{W}} - x, \qquad \text{S1.4}$$

where $\bar{W}_i$ denotes the average fitness of an individual with $i = 0, 1$ or 2 copies of the allele and

$$\bar{W} \equiv x^2 \cdot \bar{W}_2 + 2x(1-x) \cdot \bar{W}_1 + (1-x)^2 \cdot \bar{W}_0 \qquad \text{S1.5}$$

is the average fitness. The average fitness of individuals with a given genotype $i$ at the focal site, $\bar{W}_i$, depends on the total *background phenotypic contribution* of other sites. An individual with $i$ copies of the allele and background contribution $R$ has fitness

$$W_i(R) = W(R + i \cdot a). \qquad \text{S1.6}$$

where $W$ is our Gaussian fitness function (***Eq. 2***), and the average fitness of individuals with $i$ copies at the focal site is

$$\bar{W}_i = \int_{-\infty}^{\infty} W(R + i \cdot a) \cdot p(R) dR, \qquad \text{S1.7}$$

where $p(R)$ denotes the distribution of background contributions (in continuous form), which is independent of the genotype at the focal site assuming linkage equilibrium.

We apply two successive approximations to get from ***Eq. S1.4*** to our allelic equation. First, we approximate the background contribution by its mean. The mean background contribution is $\mu - 2ax$, where $\mu$ is the mean trait value in the population and $2ax$ is the mean contribution of the focal site. In this approximation, the mean fitness of individuals with $i$ copies of the allele at the focal site is

$$\bar{W}_i = W(\mu + a \cdot (i - 2x)). \qquad \text{S1.8}$$

Substituting these expressions into ***Eqs. S1.4*** and ***S1.5*** yields

$$E(\Delta x) = E(x') - x \approx \frac{x^2 \cdot W(\mu + 2a(1-x)) + x(1-x) \cdot W(\mu + a(1-2x))}{\bar{W}} - x, \qquad \text{S1.9}$$

with $\bar{W} = x^2 \cdot W(\mu + 2a(1-x)) + 2x(1-x) \cdot W(\mu + a(1-2x)) + (1-x)^2 \cdot W(\mu - 2ax)$. Neglecting variation in background contributions seems reasonable given our assumption that the phenotypic standard deviation is small relative to the width of the fitness function (i.e. $\sqrt{V_A} \ll \sqrt{V_S}$). We use this approximation in our all alleles (AA) simulations (see *the model* section in the main text and *Section 5*).

Second, we obtain our allelic equation (***Eq. S1.1***) from the 2nd order Taylor expansion of ***Eq. S1.9*** in $a/\sqrt{V_S}$ around $a/\sqrt{V_S} = 0$. Neglecting terms of order $\left(a/\sqrt{V_S}\right)^3$ and higher seems reasonable given our assumption that $a/\sqrt{V_S} \ll 1$. However, the 2nd order expansion of a Gaussian becomes inaccurate far from its peak. In our case, this manifests in having the $i^{\text{th}}$ term of the expansion include expressions of the form $\left(a/\sqrt{V_S}\right)^i \cdot \left(D/\sqrt{V_S}\right)^k$ for $k \leq i$, which may not be negligible when $|D| \gg \sqrt{V_S}$. We therefore also require that $|D| \lessapprox \sqrt{V_S}$ for our allelic equation to be accurate.

#### 1.2. Derivation of the phenotypic equation

Next, we rely on our allelic equation (***Eqs. 7*** and ***S1.1***) and on our conditions on parameters to derive our phenotypic equation. To this end, we rewrite the allelic equation in terms of the change in an allele's contribution to the mean phenotype per generation:

$$E(2a \cdot \Delta x) \approx (v^*(a,x)/V_S) \cdot D - \left(1 - D^2/V_S\right) \cdot (v_3^*(a,x)/(2V_S)), \qquad \text{S1.10}$$

where $v^*(a,x) = 2a^2x(1-x)$ and $v_3^*(a,x) = a^3x(1-x)(1-2x)$ are the allele's contributions to the 2nd and 3rd moments of the phenotypic distribution, respectively. Our phenotypic equation then follows from expressing the expected change in the mean distance to the optimum per generation as a sum of these allelic contributions:

$$\begin{aligned} E(\Delta D) &= -\sum_i E(2a_i \cdot \Delta x_i) \\ &\approx -\left(\sum_i v^*(a_i,x_i)/V_S\right) \cdot D + \left(1 - D^2/V_S\right) \cdot \sum_i v_3^*(a_i,x_i)/(2V_S) \\ &= -(V_A/V_S) \cdot D + \left(1 - D^2/V_S\right) \cdot \mu_3/(2V_S), \end{aligned} \qquad \text{S1.11}$$

where $V_A$ and $\mu_3$ are the 2nd and 3rd moments of the phenotype distribution, which equal the corresponding sum over alleles under the assumption of linkage equilibrium.

The terms in the phenotypic equation can be interpreted as follows. The first term captures the effect of directional selection driving the mean phenotype towards the optimum at a rate that is proportional to the additive genetic variance (Lande, 1976). The 2nd part of the second term, $\mu_3/(2V_S)$, captures the effect of stabilizing selection on an asymmetric (skewed) phenotypic distribution. Namely, when the mean is near the optimum, stabilizing selection pushes the mean phenotype in the direction opposite to the thicker tail of the phenotype distribution, because, given that fitness is quadratic near the optimum, reducing the distance of extreme phenotypes (more of which lie in the thicker tail) from the optimum increases mean fitness. The 1st part of second term, $(1 - D^2/V_S)$, reflects the fact that stabilizing selection, which tends to reduce phenotypic variance, becomes weaker when the phenotypic mean is far from the optimum (provided $|D| < \sqrt{V_S}$). (It does so because the magnitude of the second derivative of the Gaussian fitness function decreases as the distance from the optimum increases.) When $D^2 \ll V_S$, our phenotypic equation is well approximated by

$$E(\Delta D) \approx -(V_A/V_S) \cdot D + \mu_3/(2V_S), \qquad \text{S1.12}$$

which was derived previously under the rare alleles approximation (Barton and Turelli, 1987) and assuming a parabolic fitness function (Bürger, 1991).

#### 2. Allelic dynamics under stabilizing selection

Understanding how stabilizing selection on the trait affects allelic dynamics is crucial to characterizing the allelic response after a shift in the optimal trait value. Notably, the steady-state allelic dynamic under stabilizing selection shapes the genetic architecture of the trait (i.e., the joint distribution of phenotypic effect sizes and frequencies of alleles affecting the trait) prior to the shift and this genetic architecture shapes the short term allelic and phenotypic response to the shift. Also, after the short, initial response, when the mean phenotype is close to the new optimum, the longer-term allelic dynamics, which are out of equilibrium, are largely determined by the effects of stabilizing selection on allelic trajectories. Here we therefore describe the allelic dynamics under stabilizing selection, both at steady-state and out of equilibrium.

The analysis of the effects of stabilizing selection is greatly aided by the fact that it is stationary. In particular, the first two moments of change in the frequency of an allele with effect size $a$ and frequency $x_0$ in a single generation do not depend on time:

$$E(\Delta x_0) \approx -a^2/V_S \cdot x_0(1-x_0)(1/2-x_0)$$
$$V(\Delta x_0) \approx x_0(1-x_0)/(2N). \qquad \text{S2.1}$$

Given this time-independence, we can use the diffusion approximation to calculate the sojourn time: the density of time a derived allele spends at any given frequency $x_0'$ until fixation or loss, having been introduced at some initial frequency $p$ (Ewens, 2012). This sojourn time is

$$\tau(a, x_0'; p) = \begin{cases} 2Nh(a, x_0')\left[h_+(a,p)/h_+(a,x_0')\right] & 0 \le x_0' \le p \\ 2Nh(a, x_0')\left[h_-(a,p)/h_-(a,x_0')\right] & p < x_0' \le 1 \end{cases}, \qquad \text{S2.2}$$

where

$$\begin{aligned} h(a,x) &\equiv (\sqrt{\pi}/2a) \cdot \left(\mathrm{Exp}\left(a^2/4 \cdot (1-2x)^2\right) / \left[\mathrm{Erf}\left(\tfrac{a}{2}\right) x(1-x)\right]\right) \cdot h_-(a,x)h_+(a,x), \\ h_+(a,x) &\equiv \mathrm{Erf}\left(\tfrac{a}{2}\right) + \mathrm{Erf}\left(\tfrac{a}{2}(1-2x)\right), \\ h_-(a,x) &\equiv \mathrm{Erf}\left(\tfrac{a}{2}\right) - \mathrm{Erf}\left(\tfrac{a}{2}(1-2x)\right) \end{aligned} \qquad \text{S2.3}$$

and Erf is the Gaussian error function. We can also use the diffusion approximation to calculate the probability that an allele fixes or goes extinct, and the sojourn time conditional on these outcomes. The probability of fixation, $\pi$, or loss, $1-\pi$, are given by

$$\pi(a,x) = h_-(a,x)/\left(2\mathrm{Erf}\left(a/2\right)\right) \quad \text{and} \quad 1-\pi(a,x) = h_+(a,x)/(2\mathrm{Erf}\left(a/2\right)). \qquad \text{S2.4}$$

In our analysis of allelic behavior during the equilibration phase, we rely on the derivative of the fixation probability with respect to the initial frequency (***Eq. 18*** in the main text), which is

$$\frac{\partial \pi(a,x)}{\partial x} = \frac{2f(a)}{v(a,x)} \quad \text{where} \quad f(a) \equiv 2a^3 \cdot \frac{\mathrm{Exp}\left(-a^2/4\right)}{\sqrt{\pi} \cdot \mathrm{Erf}(a/2)}. \qquad \text{S2.5}$$

and $v(a,x) \equiv 4a^2\mathrm{Exp}\left(-a^2x(1-x)\right)$, which is the steady state density of variance per unit mutational input from segregating sites with effect size $a$ and minor allele frequency (MAF) $x$ (*Section 2.2*).

The sojourn time take a much simpler form when phrased in terms of the minor rather than derived allele frequency, because the strength of stabilizing selection depends on the former. For a new mutation ($p = 1/(2N)$), we call this the *folded sojourn time*, and the expression for it is

$$\begin{aligned} \tau_M(a, x_0) &\equiv \tau\left(a, x_0; 1/(2N)\right) + \tau\left(a, 1-x_0; 1/(2N)\right) \\ &\approx \begin{cases} (2Nx_0) \cdot 2 \cdot \mathrm{Exp}\left(-v^*(a,x_0)/2\right)/\left[x_0(1-x_0)\right] & 0 \le x_0 \le 1/(2N) \\ 2 \cdot \mathrm{Exp}\left(-v^*(a,x_0)/2\right)/\left[x_0(1-x_0)\right] & 1/(2N) < x_0 \le 1/2 \end{cases}, \end{aligned} \qquad \text{S2.6}$$

where $v^*(a, x_0) \equiv 2a^2x_0(1-x_0)$ is the genetic variance contributed by an allele with phenotypic effect $a$ and frequency $x_0$. As this equation shows, during a mutation's sojourn through the population, the expected time that the minor allele segregates at a particular frequency declines exponentially with its contribution to variance, $v^*(a, x_0)$. This accords with the intuition that stabilizing selection acts to reduce phenotypic variance.

##### 2.1. Summaries of architecture

Assuming that mutation is symmetric (i.e., with equal rates of positive and negative effects of any given magnitude), we can use the folded sojourn time to calculate summaries of the genetic architecture at steady-state. Notably, the steady-state density of sites segregating with effects of size $\pm a$ and MAF $x_0$ per unit input of mutations of that size is:

$$\rho\left(a,x_0\right)=\tau_M(a,x_0)/2 \tag{S2.7}$$

(and the number of segregating sites is $2NU\cdot g(a)\cdot\rho\left(a,x_0\right)$, where $2NU\cdot g(a)$ is the expected number of new mutations per generation with effect $a$). For brevity, henceforth we refer to such densities as being per unit mutational input, by which we mean mutations with a given magnitude. The exponential decline of $\rho\left(a,x_0\right)$ with $v^*\left(a,x_0\right)$ implies that at steady-state, alleles rarely segregate at MAFs much greater than $1/a^2$.

We can calculate the expectations of most summaries of interest at steady state using the density $\rho\left(a,x_0\right)$. Consider a summary $K$ that depends only on segregating sites, and to which a segregating site whose minor allele has effect size $a$ and frequency $x_0$ contributes $k^*\left(a,x_0\right)$. The density of contribution of such sites per unit mutational input is $k^*\left(a,x_0\right)\cdot\rho\left(a,x_0\right)$, and the corresponding density for sites with effects of magnitude $|a|$ is

$$k\left(a,x_0\right)=\left(k^*\left(a,x_0\right)+k^*\left(-a,x_0\right)\right)\cdot\rho(a,x_0). \tag{S2.8}$$

The marginal density for all sites with this magnitude (summing over MAFs) is

$$k\left(a\right)=\int_0^{1/2}k(a,x_0)dx_0, \tag{S2.9}$$

and the expected value of the summary, weighted by the input of mutations with any given effect size, $2NU\cdot g\left(a\right)$, is

$$K=2NU\cdot\int k\left(a\right)g\left(a\right)da. \tag{S2.10}$$

In some cases, we are interested in the expected contribution to a summary *per segregating site* rather than *per unit mutational input.* We can calculate such contributions by replacing the density $\rho\left(a,x_0\right)$ in ***Eqs. S2.8*** and ***S2.9*** by the proportional density of time that segregating sites with effect size $a$ spend with MAF $x_0$ (which is also the MAF distribution at steady state):

$$\rho\left(x_0|a\right)=\tau_M(a,x_0)/\int_0^{1/2}\tau_M(a,y)dy. \tag{S2.11}$$

The denominator of this expression is the expected time that a mutation with effect size $a$ segregates before fixation or loss.

##### 2.2. Phenotypic variance at steady state

One summary of genetic architecture at steady state that is central to our analysis is the distribution of phenotypic variance among alleles. An allele with effect size $a$ and MAF $x_0$ contributes

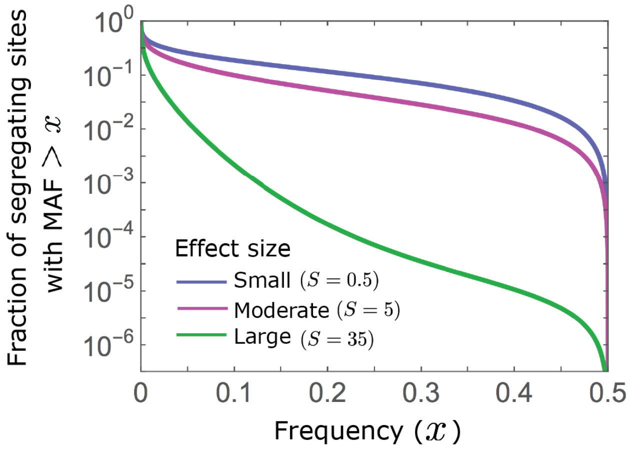

**Figure S2.1.** The allelic densities at steady-state. We show the proportion of segregating sites with MAF $> x$, as a function of $x$. Large effect minor alleles segregate at much lower frequencies than minor alleles with intermediate and small effects (see ***Eqs. S2.11*** and ***S2.6***).

$v^*(a, x_0) = 2a^2 x_0(1 - x_0)$ to phenotypic variance. The density of variance per unit mutational input arising from minor alleles with effects of magnitude $|a|$ and frequency $x_0$ is therefore

$$v(a, x_0) \approx \begin{cases} (2Nx_0)\, 4a^2 \cdot \text{Exp}\left(-v^*(a, x_0)/2\right) & 0 \le x_0 \le 1/(2N) \\ 4a^2 \cdot \text{Exp}\left(-v^*(a, x_0)/2\right) & 1/(2N) \le x_0 \le 1/2 \end{cases}, \quad \text{S2.12}$$

and the marginal density of alleles with a given effect size is

$$v(a) \equiv \int_0^{1/2} v(a, x_0)\, dx_0 = 4a \cdot D_+(a/2), \quad \text{S2.13}$$

where $D_+$ is the Dawson function $\left(D_+(y) \equiv \frac{\sqrt{\pi}}{2} \cdot \text{Exp}\left(y^2\right) \cdot \int_0^y \text{Exp}\left(-u^2\right) du\right)$.

These expressions clarify how the contributions of alleles to variance depend on their effect sizes (Simons et al. (2018) and ***Figs. 4A*** and ***S2.2***). From the expression for $v(a, x_0)$, we see that the bulk of variance from sites with large effect sizes ($S = a^2 \ll 1$) arises from rare minor alleles ($x_0 \lessapprox 1/a^2$), whereas the variance from sites with small effect sizes ($S = a^2 \ll 1$) is approximately uniformly distributed across MAFs (since the exponent $v^*(a, x_0) \ll 1$) (***Fig. 6B***). Other properties of $v(a)$ are detailed in the main text, as they shape the short-term allelic response to selection (section on *the allelic response in the rapid phase*, *Section 3* and ***Figs. 4***A and ***S2.2***).

The properties of $v(a)$ also allow us to translate our conditions on the shift size and phenotypic standard deviation (***App. A Table 2***) into conditions on basic parameters. Notably, the standard deviation of the phenotypic distribution at steady-state can be written as

$$\sqrt{V_A(0)} = \sqrt{2NU}\sqrt{\int_0^\infty v(a)\, g(a) da} \cdot \delta, \quad \text{S2.14}$$

where $\delta = \sqrt{V_S/(2N)}$ is the standard deviation of the distance between the mean and optimal phenotype at steady state (due to stochastic fluctuations). While we generally work in units in which $\delta = 1$, we have written $\delta$ explicitly here to emphasize the relationship between $\sqrt{V_A(0)}$ and $\delta$. Specifically, given our conditions that the trait is highly polygenic, i.e., that $\sqrt{2NU} \gg 1$, and that a substantial portion of mutations are not effectively neutral, with effect sizes such that $v(a) \gtrapprox 1$ (***Fig. S2.2***), ***Eq. S2.14*** implies that our assumption that $\sqrt{V_A(0)} \gg \delta$ holds. Given our condition

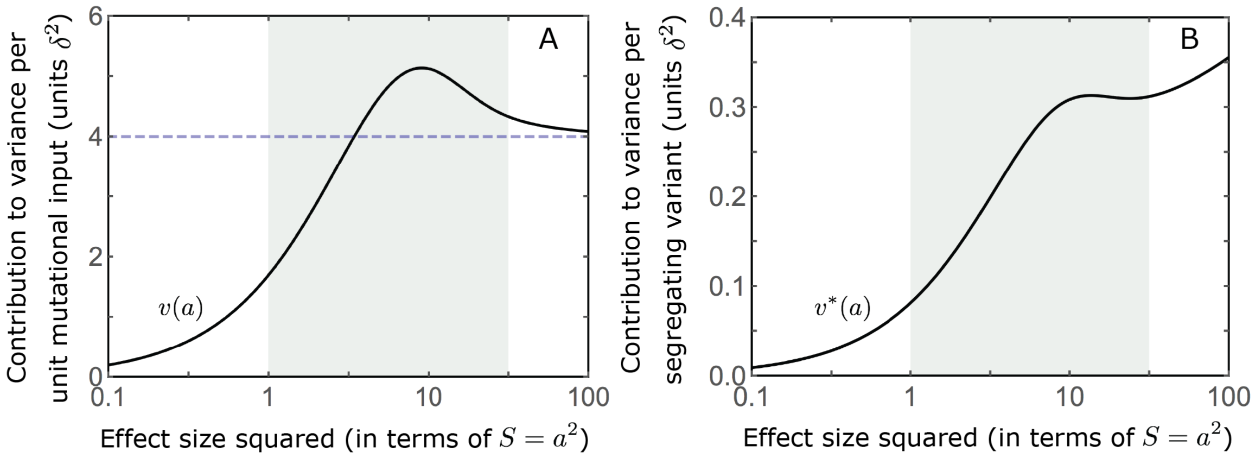

**Figure S2.2.** The distribution of variance at steady state. We show the contribution to variance per unit mutational input (A) and per segregating variant (B), as a function of effect size squared.

that the shift size $\Lambda \lessapprox \sqrt{V_S}$ (i.e., the shift does not drastically reduce mean fitness), ***Eq. S2.14*** also implies that

$$\frac{\Lambda}{\sqrt{V_A(0)}} \lessapprox \frac{1}{\sqrt{U}} \cdot \frac{1}{\sqrt{\int_0^\infty v\,(a)\,g(a)da}} \tag{S2.15}$$

(***Fig. S2.3***). The constraint on shift size is the strongest (i.e., the right-hand side is smallest) when most effect sizes are moderate and large. In this case, $v(a) \approx 4$ (***Eq. S2.13*** and ***Figs. 4A*** and ***S2.2***) implying that $\Lambda/\sqrt{V_A(0)} \lessapprox 1/(2\sqrt{U})$ and that the mutation rate $U$ constrains how large a shift can be relative to the phenotypic standard deviation (***Fig. S2.3***). However, when the mutation rate is low, specifically when $1/\sqrt{U} > \sqrt{2NU}$, this constraint is weak and our assumption that the shift is not massive relative to the initial phenotypic standard deviation i.e., that $\Lambda/\sqrt{V_A(0)} \lessapprox 1/2 \cdot \sqrt{2NU}$ (***App. A Table 2***), provides a tighter upper bound for the size of the shift.

Lastly, the properties of $v(a)$ allow us to derive bounds on the effect of directional selection on allele frequencies during the rapid phase. Specifically, the total change in allele frequency due to directional selection during the rapid phase is approximately proportional to $\int (D_L(\tau)/V_S)\, d\tau \approx \Lambda/V_A(0)$ (***Eq. 10***), whereas ***Eq. S2.15*** and our condition that the shift is not massive, i.e. $\Lambda/\sqrt{V_A(0)} \lessapprox 1/2 \cdot \sqrt{2NU}$ (***App. A Table 2***), and $v(a) \lessapprox 4$, imply that

$$\begin{aligned}\frac{\Lambda}{V_A(0)} &= \frac{\Lambda}{\sqrt{V_A(0)}} \cdot \frac{1}{\sqrt{V_A(0)}} = \frac{\Lambda}{\sqrt{V_A(0)}} \cdot \frac{1}{\sqrt{2NU}\,\sqrt{\int_0^\infty v\,(a)\,g(a)da}} \\ &\lessapprox \frac{1}{2\sqrt{\int_0^\infty v\,(a)\,g(a)da}} \lessapprox \frac{1}{2}.\end{aligned} \tag{S2.16}$$

As expected, a smaller shift size ($\Lambda/\sqrt{V_A(0)}$) and greater extent of polygenicity ($\sqrt{2NU}$) reduce the integral effect of directional selection on allele frequencies. Additionally, our conditions on parameters impose a bound on this effect.

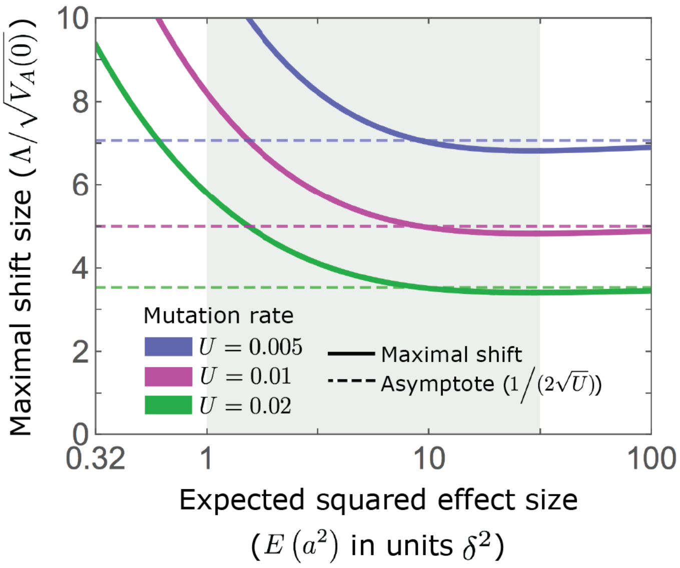

**Figure S2.3.** An upper bound on shift size measured in phenotypic standard deviations. As detailed in the text, our condition that $\Lambda \lessapprox \sqrt{V_S}$ sets an upper bound on $\Lambda/\sqrt{V_A(0)}$, which in turn depends on the mutation rate $U$ and the distribution of effect sizes (***Eq. S2.15***). As an illustration, we assume an exponential distribution of squared effect sizes, with the expectation $E(S) = E(a^2)$ shown on the $x$-axis; the $x$-axis starts at 0.32 because of our condition that a substantial portion of mutations are not effectively neutral (see ***App. A Table 2***).

#### 2.3. 3rd phenotypic moment

The 3rd moment of the phenotypic distribution also plays a central role in the response to selection. Notably, when alleles with large effects contribute markedly to genetic variance, the 3rd moment increases substantially from zero shortly after the shift in optimum, and the long-term phenotypic response takes what we refer to as the "non-Lande" form (see the *phenotypic response* section and ***Fig. 2C***). Here, we propose an explanation for why alleles with large effects lead to a substantial increase in the 3rd moment whereas alleles with small effects do not.

First consider the distribution of the allelic contributions to the 3rd moment at steady-state. An allele with effect size $a$ and MAF $x_0$ contributes $v_3^*(a, x_0) = 2a^3x_0(1-x_0)(1-2x_0)$ to the 3rd moment of the phenotype distribution. At steady-state, the contributions of alleles with opposing effects cancel out, because for each effect size and MAF the density is $v_3(a,x) = [v_3^*(a,x) + v_3^*(-a,x)] \cdot \rho(a,x) = [v_3^*(a,x) - v_3^*(a,x)] \cdot \rho(a,x) = 0$. To learn about the genetic architecture of the 3rd moment at steady-state, we therefore consider the contributions of alleles with positive and negative effect sizes separately. The density of the 3rd moment per unit mutational input of alleles with positive $(+)$ or negative $(-)$ effect sizes is

$$v_3^{\pm}(a, x_0) \equiv v_3^*(\pm a, x_0) \cdot \rho(a, x)$$
$$\approx \pm \begin{cases} (2Nx_0) \cdot 4a^2 \cdot \mathrm{Exp}\left(-v^*(a,x_0)/2\right) \cdot (1/2 - x_0) & 0 \le x_0 \le 1/(2N) \\ 4a^2 \cdot \mathrm{Exp}\left(-v^*(a,x_0)/2\right) \cdot (1/2 - x_0) & 1/(2N) < x_0 \le 1/2 \end{cases}, \quad \text{S2.17}$$

and the corresponding marginal density of alleles with a given effect size is

$$v_3^{\pm}(a) = \int_0^{1/2} v_3^{\pm}(a, x_0)\, dx_0 = \pm 2a \cdot \left(1 - \mathrm{Exp}\left(-a^2/4\right)\right) \quad \text{S2.18}$$

(***Fig. S2.4***). It follows that alleles with smaller effects ($S = a^2 \ll 4$) contribute negligibly to the 3rd moment (because $\mathrm{Exp}(-a^2/4) \approx 1$), whereas alleles with large effects ($S = a^2 \gg 4$) can have substantial contributions, which increase linearly with their effect size (because $\mathrm{Exp}(-a^2/4) \ll 1$).

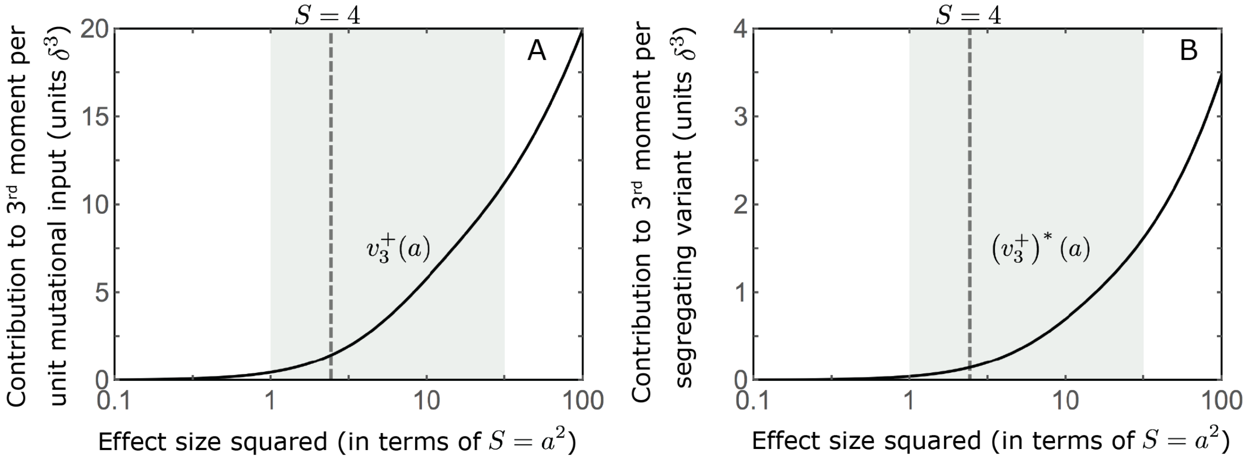

**Figure S2.4.** The distribution of the 3rd phenotypic moment at steady state. We show the positive/negative contribution to the 3rd moment per unit mutational input (A) (based on ***Eq. S2.18***) and per segregating variant (B), as a function of effect size squared. The contributions become substantial for large effect sizes, i.e., $S = a^2 \gg 4$.

After the shift, the frequency increase of aligned alleles relative to opposing ones introduces a non-zero 3rd (***Fig. 2C***). Large effect alleles contribute substantially more to this 3rd moment, plausibly because their 'hidden' steady-state contributions to the 3rd moment are substantially greater and because they exhibit large changes in frequency relative to their initial frequency before the shift.

#### 3. Allelic dynamics during the rapid phase

Immediately after the shift in optimum, the population exhibits a rapid phenotypic and allelic adaptive response. Phenotypically, the population mean rapidly approaches the new optimum, as described by Lande's approximation (***Eq. 5***). In the *allele dynamics* section, we define the end of the rapid phase as the time at which the Lande's approximation for the distance $D_L(t_1)$ first equals $\delta = \sqrt{V_s/(2N)}$ (***Eq. 9*** and ***Fig. S7.1***); while somewhat arbitrary, the precise definition of this time is not fundamental for our analysis. The rapid phenotypic response arises from rapid changes to allele frequencies. Here we derive two approximations for these frequency changes and the corresponding contributions of alleles to phenotypic change.

##### 3.1. Approximate solutions for allelic trajectories

Allelic trajectories during the rapid phase are nearly deterministic, because it is too short for genetic drift to have a substantial effect. Further, deviations from Lande's approximation for the distance of the mean phenotype have negligible effects (see ***Fig. S3.1D-F***). We can therefore approximate these trajectories based on our expression for the first moment of change in allele frequency (***Eq. S1.1***) and Lande's approximation for the mean phenotype (***Eq. 5***). Rewriting this expression in continuous time and in the standard semi-dominant form, we find that

$$\frac{dx}{dt} = \frac{s(t)}{2}x(1-x), \quad \text{S3.1}$$

with selection coefficient

$$s(t) = \pm 2\left(a \cdot D_L(t)/V_S\right) - a^2/V_S \cdot \left(1 - D_L^2(t)/V_S\right) \cdot (1-2x). \tag{S3.2}$$

The first term in the selection coefficient reflects directional selection and the second term reflects stabilizing selection. The selection coefficient varies with time, because the strengths of directional and stabilizing selection depend on the changing distance to the new optimum, $D(t)$. It also depends on frequency, because stabilizing selection is stronger when the MAF is lower.

We can derive an implicit solution for the ordinary differential equation (ODE) for allelic trajectories. Rewriting ***Eq. S3.2*** as $1/\left[x(1-x)\right] \cdot dx = s(t)/2 \cdot dt$ and integrating both sides, we find that

$$\Delta x_t(a,\ x_0) = x_t - x_0 = x_0(1-x_0)\left(\frac{\text{Exp}\left[\int_0^t \frac{s(\tau)}{2} d\tau\right] - 1}{1 + x_0\left(\text{Exp}\left[\int_0^t \frac{s(\tau)}{2} d\tau\right] - 1\right)}\right). \tag{S3.3}$$

In the standard semi-dominant case, with a constant selection coefficient, this expression with $\int_0^t s(\tau)\, d\tau = s \cdot t$ yields the standard explicit solution for allelic trajectories. In our case, the selection coefficient and thus $\int_0^t s(\tau)\, d\tau$ depends on $x_t$ making ***Eq. S3.3*** an implicit solution. Substituting our expression for the selection coefficient into this integral, we can express it as

$$\int_0^t \frac{s(\tau)}{2} d\tau = \left(\bar{s}_D(a,t) + \bar{s}_S(a,x_0,t)\right) \cdot t, \tag{S3.4}$$

where $\bar{s}_D$ and $\bar{s}_S$ denote the average directional and stabilizing selection coefficients respectively. In turn

$$\begin{aligned} \bar{s}_S(a,x_0,t) &= -\frac{a^2}{V_S} \cdot \overline{(1/2 - x)(1 - D_L^2/V_S)}, \\ \bar{s}_D(a,t) &= \frac{a \cdot \overline{D}_L(t)}{V_S} \text{ and } \overline{D}_L(t) = \Lambda/V_A(0) \cdot (1 - D_L(t)/\Lambda) \cdot V_S/t, \end{aligned} \tag{S3.5}$$

with all averages being taken from the time of the shift to time $t$.

In developing our two explicit approximations, we rewrite the implicit solution as

$$\Delta x_t(a,\ x_0) = x_t - x_0 = a x_0(1-x_0) \cdot F_t(a,x_0), \tag{S3.6}$$

where

$$F_t(a,x_0) \equiv \frac{1}{a} \cdot \left(\frac{\text{Exp}\left[\left(\bar{s}_D(a,t) + \bar{s}_S(a,x_0,t)\right) \cdot t\right] - 1}{1 + x_0 \cdot \left(\text{Exp}\left[\left(\bar{s}_D(a,t) + \bar{s}_S(a,x_0,t)\right) \cdot t\right] - 1\right)}\right). \tag{S3.7}$$

In the simple, *linear approximation* that we use in the main text, the effect of directional selection on $F_t(a,x_0)$ depends only on the sign of the effect size, and the effect of stabilizing selection depends only on the magnitude of the effect size; as a result, when we consider the frequency difference between matched pairs of opposing alleles below (see also the section on *the allelic response in the rapid phase*), the linear approximation of $F_t(a,x_0) - F_t(-a,x_0)$ is independent of $a$.

###### 3.1.1. The linear approximation

In the linear approximation, we neglect the effect of changes in allele frequency on the strength of stabilizing selection, assuming that

$$\overline{s}_S(a,x_0,t) = -\frac{a^2}{V_S}\cdot\overline{(1/2 - x.(a,x_0))(1-D_L^2/V_S)} \approx -\frac{a^2}{V_S}\cdot(1/2-x_0)\overline{(1-D_L^2/V_S)}. \qquad \text{S3.8}$$

We further assume that the integral effect of selection during the rapid phase is sufficiently small such that $(\overline{s}_D(a,t)+\overline{s}_S(a,x_0,t))\cdot t \ll 1$. Under these assumptions, we find that

$$F_t(a,x_0) \approx \frac{(\overline{s}_D(a,t)+\overline{s}_S(a,x_0,t))\cdot t}{a} \approx \frac{\left(\overline{s}_D(a,t)+\overline{s}_S^l(a,x_0,t)\right)\cdot t}{a}, \qquad \text{S3.9}$$

where $\overline{s}_D$ is defined in ***Eq. S3.5*** and

$$\overline{s}_S^l(a,x_0,t) \equiv -\frac{a^2}{V_S}\cdot\left(1-\overline{D_L^2}(t)/V_S\right)\cdot(1/2-x_0), \qquad \text{S3.10}$$

with

$$\overline{D_L^2}(t)/V_S\cdot t = 1/2\cdot\frac{\Lambda^2}{V_A}\left(1-D_L^2(t)/\Lambda^2\right). \qquad \text{S3.11}$$

This linear approximation captures the qualitative features of the allelic dynamics during the rapid phase (***Figs. 4 A***, ***S3.2***, ***S3.3*** and ***S3.4***).

###### 3.1.2. The nonlinear approximation

We build on the linear approximation to derive a nonlinear approximation that is more accurate, especially for large effect sizes, toward the end of the rapid phase. Specifically, we substitute the frequency based on the linear approximation, $x_t^l$, for $x_t$ in the expression for the average stabilizing selection coefficient (***Eq. S3.5***), to obtain:

$$\begin{aligned}\overline{s}_S^N(a,x_0,t) &\equiv -\frac{a^2}{V_S}\cdot\overline{(1/2-x_\cdot^l(a,x_0))\cdot(1-D_L^2/V_S)} \qquad \text{S3.12}\\ &= -\frac{a^2}{V_S}\cdot\left((1/2-x_0)(1-\overline{D_L^2}(t)/V_S) - ax_0(1-x_0)(\pm I_d(t) - a\cdot(1/2-x_0)\cdot I_s(t))\right),\end{aligned}$$

where

$$I_S(t) \equiv \frac{1}{2}\cdot\frac{t}{2V_S}\left(1-\overline{D_L^2}(t)/V_S\cdot\left(2-\overline{D_L^2}(t)/V_S\right)\right), \qquad \text{S3.13}$$

$$I_d(t) \equiv \frac{\Lambda}{V_A}\cdot\left(1-\overline{D_L}(t)/\Lambda\cdot\left(1+1/6\cdot\frac{\Lambda^2}{V_S}\cdot(1-D_L(t)/\Lambda)\cdot(1+2\cdot D_L(t)/\Lambda)\right)\right),$$

and with $\overline{D_L^2}(t)$ given in ***Eq. S3.11***. We obtain an explicit, nonlinear approximation by substituting $\overline{s}_S^N(a,x_0,t)$ for $\overline{s}_S(a,x_0,t)$ in our implicit form solution (***Eq. S3.3***), i.e.,

$$F_t(a,x_0) \approx \frac{1}{a}\cdot\left(\frac{\mathrm{Exp}\left[\left(\overline{s}_D(a,t)+\overline{s}_S^N(a,x_0,t)\right)\cdot t\right]-1}{1+x_0\left(\mathrm{Exp}\left[(\overline{s}_D(a,t)+\overline{s}_S^N(a,x_0,t))\cdot t\right]-1\right)}\right), \qquad \text{S3.14}$$

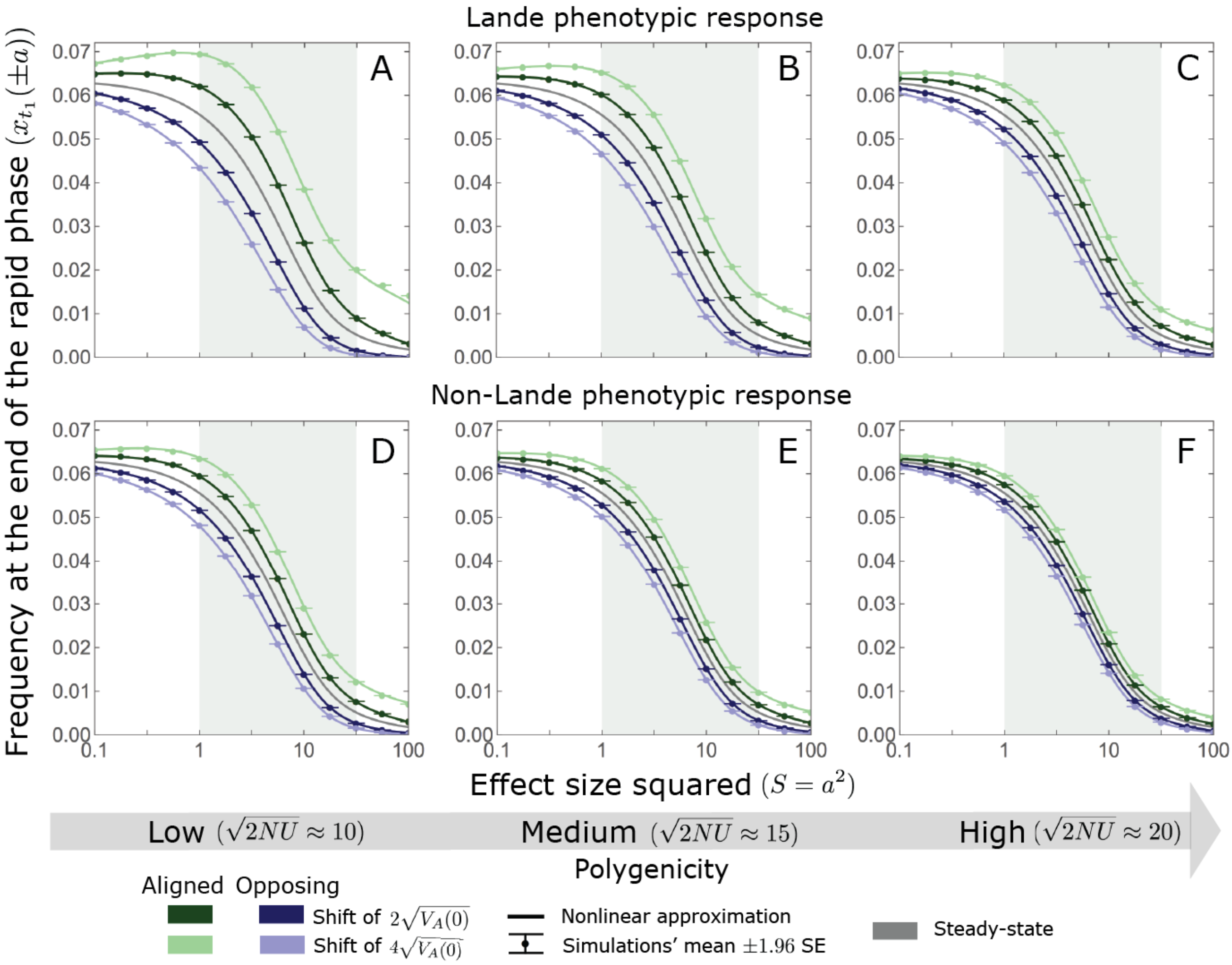

**Figure S3.1.** The nonlinear approximation accurately predicts average allele frequencies at the end of the rapid phase for a wide range of model parameters. We consider parameter values corresponding to the Lande (A-C) and non-Lande (D-F) phenotypic behaviors, with our standard parameter values for each case (*Section 5.2*). We examine a range of population mutational inputs, corresponding to low (A and D), moderate (B and E), and high (C and F) extent of polygenicity ($\sqrt{2NU}$), and two shift sizes (see legend). As expected, when polygenicity is lower and shifts are larger the expected changes to allele frequencies are greater (compare A with C and D with F). Perhaps less obvious, despite the fact that our nonlinear approximation relies on Lande's approximation, it performs well even in the cases with non-Lande phenotypic dynamics (D-F). The simulation results for each point were averaged over $2.5 \cdot 10^4$ alleles simulated with our individual allele simulations (OA) (*Section 5*), assuming $N = 10^4$.

with $\overline{s}_D$ given in ***Eq. S3.5***. The *nonlinear approximation* is quite accurate under a wide range of parameter values (***Figs. S3.2*** and ***S3.3***). It underestimates the increase in frequency of large effect, aligned alleles when the shift is large and polygenicity is low (***Fig. S3.2C***), because $|\overline{s}_S^N|$ is greater than $|\overline{s}_S|$ when directional selection is strong and lasts longer.

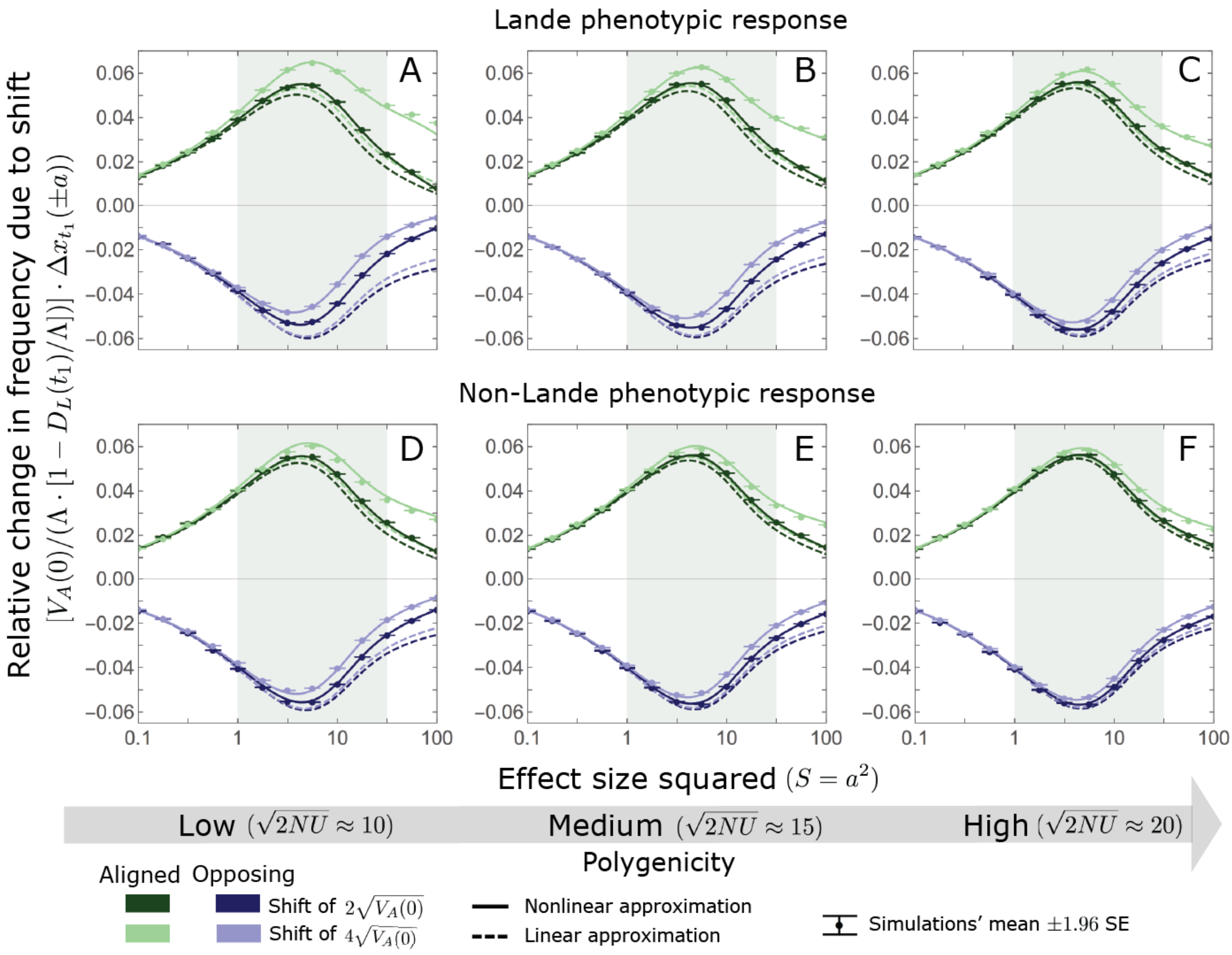

**Figure S3.2.** A comparison of the linear and nonlinear approximations with simulation results. The nonlinear approximation is quite accurate throughout and substantially more accurate than the linear approximation for larger effect sizes and low polygenicity (see, e.g., A, B, and D). In the linear approximation, frequency changes with directional selection alone scale with $\Lambda/V_A(0)\cdot(1-D_L(t_1)/\Lambda)$ (***Eqs. S3.5*** and ***S3.9***). We therefore normalized the frequency changes by this factor to make the results for the two shift sizes and three extents of polygenicity comparable. The model parameters and individual allele simulations (OA) are the same as in ***Fig. S3.1***.

#### 3.2. Contribution to phenotypic change

Phenotypic adaptation after the shift in optimum arises from the increase in frequency of alleles whose effects align with the shift relative to those with opposite effects. Here, we generalize the steps that we took in the main text in order to approximate the allelic contributions to change in mean phenotype. We begin by considering a pair of alleles that are initially minor, with the same initial frequency, and that have opposite effects of the same magnitude. The frequency difference between them during the rapid phase is given by

$$\Delta x_t^*\ (a, x_0) \equiv x_t\,(a, x_0) - x_t\,(-a, x_0) = 2ax_0\,(1 - x_0) \cdot \Delta F_t\,(a, x_0)\,, \qquad \text{S3.15}$$

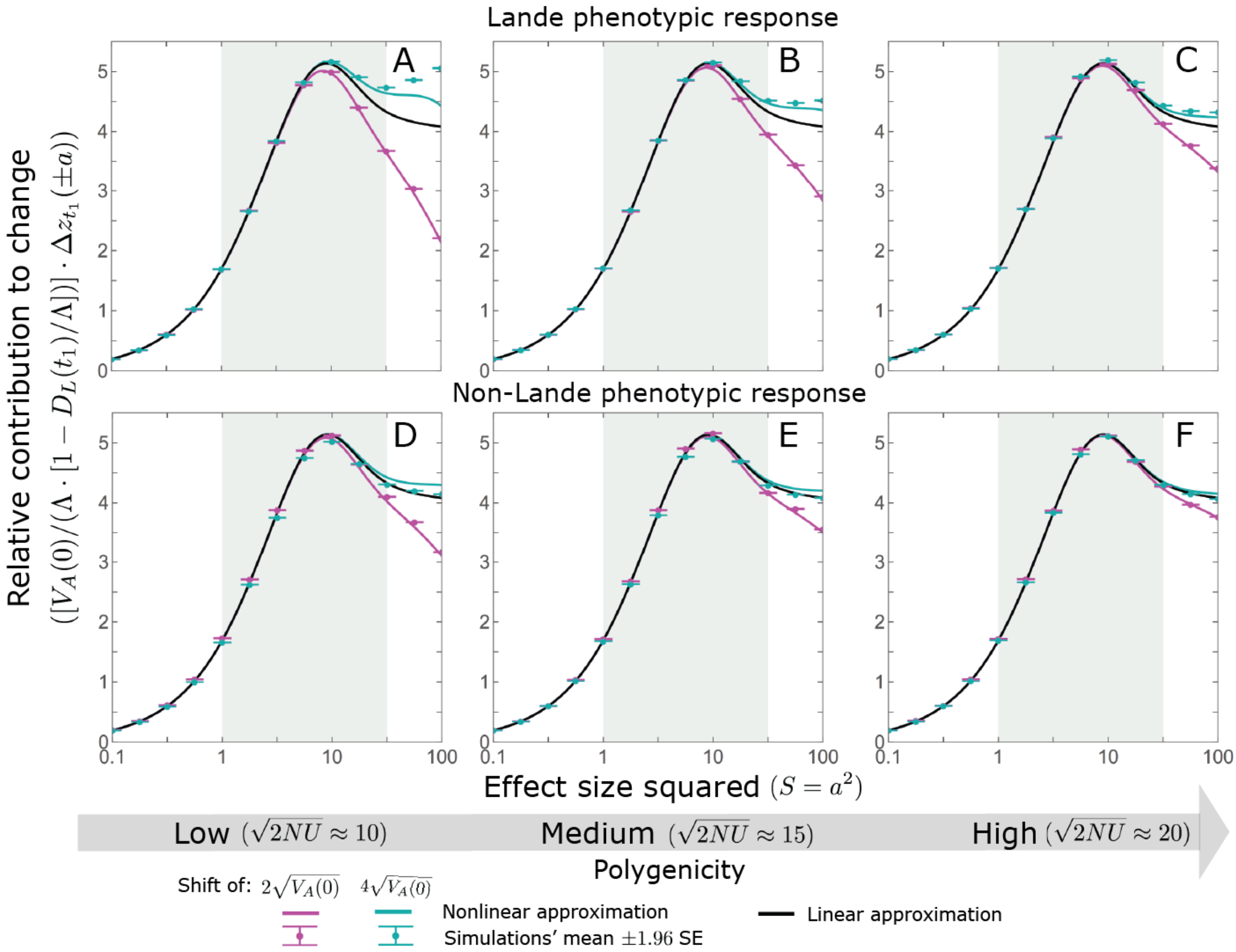

**Figure S3.3.** The allelic contribution to change in mean phenotype during the rapid phase. As for the frequency changes, the nonlinear approximation for the expected contribution to change is quite accurate throughout and substantially more accurate than the linear approximation for larger effect sizes and low polygenicity (see, e.g. A and B). In the linear approximation, the contribution to change in mean phenotype scales with $\Lambda/V_A(0)\cdot(1-D_L(t_1)/\Lambda)$ (see ***Eqs. S3.18*** and ***S3.19***). We therefore normalized the contributions by this factor to make them comparable for different initial variances and shift sizes. The model parameters and individual allele simulations (OA) are the same as in ***Fig. S3.1***.

where

$$\Delta F_t(a, x_0) \equiv (F_t(a, x_0) - F_t(-a, x_0))/2. \qquad \text{S3.16}$$

The pair's contribution to the change in mean phenotype is

$$\Delta z_t^*\ (a, x_0) = 2a \cdot \Delta x_t^*\ (a, x_0) = \ 2v^*\,(a, x_0) \cdot \Delta F_t(a, x_0), \qquad \text{S3.17}$$

where $v^*\,(a, x_0) = 2a^2 x_0(1 - x_0)$ is the initial contribution of each of the alleles to phenotypic variance. The expected contribution *per unit mutational input* of alleles with a given effect size and initial MAF follows from multiplying the contribution of a pair by the density of pairs, $\rho(a, x_0)$, namely,

$$\Delta z_t\,(a, x_0) = \rho(a, x_0) \cdot \Delta z_t^*\ (a, x_0) = v\,(a, x_0) \cdot \Delta F_t(a, x_0), \qquad \text{S3.18}$$

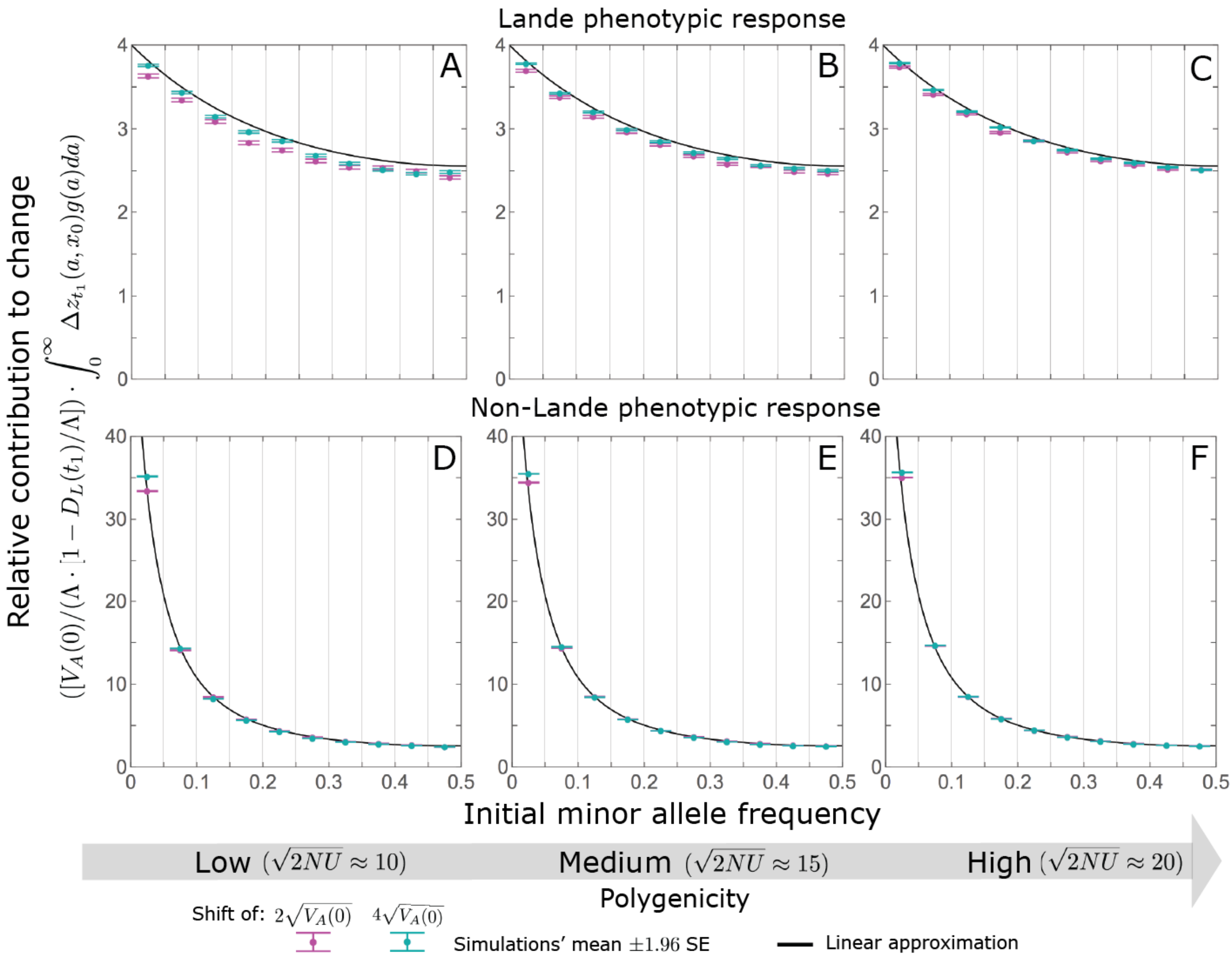

**Figure S3.4.** The relative contribution to change in mean phenotype during the rapid phase coming from different initial MAFs is well approximated in terms of the initial contribution to phenotypic variance, i.e., by the linear approximation (***Eq. 12***). The model parameters are the same as in ***Fig. S3.1*** and simulation results for each point were averaged over 2500 runs of our allelic simulations (AA). The results of these allelic simulations were binned by initial minor allele frequency (with boundaries between the bins marked by the light grey vertical lines) and the total contributions of alleles in frequency bin $(x_1, x_2]$ is divided by the total mutational input per generation times the size of the bin, $2NU(x_2 - x_1)$. As in ***Fig. S3.3***, we normalized the contributions by $\Lambda/V_A(0) \cdot (1 - D_L(t_1)/\Lambda)$ to make them comparable for different initial variances and shift sizes.

where $v(a, x_0) = 2\rho(a, x_0) \cdot v^*(a, x_0)$ is the density of genetic variance at steady-state (*Section 2.2*). Lastly, the expected total contribution of these alleles to phenotypic change follows from multiplying $\Delta z_t(a, x_0)$ by the mutational input per generation, $2NU \cdot g(a)$.

We can now use our approximations of $\Delta F_t(a, x_0)$ to obtain corresponding approximations for the allelic contributions to phenotypic change. Notably, in our linear approximation, $\Delta F_t$ takes the simple form

$$\Delta F_t \approx \Lambda / V_A \cdot (1 - D_L(t)/\Lambda), \qquad \text{S3.19}$$

which is independent of effect size and initial frequency. We use this approximation in our expressions for the phenotypic contribution in the main text. It captures the qualitative properties of the allelic contribution and is also fairly accurate for small and intermediate effect alleles. When the rapid phase is longer, the contributions of large effect alleles substantially deviate from the linear approximation in one of two ways. First, when the shift in optimum is large, large effect alleles contribute more than predicted by the linear approximation, because the change in frequency of the aligned allele accelerates as its frequency increases. Second, when the shift is small, large effect alleles contribute less than predicted, because the linear approximation underestimates the reduction in the frequency of both aligned and opposite alleles caused by stabilizing selection. Both of these effects are captured by the nonlinear approximation (see ***Fig. S3.3***).

#### 4. Allelic dynamics during equilibration

In the long run, phenotypic adaptation transitions from being based on small frequency differences between alleles whose effects align and oppose the shift in optimum to being based on small differences between the numbers of fixations of these opposite alleles (***Fig. 3***). Here we characterize the architecture of these long-term fixed differences. Specifically, we derive approximations for their relative contributions to phenotypic change as a function of allelic effect size and initial frequency (before the shift in optimum).

To this end, we approximate an allele's probability of fixation in two steps. First, we model the effect of directional selection on frequency as an instantaneous, deterministic pulse. We assume that the pulse occurs immediately after the shift for alleles already present in the population, and for mutations that occur after the shift, immediately after they arise. Second, we apply the diffusion approximation for the fixation probability, assuming stationary stabilizing selection and genetic drift and given the allele frequency immediately after the instantaneous pulse of directional selection.

We consider four approximations of increasing complexity, which differ in two of their assumptions. The first is whether the phenotypic approach to the new optimum is well approximated by Lande's solution; we refer to the corresponding approximations as *Lande* and *non-Lande*. The second is whether the expected changes to allele frequency due to directional selection are sufficiently small that we can apply linear approximations in calculating these changes and their effects on fixation probabilities; we refer to the corresponding approximations as *linear* and *nonlinear*. We consider each of the four approximations that follow from the combinations of these assumptions in turn.

##### 4.1. The linear Lande approximation

The linear Lande approximation is the simplest and the one that we detail in the main text. Here we briefly describe it for completeness and detail the conditions on model parameters under which we expect it to be accurate.

First, we approximate the change in allele frequency caused by directional selection. We use a deterministic approximation, which neglects the effects of drift, and in contrast to our approximations for the rapid phase, also neglects the effects of stabilizing selection. Under these simplifications,

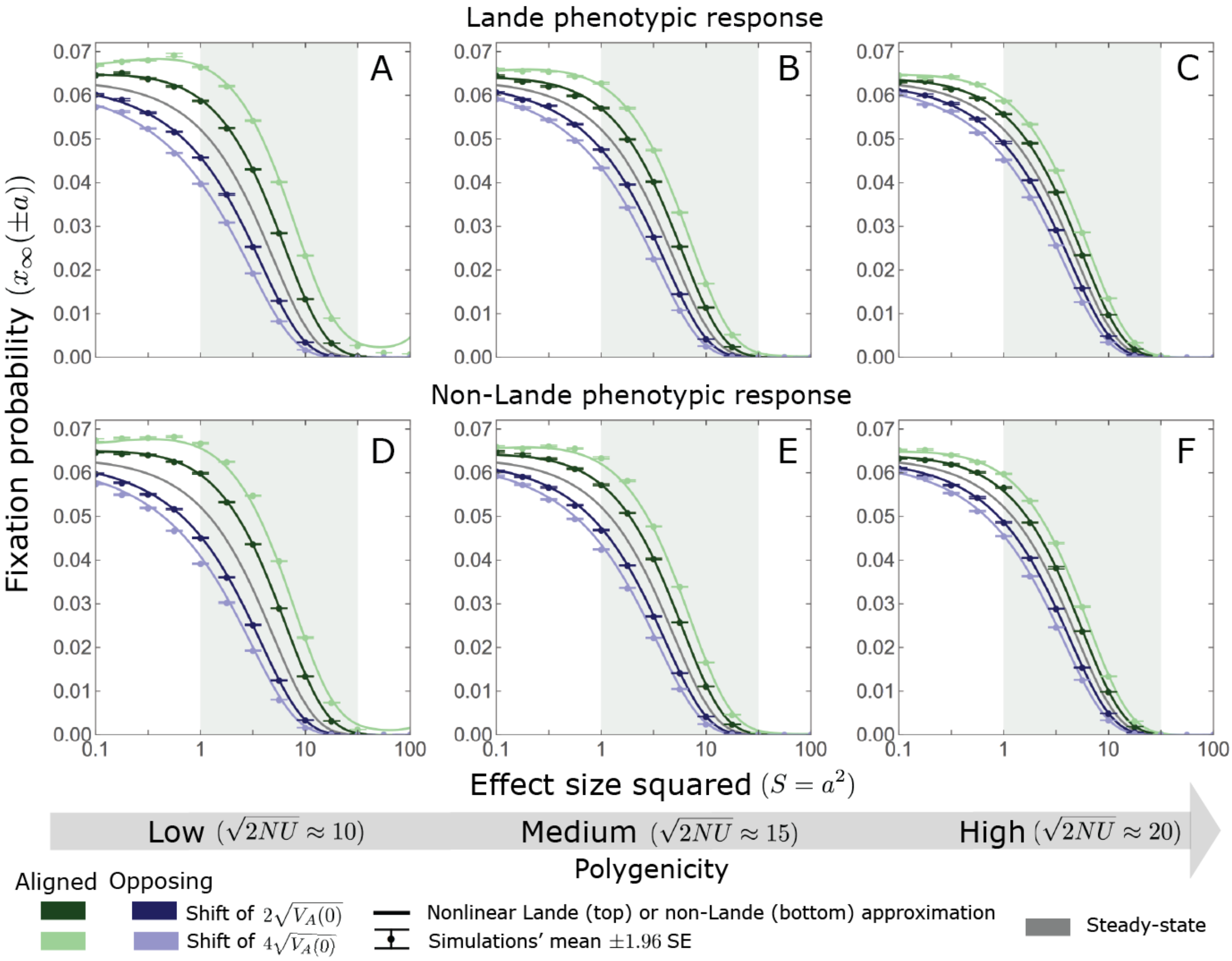

**Figure S4.1.** For small and intermediate alleles, changes in fixation probability are fairly small relative to the steady state fixation probability. The simulation results for each point were averaged over $2.5 \cdot 10^4$ alleles simulated with our individual allele simulations (OA) assuming $N = 10^4$, with our standard parameter values for the Lande and non-Lande cases (*Section 5.2*) and with the extent of polygenicity and shift sizes specified in the legend.

the change in allele frequency due to directional selection can be described by the ODE

$$\frac{dx}{dt} = \frac{s_d(t)}{2} x(1-x) \;\text{ with }\; s_d(t) = \pm 2a \cdot D(t)/V_S \tag{S4.1}$$

(see ***Eq. S3.2***). Further assuming that changes to allele frequency due to directional selection are small, we can approximate this ODE with

$$\frac{dx}{dt} = \frac{s_d(t)}{2} x_0(1-x_0), \tag{S4.2}$$

where $x_0$ is the initial frequency, before the shift in optimum. The total effect of directional selection is then given by integrating this ODE:

$$\Delta x_d(a, x_0) \equiv x_d(a, x_0) - x_0 \approx \; a x_0(1-x_0) \int_0^\infty (D(\tau)/V_S)\, d\tau. \tag{S4.3}$$

Here and throughout this section, we approximate the integral of the distance from the optimum by its expectation, where we assume that this expectation is finite and integrable; for brevity, we denote it by $\int D(\tau)d\tau$. Further assuming Lande's approximation for the change in mean phenotype

$$\int_0^\infty D(\tau)d\tau = \int_0^\infty D_L(\tau)\, d\tau = \Lambda \cdot V_S/V_A(0), \tag{S4.4}$$

implying that

$$\Delta x_d(a, x_0) \approx a x_0 (1 - x_0) \cdot \Lambda/V_A(0). \tag{S4.5}$$

We assume that this change in frequency occurs instantaneously, immediately after the shift in optimum.

Next, we approximate the fixation probability, using the diffusion approximation with stationary stabilizing selection and drift (***Eq. S2.4***) and the initial allele frequency after the instantaneous pulse due to directional selection (***Eq. S4.3***). We denote the expected allele frequency in the long run, which equals the fixation probability, by $x_\infty(a, x_0)$. Assuming that changes in frequencies due to directional selection are small, we can approximate their long-term effect on frequency (or fixation probability), by

$$\begin{aligned}\Delta x_\infty(a, x_0) &\equiv x_\infty(a, x_0) - \pi(a, x_0) = \pi(a, x_d(a, x_0)) - \pi(a, x_0) \approx \frac{\partial \pi}{\partial x}(a, x_0) \cdot \Delta x_d(a, x_0)\\ &= \frac{2f(a)}{v(a, x_0)} \cdot a x_0 (1 - x_0) \cdot \Lambda/V_A(0) = \frac{f(a)}{a} \cdot \frac{v^*(a, x_0)}{v(a, x_0)} \cdot \Lambda/V_A(0),\end{aligned} \tag{S4.6}$$

where $\partial_x \pi(a, x) = 2f(a)/v(a, x)$ with $f(a) \equiv 2a^3 \cdot \mathrm{Exp}\left(-a^2/4\right) / \left(\sqrt{\pi} \cdot \mathrm{Erf}(a/2)\right)$. The expected long-term fixed contribution to phenotypic change of a pair of opposite minor alleles with effect sizes $\pm a$ and initial frequency $x_0$ is then approximated by

$$\begin{aligned}\Delta z^*_\infty(a, x_0) &= 2a\left(x_\infty(a, x_0) - x_\infty(-a, x_0)\right) \approx \Delta z^*_d(a, x_0) \cdot \frac{\partial \pi}{\partial x}(a, x_0)\\ &= 2 \cdot \Lambda/V_A(0) \cdot f(a) \cdot \frac{2v^*(a, x_0)}{v(a, x_0)},\end{aligned} \tag{S4.7}$$

where $\Delta z^*_d(a, x_0) \equiv 2a(x_d(a, x_0) - x_d(-a, x_0))$. The contribution per unit mutational input of such pairs is approximated by

$$\begin{aligned}\Delta z_\infty(a, x_0) &= \Delta z^*_\infty(a, x_0) \cdot \rho(a, x_0) = 2 \cdot \Lambda/V_A(0) \cdot \frac{v^*(a, x_0) \cdot 2\rho(a, x_0)}{v(a, x_0)} \cdot f(a)\\ &= 2 \cdot \Lambda/V_A(0) \cdot f(a),\end{aligned} \tag{S4.8}$$

which is independent of the initial allele frequency. The expected marginal contribution per unit mutational input of alleles with effect sizes $\pm a$ is approximated by

$$\Delta z_\infty(a) = \int_0^{1/2} \Delta z_\infty(a, x_0)\, dx_0 \approx \frac{\Lambda}{V_A(0)} \cdot f(a). \tag{S4.9}$$

Lastly, the approximation for the total long-term contribution change in mean phenotype is

$$\begin{aligned}2NU \cdot \Delta z_\infty &= 2NU \cdot \int_0^\infty \Delta z_\infty(a) \cdot g(a)da \approx \Lambda \cdot \frac{2NU \cdot \int_0^\infty f(a) \cdot g(a)da}{V_A(0)}\\ &= \Lambda \cdot \frac{\int_0^\infty f(a) \cdot g(a)da}{\int_0^\infty v(a) \cdot g(a)da} = \frac{\Lambda}{1 + C},\end{aligned} \tag{S4.10}$$

where $C \equiv \frac{\int_0^\infty v(a)\cdot g(a)da}{\int_0^\infty f(a)\cdot g(a)da} - 1$.

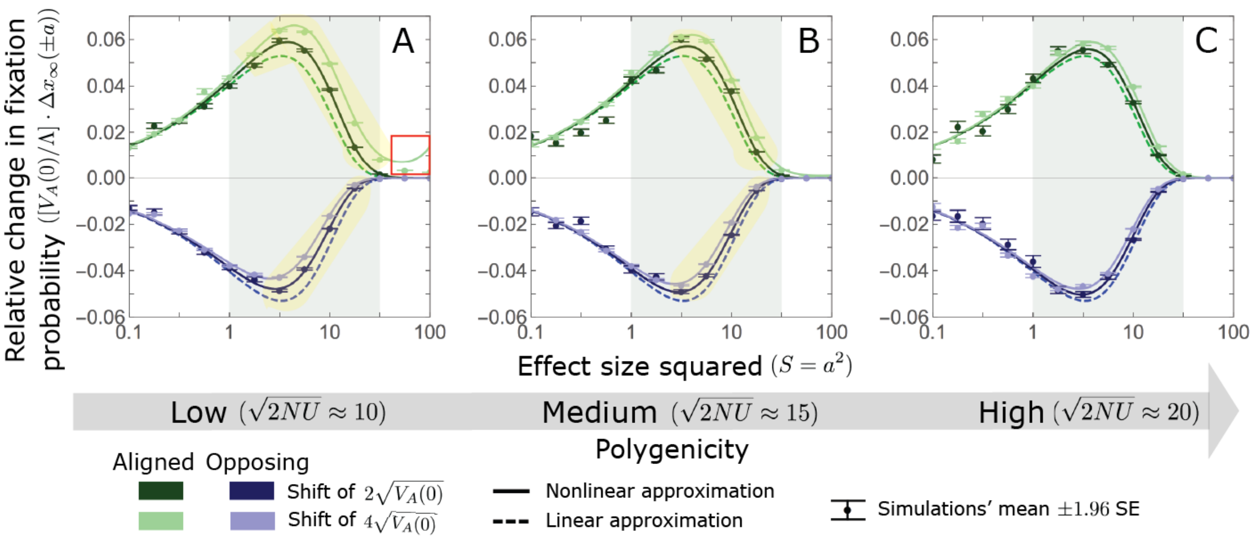

**Figure S4.2.** The Lande case: A comparison the linear and nonlinear approximations for the expected change in fixation probability of alleles already present at the time of the shift. The nonlinear Lande approximation performs better than the linear Lande approximation for alleles with large and intermediate effects, especially when polygenicity is lower (see the golden shaded regions in A and B). In the linear approximation, the change in fixation probability scales with $\Lambda/V_A(0)$ (*Eq. S4.6*). We therefore normalized the changes in fixation probability by this factor to make them comparable for different initial variances and shift sizes. Note that with this normalization, the linear curves for the two shifts overlap, which is why only one is visible. The simulation results for each point were averaged over $2.5 \cdot 10^4$ alleles simulated with our individual allele simulations (OA) assuming $N = 10^4$, with our standard parameter values for the Lande case (*Section 5.2*) and with the extent of polygenicity and shift sizes specified in the legend.

This leads to the first of two conditions under which we expect the linear Lande approximation to be accurate. In the long run, we expect the total change in mean phenotype to be equal to the shift in optimum. *Eq. S4.10* implies that a necessary condition for it to be the case, and thus for our approximation to be accurate, is that

$$C \equiv \frac{\int_0^\infty v(a)\cdot g(a)da}{\int_0^\infty f(a)\cdot g(a)da} - 1 \ll 1. \qquad \text{S4.11}$$

This condition implies that the bulk of genetic variance before the shift arises from alleles with fairly small effects (because $f(a) \approx v(a)$ for $a^2 \lessapprox 4$, but $f$ is substantially smaller than $v$ for larger effect sizes; *Fig. 5*). The second condition ensures that our linear expansions are accurate. In *Section 4.3*, we show that it will be the case when

$$|a| \cdot \Lambda/V_A(0) \ll 1. \qquad \text{S4.12}$$

For alleles with $|a| \ll 2$, this condition always holds under our conditions on parameters, since $\Lambda/V_A(0) \lessapprox 1/2$ (*Eq. S2.16*). For alleles with larger effects, this condition should hold when: i) the shift in optimum is not too large compared to the phenotypic standard deviation (e.g., $\Lambda/\sqrt{V_A(0)} \sim 1$); ii) the trait is sufficiently polygenic (specifically, that $\sqrt{2NU} \gg |a|$; because $\sqrt{V_A(0)} \sim \sqrt{2NU}$ under our assumption that a substantial proportion of mutations are not effectively neutral—see

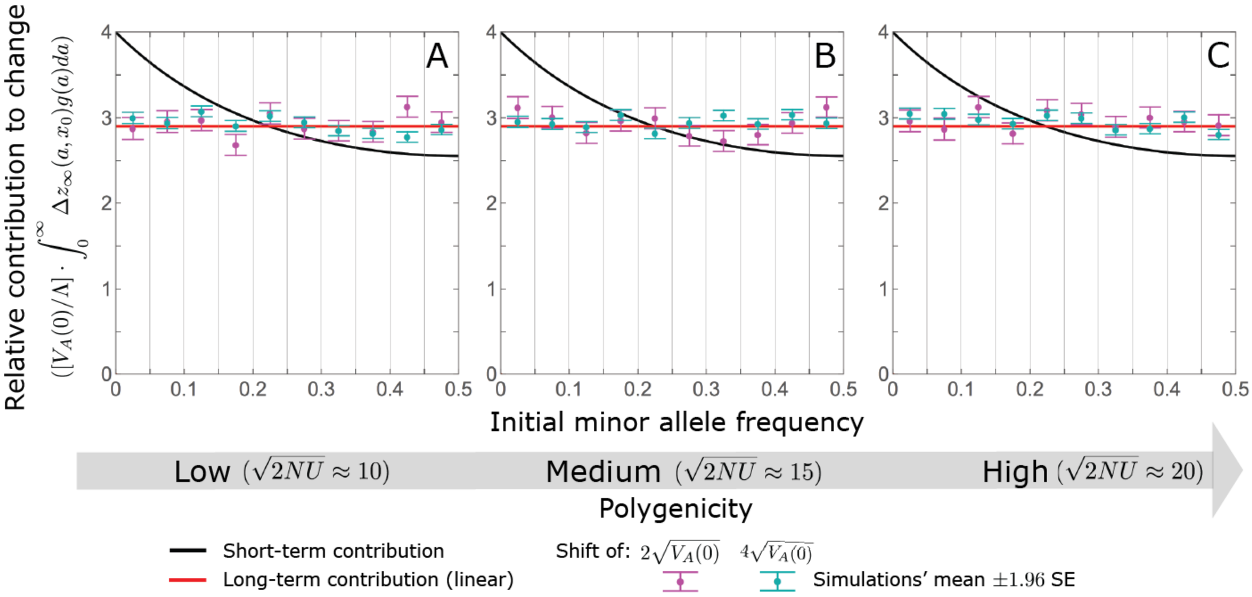

**Figure S4.3.** The Lande case: the long-term relative contribution to the change in mean phenotype coming from different initial MAFs is approximately constant. Model parameters are the same as in ***Fig. S4.2*** and simulation results for each point were averaged over 2500 runs of our allelic simulations (AA). The results of these allelic simulations were binned by initial minor allele frequency (with boundaries between the bins marked by the light grey vertical lines). The total contributions of alleles in frequency bin $(x_1, x_2]$ is divided by the total mutational input per generation times the size of the bin, $2NU(x_2 - x_1)$.

***Eq. S2.14***). With regards to (ii), we note that $\sqrt{2NU} \gg 1$ (by assumption; see ***App. A Table 2***) and that when condition (***S4.11***) holds, we already expect the bulk of long-term contributions to phenotypic adaptation to come from alleles with $|a| \leq 2$. ***Figs. S4.2-S4.4*** illustrate that the linear Lande approximation is accurate when conditions (***S4.11***) and (***S4.12***) are met.

While we do not prove that our conditions are sufficient, we conjecture that they are, for the following reasons. First, in *Section 6* and 7 we illustrate by simulation that Lande's approximation is accurate when condition (***S4.11***) is met. Intuitively, when the bulk of variation has fairly small effect sizes, we expect little build-up of a third phenotypic moment during the rapid phase (also see *Section 2.3*). When Lande's approximation applies, directional selection has non-negligible effects for only a brief period after the shift in optimum (for $\sim 1/U$ generations; see section on *allele dynamics*). Consequently, these effects should be well approximated as an instantaneous pulse that neglects drift and stabilizing selection, whose effects manifest over longer timescales. Additionally, newly arising mutations should contribute negligibly to phenotypic change. This is because fairly few new mutations arise while directional selection is effective (i.e. during the rapid phase), and because the fixation probabilities (let alone the difference in fixation probabilities between new mutations with opposing effects) of those that do are minute, given that they start from an initial frequency of $1/(2N)$.

Lastly, we require that our first order approximations for the change in allele frequency (***Eq. S4.3***) and change in fixation probability (***Eq. S4.6***) be accurate. With high polygenicity and a moderate shift, we expect the average contribution to phenotypic change per segregating allele to be small. Nevertheless, large effect alleles could experience changes in frequency that are large relative to

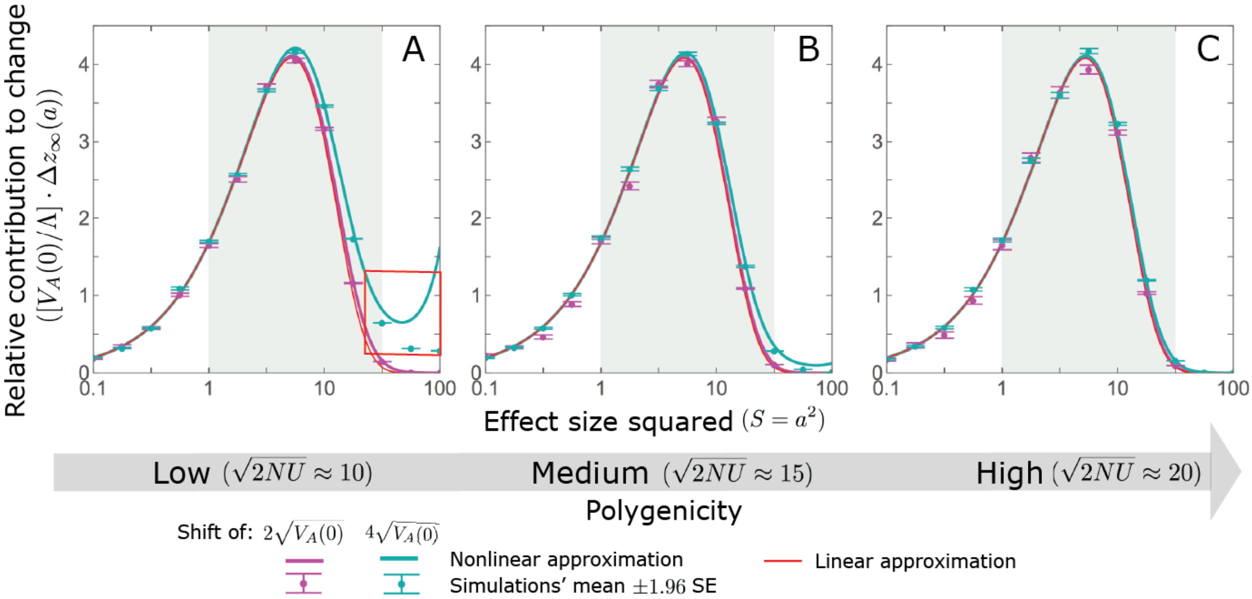

**Figure S4.4.** The Lande case: As predicted by the linear approximation, the long-term relative contribution to the change in mean phenotype coming from standing variation scales with $\Lambda/V_A(0)$. Both the linear and nonlinear Lande approximation are fairly accurate for alleles with small effects. When polygenicity is low, the nonlinear Lande approximation is more accurate for alleles with intermediate effects (see the blue line in A and B). Both approximations perform badly for large effect alleles when the polygenicity is low and the shift is large (the red box in A). However, in the Lande case, we expect very few alleles to have large effect sizes. The model parameters and individual allele simulations (OA) are the same as in ***Fig. S4.2***.

their low initial frequencies, causing a substantial relative error in our linear instantaneous pulse approximation (***Eq. S4.3***). This reasoning provides some intuition for why condition (***S4.12***) combines requirements on the extent of polygenicity, shift size, and effect size. In *Section 4.3*, we show why this particular combination is required.

#### 4.2. The linear non-Lande approximation

Next, we derive approximations for the case in which our linear expansions are accurate but Lande's approximation is not. Lande's approximation for the mean performs poorly when alleles with large effect sizes $a^2 \geq 4$ contribute substantially to genetic variance before the shift, or more precisely when $C$ is substantial (see ***Eq. S4.10***). As we describe in the main text, in this case, the phenotypic distribution develops a substantial third moment during the rapid phase, leading to a much slower approach to the new optimum afterwards. This results in prolonged weak directional selection that amplifies the difference in fixation probabilities between alleles with opposite effects present before the shift in optimum. In addition, despite it being weak, directional selection acts for so long (on the order of $N$ generations; see ***Figs. S6.1*** and ***S7.2***) that it can produce a substantial cumulative difference between the numbers of fixations of mutations with opposite effects that arise after the shift in optimum. In what follows, we denote the proportions of the long-term (fixed) contribution to phenotypic change that arise from standing variation and from new mutations by $\eta$ and $1 - \eta$ respectively.

###### 4.2.1. Standing variation

First, we consider the contribution from standing variation. In this case, the integral effect of directional selection is greater than under Lande's approximation,

$$\int_0^\infty D(\tau)d\tau > \int_0^\infty D_L(\tau)\,d\tau = \Lambda \cdot V_S/V_A(0). \qquad \text{S4.13}$$

Since directional selection acts on the order of $N$ generations after the shift, an allele present at the time of the shift may not experience the full integral effect of directional selection. We define $t_*$ as the effective number of generations over which an allele is subject to directional selection and $A > 0$ such that

$$\int_0^{t_*} D(\tau)d\tau = (1+A)\cdot \int_0^\infty D_L(\tau)\,d\tau = (1+A)\cdot \Lambda \cdot V_S/V_A(0). \qquad \text{S4.14}$$

By analogy with the derivations of the previous section, the instantaneous pulse approximation for the effect of directional selection is

$$\Delta x_d\,(a,x_0) \approx\ a x_0\,(1-x_0)\cdot \int_0^{t^*} \left(D\,(\tau)\,/V_S\right)d\tau = a x_0\,(1-x_0)\cdot (1+A)\cdot \Lambda/V_A(0). \qquad \text{S4.15}$$

The expressions for the fixed contribution are also analogous to those derived in the previous section, where here we replace $\Lambda/V_A(0)$ by $(1+A)\cdot\Lambda/V_A(0)$. In particular, the expected long-term (fixed) contribution to phenotypic change per unit mutational input of pairs of opposite alleles with effect sizes $\pm a$ and initial MAF $x_0$ is approximated by

$$\Delta z_\infty\,(a,x_0) = \Delta z_d\,(a,x_0)\cdot \frac{2f\,(a)}{v\,(a,x_0)} \approx (1+A)\cdot \frac{\Lambda}{V_A(0)}\cdot 2f(a), \qquad \text{S4.16}$$

where $\Delta z_d\,(a,x_0) \equiv 2a(x_d\,(a,x_0) - x_d\,(-a,x_0))\cdot \rho(a,x_0) \approx v\,(a,x_0)\cdot(1+A)\cdot\Lambda/V_A(0)$. Similar to what we obtained for the linear Lande approximation, this contribution is independent of the initial allele frequency. The expected marginal contribution per unit mutational input of pairs of alleles with effect size $\pm a$ is approximated by

$$\Delta z_\infty^v\,(a) = \int_0^{1/2} \Delta z_\infty\,(a,x_0)\,dx_0 \approx (1+A)\cdot\Lambda/V_A(0)\cdot f(a). \qquad \text{S4.17}$$

Lastly, the total fixed contribution to phenotypic change from standing variation is approximated by

$$2NU\cdot\Delta z_\infty^v = 2NU\cdot\int_0^\infty \Delta z_\infty^v(a)\cdot g(a)da \approx \frac{2NU\cdot\int_0^\infty f(a)\cdot g(a)da}{V_A(0)}\cdot(1+A)\cdot\Lambda = \frac{1+A}{1+C}\cdot\Lambda. \qquad \text{S4.18}$$

Thus, we find that the proportional long-term contribution from standing variation is

$$\eta = \frac{1+A}{1+C}. \qquad \text{S4.19}$$

In this case, the justification for the instantaneous pulse approximation is less obvious, because directional selection substantially affects allele frequencies over an extended period. Assuming that the pulse approximation is accurate nonetheless, the same reasoning that we employed for the linear Lande approximation suggests that our linear expansions are accurate when polygenicity is sufficiently high and the shift and magnitude of effect size are not too large. However, given that directional selection acts for longer, we expect the condition on these quantities to be more stringent than for the linear Lande approximation. By analogy, we would guess that this condition becomes $|a| \cdot (1 + A) \cdot \Lambda/V_A(0) \ll 1$ (see *Section 4.4*).

To test the linear non-Lande approximation for standing variation, we estimate the proportional contribution of standing variation to phenotypic change, $\eta$, using our AA simulations, and then calculate $A$ using ***Eq. S4.19***. We find the approximation to be quite accurate provided high polygenicity and a moderate shift (***Figs. S4.5-S4.7***). However, when polygenicity is low and/or the shift is large, it underestimates the fixation probabilities of both aligned and opposing intermediate effect alleles (see the golden shaded region in ***Fig. S4.5***). The nonlinear non-Lande approximation performs better in some of these cases (see *Section 4.4*).

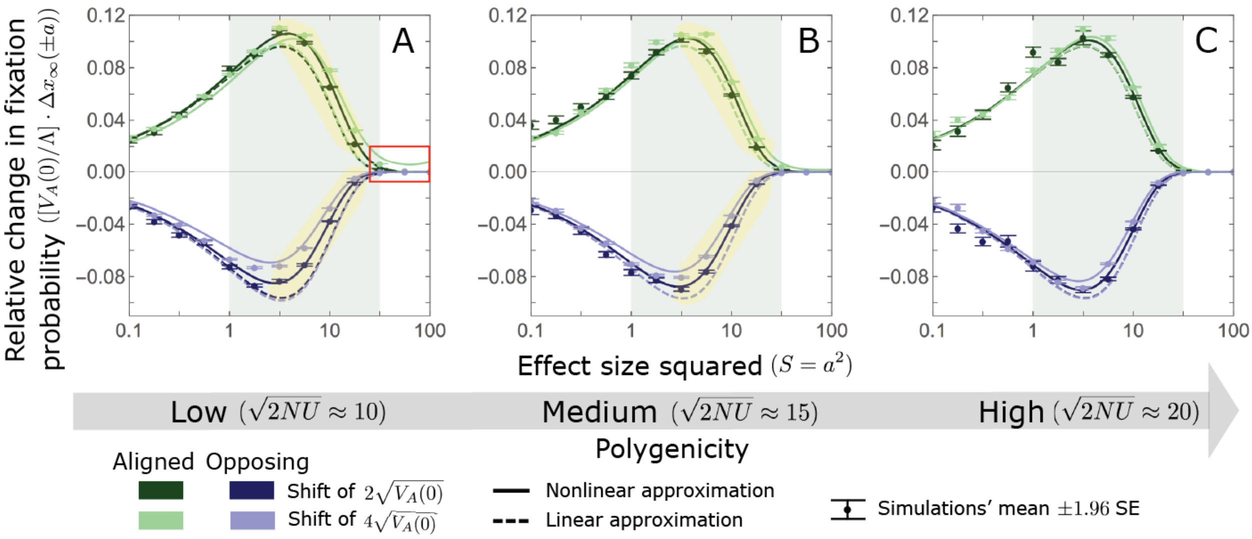

**Figure S4.5.** The non-Lande case: A comparison of the linear and nonlinear approximations for the expected change in fixation probability of alleles segregating at the time of the shift. The nonlinear approximation performs better the linear one for intermediate effect alleles when polygenicity is low (see the gold shaded regions in A and B). In the linear approximation, the change in fixation probability scales with $\Lambda/V_A(0)$ (***Eq. S4.6***). We normalized the changes in fixation probability by this factor to make them comparable for different initial variances and shift sizes. With this normalization, the linear curves for the two shifts are almost identical, which is why only is visible. The simulation results for each point were averaged over $2.5 \cdot 10^4$alleles simulated with our individual allele simulations (OA) assuming $N = 10^4$, with our standard parameter values for the non-Lande case (*Section 5.2*) and with the extent of polygenicity and shift sizes specified in the legend.

###### 4.2.2. New Mutations

First, we consider a new mutation with effect size $a$ that arose $t$ generations after the shift in optimum. The instantaneous pulse approximation for the effect of directional selection on such a

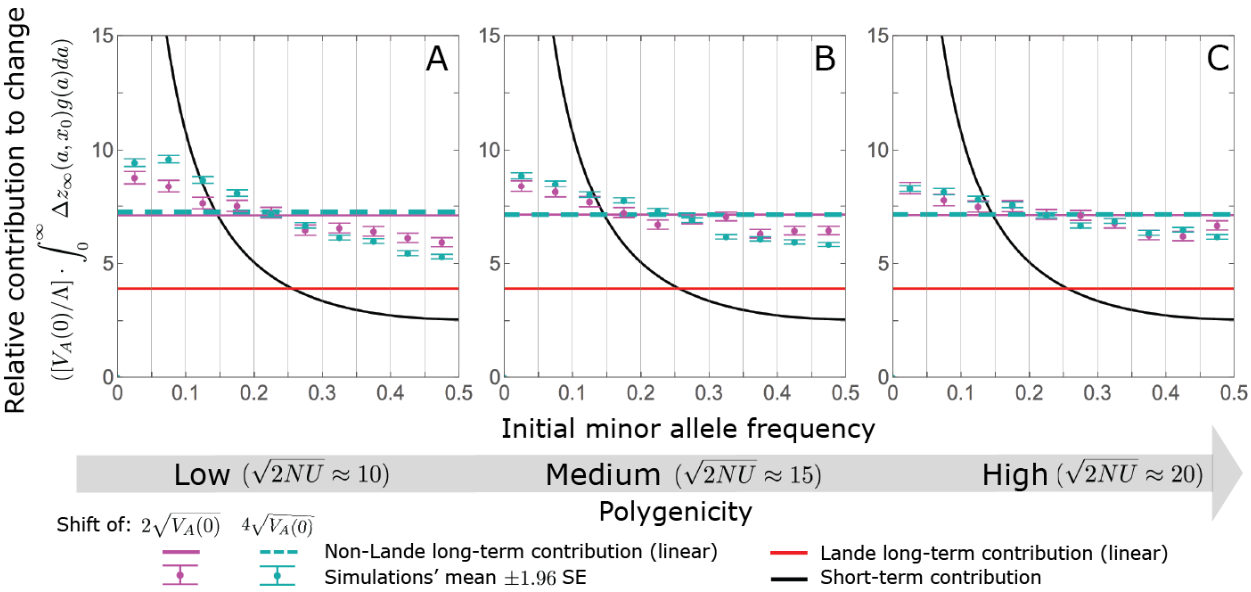

**Figure S4.6.** The non-Lande case: The long-term relative contribution to the change in mean phenotype coming from different initial MAFs is approximately constant, with low initial MAF alleles contributing slightly more than high initial MAF alleles. The slight excess in the contribution of alleles with low initial MAF is especially pronounced when the shift is large and the polygenicity is low. Model parameters are the same as in ***Fig. S4.5*** and simulation results for each point were averaged over 2500 runs of our allelic simulations (AA). We calculate the relative contribution of alleles in each MAF bin $(x_1, x_2]$ (between the gray gridlines) by dividing the contribution of all fixations in the bin by the total mutational input per generation times the size of the bin, $2NU(x_2 - x_1)$.

mutation is

$$\Delta x_d(a|t) \approx a\frac{1}{2N}\left(1 - \frac{1}{2N}\right) \cdot \int_t^{t+\Delta_t} (D(\tau)/V_S)\, d\tau. \qquad \text{S4.20}$$

where $\Delta_t$ is the effective number of generations over which an allele arising in generation $t$ is subject to directional selection. The expected change in mean phenotype caused by a pair of such mutations with opposite effects $\pm a$ is

$$\Delta z_d^*(a|t) \approx 2v^*(a, 1/2N) \cdot \int_t^{t+\Delta_t} (D(\tau)/V_S) d\tau. \qquad \text{S4.21}$$

Assuming that $a \cdot \Delta x_d(a|t) \ll 1$, we can approximate the expected long-term (fixed) effect of such a pair by

$$\begin{aligned}\Delta z_\infty^*(a|t) &\approx 2v^*(a, 1/(2N)) \cdot \int_t^{t+\Delta_t} (D(\tau)/V_S)\, d\tau \cdot \frac{2f(a)}{v(a, 1/(2N))}\\ &= \frac{2f(a)}{\rho(a, 1/(2N))} \cdot \int_t^{t+\Delta_t} (D(\tau)/V_S)\, d\tau\\ &= 2f(a) \cdot \int_t^{t+\Delta_t} (D(\tau)/V_S)\, d\tau/(2N), \qquad \text{S4.22}\end{aligned}$$

where $2v^*(a, 1/(2N))/v(a, 1/(2N)) = \rho^{-1}(a, 1/(2N)) = 1/(2N)$. Multiplying this contribution by the number of such pairs per unit mutational input per generation, 1/2, we find that the expected

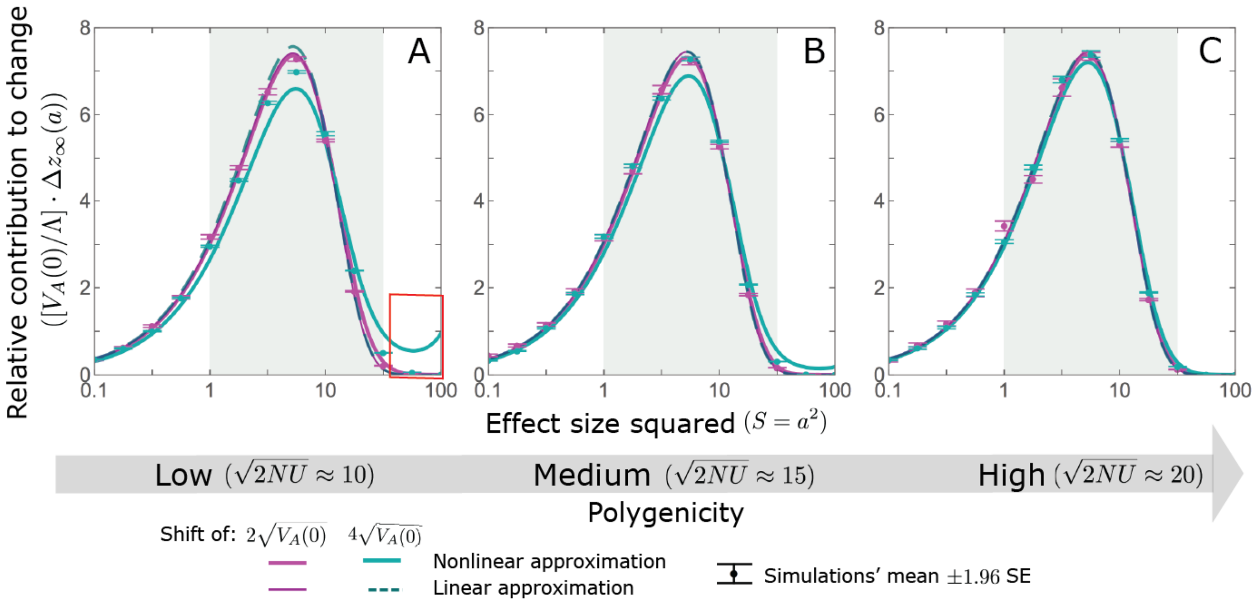

**Figure S4.7.** The non-Lande case: As predicted by the linear approximation, the long-term relative contribution to the change in mean phenotype coming from standing variation scales with $\Lambda/V_A(0)$. Both the linear and nonlinear approximations predict the long-term relative contribution to the change in mean phenotype fairly well. However, when polygenicity is low and the shift is large, the nonlinear approximation overestimates the proportion of the change in mean due to large effect alleles (see the red box in A). The model parameters and individual allele simulations (OA) are the same as in ***Fig. S4.5***.

contribution of pairs per unit mutational input is approximated by

$$\Delta z_\infty (a|t) \approx f(a) \left( \frac{\int_t^{t+\Delta_t} (D(\tau)/V_S)\, d\tau}{2N} \right). \tag{S4.23}$$

Next, considering all mutations that arise after the shift in optimum, we find that the expected long-term contribution of mutations with effect of magnitude $a$ is approximated by

$$\Delta z_\infty^m (a) = \int_0^\infty \Delta z_\infty (a|t)\mathrm{dt} \ \approx f(a) \cdot B \cdot \Lambda/V_A(0). \tag{S4.24}$$

Notably, we find that the relative contribution of mutations of any given effect size (and that arise at any time) is the same as for standing variation (i.e., proportional to $f(a)$).

We can now relate our approximations for new mutations with their total long-term (fixed) contribution to phenotypic change. To this end, we express ***Eq. S4.24*** as

$$\Delta z_\infty^m (a) \approx B \cdot \Delta z_\infty^{lL} (a) = B \cdot f(a) \cdot \Lambda/V_A(0), \tag{S4.25}$$

where $\Delta z^{lL}$ is the linear Lande approximation for the corresponding contribution from standing variation and $B \equiv \frac{\int_0^\infty \int_t^{t+\Delta_t} (D(\tau)/V_S) d\tau dt/(2N)}{\Lambda/V_A(0)}$. In these terms, our approximation for the total (fixed) contribution to phenotypic change from new mutations is

$$2NU \cdot \Delta z_\infty^m = 2NU \cdot \int_0^\infty \Delta z_\infty^m(a) \cdot g(a)da \approx \frac{2NU \cdot \int_0^\infty f(a) \cdot g(a)da}{V_A(0)} \cdot B \cdot \Lambda = \frac{B}{1+C} \cdot \Lambda. \tag{S4.26}$$

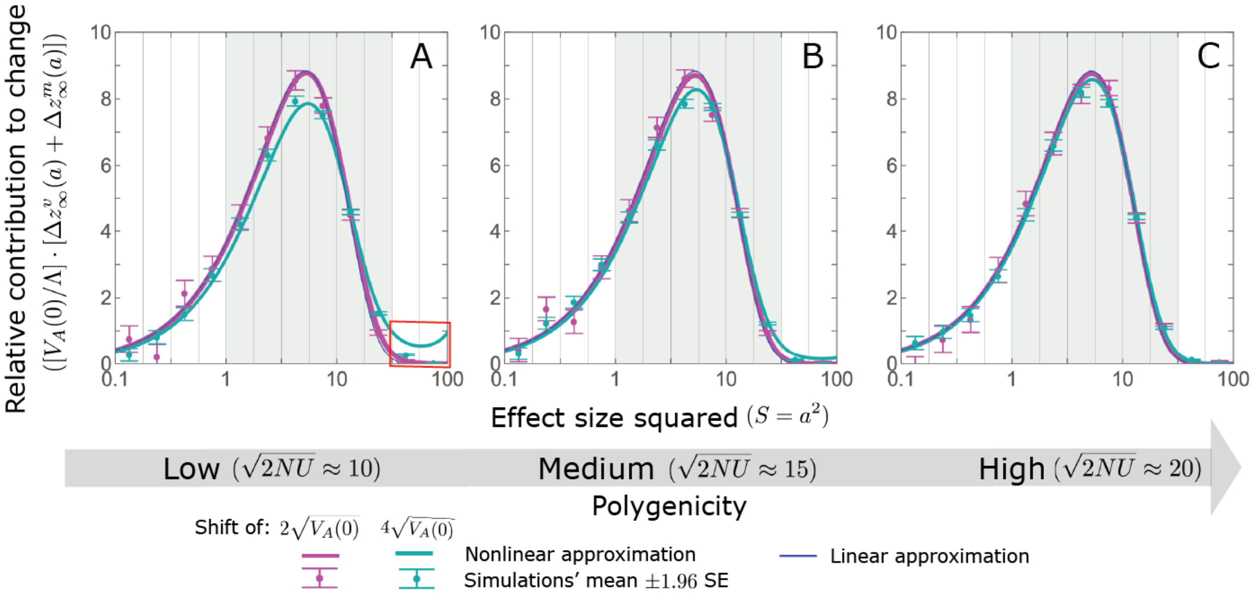

**Figure S4.8.** The non-Lande case: As predicted by the linear approximation, the long-term contribution, per unit mutational input, to the change in mean phenotype coming from standing variation and new mutations scales with $(1+C)\cdot\Lambda/V_A(0)$ (***Eq. S4.27***). Both the linear and nonlinear approximations perform fairly well. However, when the polygenicity is low and the shift is large, the nonlinear approximation overestimates the proportion of the change in mean due to large effect standing variation (***Fig. S4.7***) and therefore overestimates the proportion contribution coming from *all* large effect alleles (see the red box in A). The model parameters are the same as in ***Fig. S4.5*** and the simulation results were averaged over 2500 runs of our allelic simulations (AA). We calculate the relative contribution of alleles in each effect size bin (between the gray gridlines) by dividing the contribution of all fixations in the bin by the mutation rate per generation corresponding to that bin.

Thus, we find that the proportional long-term contribution from new mutations is $1-\eta = B/(1+C)$. Further recalling that $\eta = (1+A)/(1+C)$, we also find that $A + B = C$.

We expect the linear non-Lande approximation for new mutations to be accurate under more general conditions than it is for standing variation. First, consider our linear expansion of the fixation probability (e.g., ***Eq. S4.6***), which is accurate when $a \cdot \Delta x_d(a|t) \ll 1$. We expect this to be the case for most if not all new mutations and model parameter ranges that meet our conditions (***App. A Table 2***), because new mutations have a tiny initial frequency of $1/(2N)$ and the vast majority of them arise during the equilibration phase, when $D$ is very small. Second, consider the condition under which our linear approximation for $\Delta x_d(a|t)$ (***Eq. S4.20***) is accurate. As we already noted (and argue in *Section 4.4*), we expect the linear non-Lande approximation for standing variation to be accurate when $|a| \cdot (1+A) \cdot \Lambda/V_A(0) \ll 1$. An analogous argument suggests this is case for new mutations when $|a| \cdot \int_t^{t+\Delta_t} (D(\tau)/V_S)\, d\tau \ll 1$. However, $\int_t^{t+\Delta_t} (D(\tau)/V_S)\, d\tau \le \int_0^{t_*} (D(\tau)/V_S)\, d\tau = (1+A)\cdot\Lambda/V_A(0)$, since $D(t)$ decreases after the shift and alleles segregating at the time of the shift tend to segregate longer than new mutations (i.e., $t_* > \Delta_t$). Thus, with the exception of a few mutations that arise shortly after the shift, we expect that $|a| \cdot \int_t^{t+\Delta_t} (D(\tau)/V_S)\, d\tau \ll |a| \cdot (1+A) \cdot \Lambda/V_A(0)$.

###### 4.2.3. Standing variation and new mutations

To test the linear non-Lande approximation for both standing variation and new mutations, we note that their joint contribution per unit mutational input of alleles with effect sizes $\pm a$ is approximated by

$$\Delta z_{\infty}^{v}(a) + \Delta z_{\infty}^{m}(a) \approx [(1 + A + B) \cdot \Lambda/V_A(0)] \cdot f(a) = (1 + C) \cdot f(a) \cdot \Lambda/V_A(0), \qquad \text{S4.27}$$

where we have closed form expression for all parts of this expression. As we illustrate in ***Fig. S4.8***, the linear non-Lande approximation for this joint contribution is quite accurate provided high polygenicity and a moderate shift. However, when polygenicity is low and/or the shift is large, the linear approximation overestimates the contribution of intermediate effect alleles and underestimates the contribution of large effect alleles.

##### 4.3. The nonlinear Lande approximation

When directional selection causes large changes in allele frequencies our linear approximations becomes less accurate. as expected when trait polygenicity is low, the shift is large (relative to the initial phenotypic standard deviation) or the magnitude of effect size is large (relative to $\delta$). Here, we develop a nonlinear Lande approximation that is more accurate (especially for intermediate effect alleles) than the linear one when changes to allele frequencies due to directional selection are large, and we derive the condition under which the improvement is substantial.

Our derivation follows the same steps that we took in previous sections but without relying on linear expansions. First, we derive a nonlinear approximation for the effects of directional selection, once again assuming that they occur as an instantaneous pulse. To this end, we apply the same kind of nonlinear approximation that we used for the rapid phase (*Section 3.1*), but in this case we ignore the effects of stabilizing selection. By analogy to our derivation in *Section 4.1*, we express the change in frequency due to directional selection as

$$\Delta x_d(a, x_0) = x_d(a, x_0) - x_0 = a x_0 (1 - x_0) \cdot F_d(a, x_0), \qquad \text{S4.28}$$

where

$$\begin{aligned} F_d(a, x_0) &\equiv \frac{1}{a}\left(\frac{\operatorname{Exp}\left[a \cdot \int_0^{\infty} (D_L(\tau)/V_S) d\tau\right] - 1}{1 + x_0 \cdot \left(\operatorname{Exp}\left[a \cdot \int_0^{\infty} (D_L(\tau)/V_S) d\tau\right] - 1\right)}\right) \\ &= \frac{1}{a}\left(\frac{\operatorname{Exp}\left[a \cdot \Lambda/V_A(0)\right] - 1}{1 + x_0 \cdot \left(\operatorname{Exp}\left[a \cdot \Lambda/V_A\right] - 1\right)}\right). \end{aligned} \qquad \text{S4.29}$$

Similar to our previous derivations, we approximate fixation probabilities using the frequencies after the instantaneous pulse as an initial condition, but in this case, we do not use a linear expansion of the fixation probability in $\Delta x_d$. In particular, we approximate the expected long-term (fixed) contribution to phenotypic change of a pair of opposite alleles with effect sizes $\pm a$ and initial MAF $x_0$ by

$$\Delta z_{\infty}^{*}(a, x_0) = 2a \cdot \Delta x_{\infty}^{*}(a, x_0) \approx 2a(x_{\infty}(a, x_0) - x_{\infty}(-a, x_0)) \qquad \text{S4.30}$$

and the contribution of such pairs per unit mutational input by

$$\Delta z_\infty(a, x_0) \approx \Delta z^*_\infty(a, x_0) \cdot \rho(a, x_0). \tag{S4.31}$$

We can also write down integral forms for the marginal contribution of alleles with a given effect size and the total contribution across all effect sizes, but in this case these integrals do not simplify in any obvious way. A *Mathematica* notebook that performs these integrals numerically and was used to produce the nonlinear Lande approximations in ***Figs. S4.1***, ***S4.2*** and ***S4.4*** can be found at *https://github.com/sellalab/PolygenicAdaptation1D*.

We can now compare the results of the linear and nonlinear Lande approximations in order to ascertain when we should expect the linear approximation to be accurate. Notably, from ***Eq. S4.29***, we see that when $|a| \cdot \Lambda/V_A(0) \ll 1$ then $F_d(a, x_0) \approx \Lambda/V_A(0)$ and thus $x_d^{nL}(a, x_0) \approx x_d^{lL}(a, x_0)$, where superscripts $nL$ and $lL$ correspond to the nonlinear and linear Lande approximations, respectively. We therefore expect the linear approximation to be accurate when $|a| \cdot \Lambda/V_A(0) \ll 1$.

Our simulation results are consistent with this condition (***Fig. S4.2***), as when it is met, the linear and nonlinear approximations and simulations produce similar results. In cases with intermediate polygenicity and large shifts or low polygenicity and intermediate to large shifts, we see that the nonlinear approximation becomes substantially more accurate for intermediate effect size alleles (when $|a| \cdot \Lambda/V_A(0) \gtrapprox 1/2$; shaded gold region in ***Fig. S4.2B*** and ***C***). When polygenicity is low and both the shift and effects sizes are large, the nonlinear approximation can substantially overestimate the contribution to phenotypic adaptation (see the red box in ***Fig. S4.4A***), because many large effect alleles do not segregate long enough to feel the full integral effect of directional selection. However, when Lande's approximation applies, we expect such large effect alleles to be scarce and contribute negligibly to polygenic adaptation.

##### 4.4. The nonlinear non-Lande approximation

Lastly, we consider the case where both the linear expansions and Lande's approximation are inaccurate. We expect Lande's approximation to be inaccurate when alleles with large effect sizes contribute substantially to genetic variance before the shift, or more precisely when $C$ is not negligibly small (see *Section 4.1*). We expect the linear expansions to become inaccurate when polygenicity is low, the shift is large or the magnitude of effect size is large. More precisely, by analogy with our condition in the Lande case and as will become apparent below, we expect the linear approximation to become inaccurate when $(1 + A) \cdot |a| \cdot \Lambda/V_A(0)$ is not negligibly small. We derive an approximation for this case by combining the approaches that we used in the linear non-Lande and nonlinear Lande approximations. As the contribution of new mutations in the non-Lande case can be substantial, we consider both new mutations and standing variation.

###### 4.4.1. Standing variation

The nonlinear non-Lande approximation for standing variation is similar to the nonlinear Lande approximation. The only difference is that here the integral effect of directional selection is $\int_0^{t_*} D(\tau)\, d\tau = (1 + A) \cdot \int_0^\infty D_L(\tau)\, d\tau = (1 + A) \cdot \Lambda \cdot V_S/V_A(0)$ rather than $\Lambda \cdot V_S/V_A(0)$. By analogy, the instantaneous pulse approximation for the effect of directional selection is

$$\Delta x_d(a, x_0) = a x_0 (1 - x_0) \cdot F_d(a, x_0), \tag{S4.32}$$

where

$$F_d(a, x_0) \equiv \frac{1}{a}\left(\frac{\mathrm{Exp}\left[a \cdot \int_0^{t_*}(D(\tau)/V_S)d\tau\right] - 1}{1 + x_0\left(\mathrm{Exp}\left[a \cdot \int_0^{t_*}(D(\tau)/V_S)d\tau\right] - 1\right)}\right)$$
$$= \frac{1}{a}\left(\frac{\mathrm{Exp}\left[(1+A) \cdot a \cdot \Lambda/V_A(0)\right] - 1}{1 + x_0\left(\mathrm{Exp}\left[(1+A) \cdot a \cdot \Lambda/V_A\right] - 1\right)}\right). \quad \text{S4.33}$$

Based on ***Eq. S4.33*** and the same reasoning that we applied to the Lande approximations, we conclude that when $(1+A)\cdot|a|\cdot\Lambda/V_A(0) \ll 1$ then $\Delta x_d^{nN}(a, x_0) \approx \Delta x_d^{lN}(a, x_0)$ (where superscripts $nN$ and $lN$ correspond to the nonlinear and linear non-Lande approximations respectively), and we expect the linear non-Lande approximation to be accurate. The nonlinear non-Lande approximations for other quantities take the same form as in the corresponding nonlinear Lande approximations (e.g., as in ***Eqs. S4.30*** and ***S4.31***).

To test the nonlinear non-Lande approximation for standing variation, we first estimate the proportional contribution of standing variation to phenotypic change, $\eta$, using our AA simulations. We then solve for $A$ numerically by requiring that

$$2NU \cdot \Delta z_\infty^v(A) = 2NU \cdot \int \Delta z_\infty^v(a|A) \cdot g(a)da = \eta \cdot \Lambda, \quad \text{S4.34}$$

where $\Delta z_\infty^v(A)$ and $\Delta z_\infty^v(a|A)$ denote the nonlinear non-Lande approximations with a given value of $A$.

When we compare the nonlinear and linear non-Lande approximations for standing variation with simulation results (***Figs. S4.5*** and ***S4.7***), as we would expect from our condition for the accuracy of the linear approximation ($|a| \cdot (1+A) \cdot \Lambda/V_A(0) \ll 1$), we find the non-linear approximation can provide better estimates of fixation probability when $|a| \cdot (1+A) \cdot \Lambda/V_A(0) \gtrapprox 1/2$ (golden shaded regions in ***Fig. S4.5***). However, the linear and nonlinear non-Lande approximation perform comparably well for calculating the contribution to the change in mean (***Fig. S4.7***), since the error in the linear approximation cancels out somewhat in calculating the *difference* in fixation probabilities of opposite alleles, which is what generates the change in mean. For large shifts and low polygenicity, the nonlinear approximation can even perform worse than the linear one, because it overestimates the proportion contribution coming from large effect alleles (see the red box in ***Fig. S4.7***); and, consequently, also underestimates the proportion coming from small and intermediate effect alleles. This is because the approximation does not account for the fact that large effect alleles tend to segregate for fewer generations than those with small and moderate effects, and therefore experience fewer effective generations of directional selection.

###### 4.4.2. New Mutations

We derive the nonlinear approximation for new mutations by combining the approach that we used for new mutations in the linear non-Lande case with the one we used for standing variation in the nonlinear non-Lande case. The nonlinear instantaneous pulse approximation for the effect of directional selection on the frequency of a new mutation with effect size $a$ that arose $t$ generations after the shift in optimum is

$$\Delta x_d(a|t) = a\frac{1}{2N}\left(1 - \frac{1}{2N}\right) \cdot F_d(a|t), \quad \text{S4.35}$$

where

$$F_d(a|t) \equiv \frac{1}{a}\left(\frac{\mathrm{Exp}\left[a\cdot\int_t^{t+\Delta_t}(D(\tau)/V_S)\,d\tau\right]-1}{1+1/(2N)\cdot\left(\mathrm{Exp}\left[a\cdot\int_t^{t+\Delta_t}(D(\tau)/V_S)\,d\tau\right]-1\right)}\right)$$
$$\approx \frac{1}{a}\left(\mathrm{Exp}\left[a\cdot\int_t^{t+\Delta_t}(D(\tau)/V_S)\,d\tau\right]-1\right). \qquad \text{S4.36}$$

As we noted in *Section 4.2.2*, we expect that $a\cdot\Delta x_d(a|t)d \ll 1$ for most if not all new mutations. In this case, the first order Taylor expansion of the fixation probability in $\Delta x_d(a|t)$ remains accurate and we can approximate the expected long-term contribution of a pair of opposite mutations with effect sizes $\pm a$ by

$$\Delta z^*_\infty(a|t) = 2a\left(x_\infty(a|t) - x_\infty(-a|t)\right) \approx \frac{\partial\pi}{\partial x}(a,1/(2N))\cdot\Delta z^*_d(a|t)$$
$$\approx \frac{2v^*(a,1/(2N))}{v(a,1/(2N))}\cdot 2f(a)\cdot\Delta F_d(a|t)$$
$$\approx 1/(2N)\cdot 2f(a)\cdot\Delta F_d(a|t), \qquad \text{S4.37}$$

where

$$\Delta F_d(a|t) \equiv (F_d(a|t) - F_d(-a|t))/2 = \frac{1}{a}\mathrm{Sinh}\left(a\cdot\int_t^{t+\Delta_t}(D(\tau)/V_S)\,d\tau\right). \qquad \text{S4.38}$$

The expected contribution of such pairs per unit mutational input is then approximated by

$$\Delta z_\infty(a|t) \approx f(a)\left(\frac{1}{a}\mathrm{Sinh}\left(a\cdot\int_t^{t+\Delta_t}(D(\tau)/V_S)\,d\tau\right)/(2N)\right). \qquad \text{S4.39}$$

We can also write down integral forms for the marginal contribution of alleles with a given effect size and the total contribution across all effect sizes, but these integrals do not simplify in any obvious way.

We provided this approximation for completeness, but as we noted in *Section 4.2.2*, we expect the linear non-Lande approximation for new mutations to be accurate for considerably wider parameter ranges than it is for standing variation, possibly for the entire range permitted by our conditions (***App. A Table 2***).

###### 4.4.3. Standing variation and new mutations

We test the combination of the nonlinear non-Lande approximations for standing variation and linear non-Lande approximation for new mutations against simulation results, approximating their joint contribution by $\Delta z^v_\infty(a) + \Delta z^m_\infty(a)$ (***Eq. S4.27***). As we illustrate in ***Fig. S4.8***,the linear and nonlinear non-Lande approximations perform similarly well in most cases. However, as discussed in *Section 4.4.1*, when polygenicity is low or the shift is large, the nonlinear approximation for standing variation ($\Delta z^v_\infty(a)$) can overestimate the contribution coming from large effect alleles. Since the joint contribution is the sum of the contribution from standing variation and new mutations, the nonlinear non-Lande approximation for the joint contribution also overestimates the contribution coming from large effect alleles (see the red box in ***Fig. S4.8***).

#### 5. Simulations

We compared our analytic results to results from three kinds of simulations. The first type implements the full model. The second traces (AA) rather than individuals. Specifically, the frequency of an allele in the next generation follows a binomial distribution $x' \sim B\left(2N, E\left(x'\right)\right)$ with $E\left(x'\right)$ approximated by ***Eq. S1.9*** and the number of new mutations per generation follows a Poisson distribution with mean $2NU$. Tracing alleles rather than individuals in this way entails two potential sources of error. First, our approximation for $E\left(x'\right)$ replaces the distribution of background phenotypic contributions by its mean, thus neglecting the effects of variance in this distribution (*Section 1.1*). Second, we assume linkage equilibrium rather than free recombination, thus neglecting the effects of short-term linkage disequilibrium (Robertson, 1961).

The 3$^{\text{rd}}$ kind of simulations trace one allele at a time (OA). For alleles present before the shift in optimum, we sample the initial MAFs from the approximate closed form, equilibrium distribution (***Eq. S2.11***), using importance sampling based on the density of variance at different MAFs (***Eq. S2.12***). Alleles that arise after the shift start at frequency $1/(2N)$. Changes in frequency are modeled as we described for the AA simulations, but here we use the mean distance to the optimum, $\bar{D}(t)$, which is given from the outset. In the Lande case, we take the mean distance to equal the expectation under the Lande approximation (***Eq. 5***), which depends only on the initial phenotypic variance $V_A(0)$; this follows from the distribution of effect sizes and extent of polygenicity as described in ***Eq. S2.14***. In the non-Lande case, we estimate the mean distance by averaging over 2500 AA simulations with the same model parameters. OA simulations share the potential sources of error with the AA simulations, in addition to the potential errors arising from relying on the analytic approximation for the initial MAF and on the mean rather than on the actual (fluctuating) distance $D(t)$.

We use the AA and OA simulations throughout because they are considerably more computationally tractable, allowing us to obtain larger sample sizes and thus more precise comparisons with our analytic predictions. For example, running a single simulation, with the parameters of the non-Lande case used in the main text (see ***Fig. 2***), takes ∼340 hours with the full simulations and ∼10 with the OA simulations (for a burn-in period of $10N$ generations before the shift and a period of $12N$ generations after). Unfortunately, accurate estimates of fixation probabilities of alleles with a particular effect size can require hundreds or thousands (or even orders of magnitude more, if the effect size is extremely rare) of full or AA simulations. This problem is overcome by the OA simulations. For example, with the same parameters, running an OA simulation for 1000 aligned and 1000 opposing alleles with each of 13 different effect sizes (those used in many of the figures, e.g. ***Fig. S3.1***) takes ∼10 hours.

##### 5.1. Validating our main results with the full simulations

In ***Fig. S5.1***, we compare our main results about the phenotypic dynamics with those from the full simulation. Because of the computational intensity of the full simulation, we use only one set of parameters, the non-Lande case used in the main text (see ***Fig. 2***) with a shift of $\Lambda = 2\sqrt{V_A(0)}$, and run a sample of only 500 simulations, which is why our error is sizable. The phenotypic variance before the shift is greater in the full simulation than in our analytic approximation and AA simulations (***Fig. S5.1A***), plausibly because only the full simulation incorporates the effect

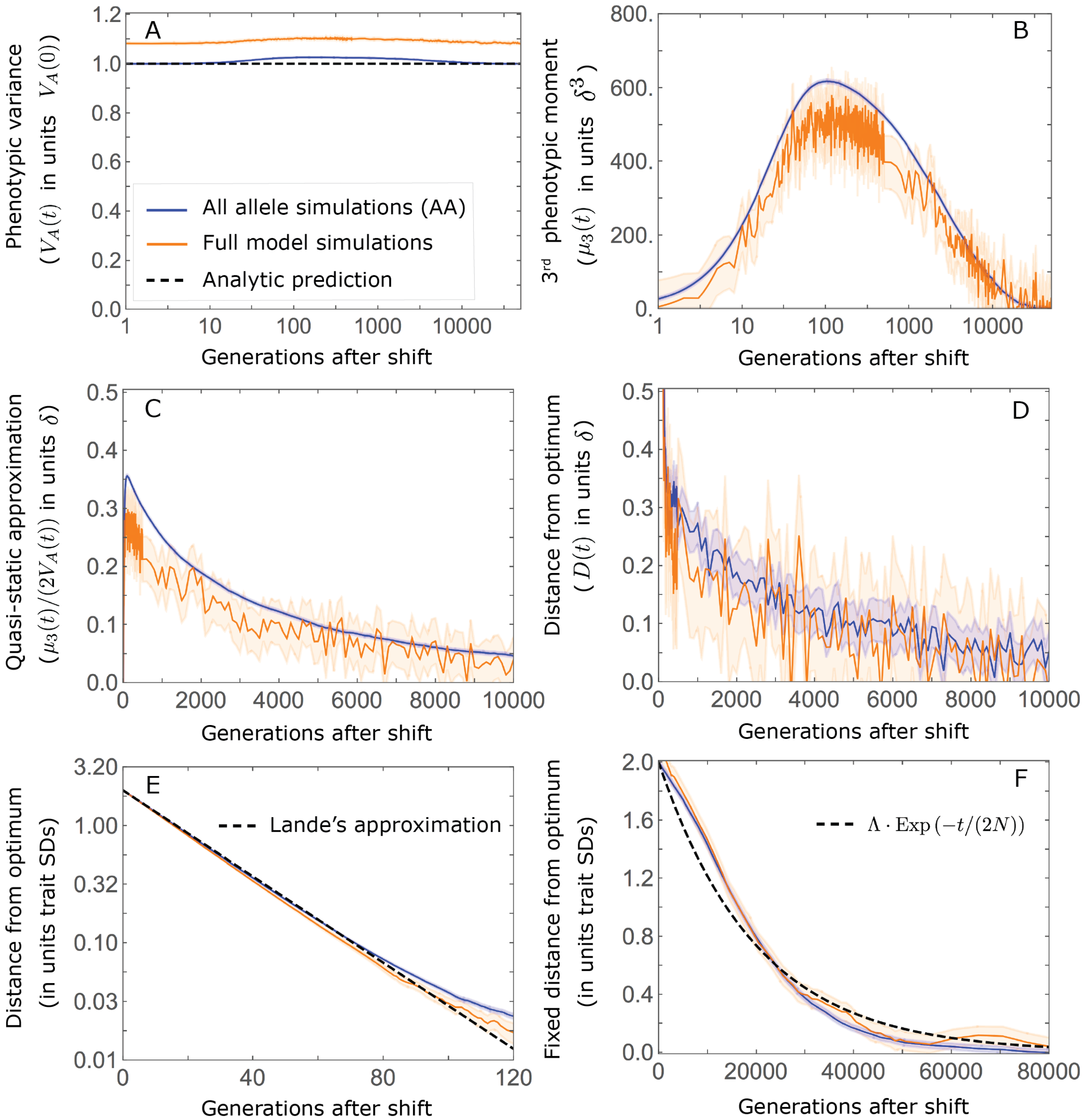

**Figure S5.1.** Comparison of our main results about the phenotypic dynamics with the corresponding results in the full simulations. The results shown are based on 2500 AA and 500 full simulations with the parameters of the non-Lande case used in the main text, i.e., $N = 10^4$, $U = 0.01$, $\Lambda = 2 \cdot \sqrt{V_A(0)}$ (with $\sqrt{V_A(0)} = 29 \cdot \delta$) and $a^2 \sim \Gamma(1, 16)$ $(E(a^2) = 16$ and $V(a^2) = 256)$, where effect sizes are measured in units of $\delta$. A) Phenotypic variance. B) $3^{rd}$ phenotypic moment. C) The quasi-static approximation for the mean phenotypic distance from optimum $(\mu_3(t)/(2V_A(t))$; ***Eq. 6***). D) The mean phenotypic distance from optimum. E) The mean phenotypic distance from optimum shortly after the shift (during the rapid phenotypic response). F) The fixed distance from the optimum.

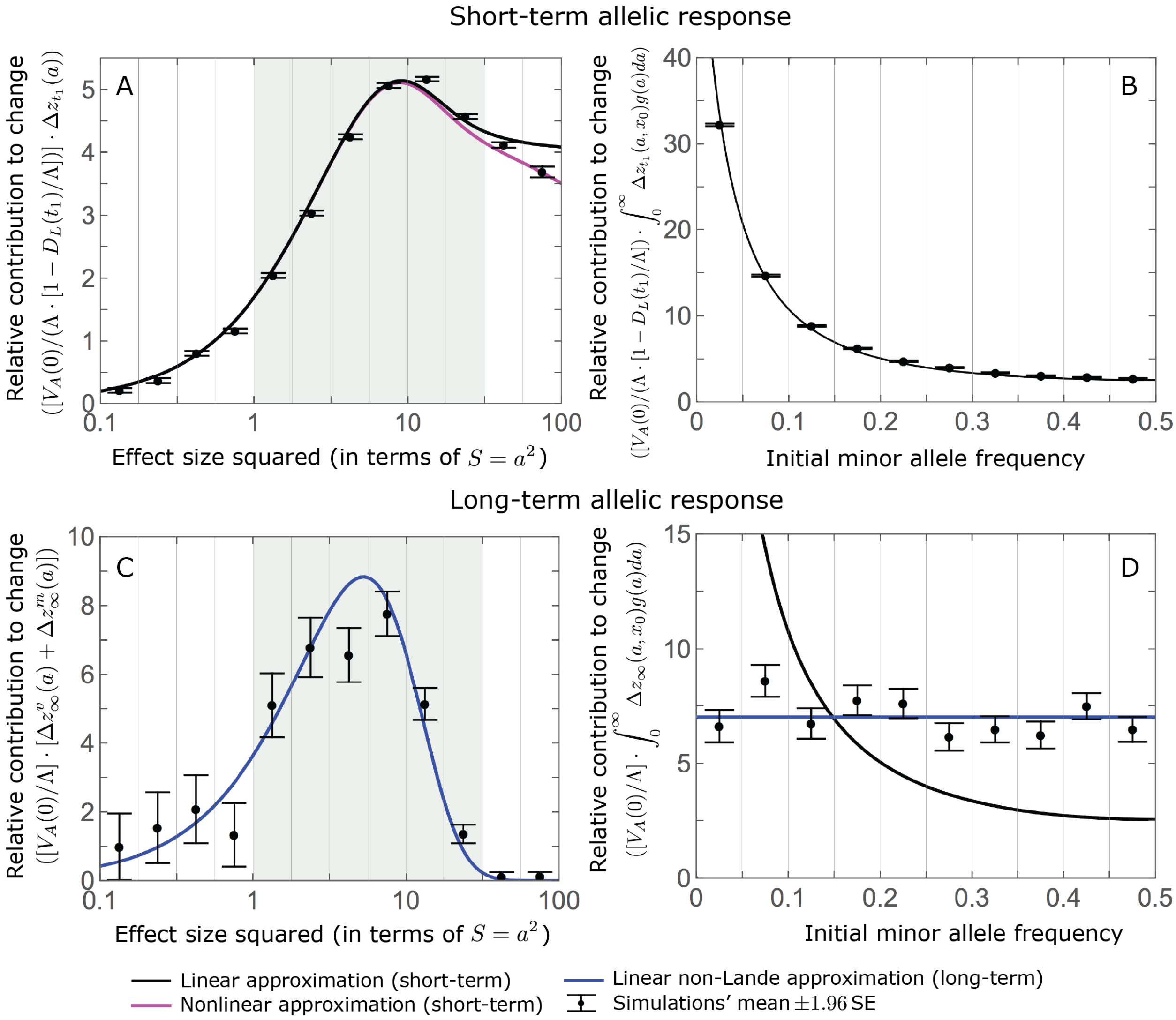

**Figure S5.2.** Comparison of our main results about the allelic dynamics with the corresponding results of the full simulations. The model parameters and full simulations are the same as in ***Fig. S5.1***. We calculate the relative contribution of alleles in each effect size bin (between the gray gridlines) by dividing the contribution of all fixations in the bin by the mutation rate per generation corresponding to that bin. We show the relative phenotypic contribution per unit mutational input of alleles as a function of (squared) effect size (A and C) and initial MAF (B and D) in the rapid (top) and equilibration (bottom) phases.

of variance in the phenotypic background of a focal allele. Variance in the phenotypic background reduces the efficacy of stabilizing selection—akin to how environmental contributions to phenotypic variance effectively increase $V_S$ (Turelli, 1984; Bürger, 2000)—which increases alleles' initial MAFs and contributions to phenotypic variance. (A similar slight underestimation of genetic variance at steady-state is seen in Fig. A6(a) of Simons et al. (2018).)

The same effect plausibly accounts for most of the other small phenotypic differences between the full and AA simulations. The short-term term phenotypic response is slightly faster in the full than in the AA simulations (***Fig. S5.1E***), because of the greater initial phenotypic variance in the full model. The increase in the 3rd phenotypic moment is smaller in the full than in the AA simulations (***Fig. S5.1B***), possibly because the greater initial phenotypic variance allows the same shift in mean phenotype to be achieved by smaller changes in allele frequency (e.g., ***Eq. 10***). Lastly, the greater variance and smaller 3rd moment in the full simulations plausibly explain the smaller distance from the optimum during the prolonged phase in the full relative to the AA simulations, seen both directly and in the quasi-static approximation (***Fig. S5.1D*** and ***C***). Small differences notwithstanding, the results of the full simulations support all of our main results about the phenotypic dynamic. The same is true of our main analytic results about the allelic dynamics (***Fig. S5.2***).

#### 5.2. Choice of simulation parameters

Our simulation parameter values were primarily chosen to span as wide a range as possible (***Fig. S5.3***) given our conditions on them (***App. A Table 2***). The one exception is our choice of a population size of $N = 10000$, which was motivated by being on the order of the effective size in human populations (Schiffels and Durbin, 2014), though on the lower side in order to allow for greater computational tractability. For a given population size, the choice of mutation rate per haploid genome per generation, $U$, is constrained from below by the assumption that the trait is highly polygenic, i.e., $\sqrt{2NU} \gg 1$ (***App. A Table 2***). Consequently, the lowest mutation rate that we used was U =0.005, corresponding to our low end of polygenicity, i.e., $\sqrt{2NU} = 10 \gg 1$. The choice of mutation rate is also constrained from above by the assumption that $U \ll 0.2$ (***App. A Table 2***). The highest mutation rate that we used was therefore $U = 0.02 \ll 0.2$, corresponding to our high end of polygenicity, i.e., $\sqrt{2NU} = 20$. As an intermediate value, we used $U = 0.01$, corresponding to an extent of polygenicity of $\sqrt{2NU} \approx 15$ (***Fig. S5.3A***).

The choice of the effect size (measured in units of $\delta$) distribution of new mutations is constrained by the requirement that a substantial proportion of them are not effectively neutral, i.e., $a^2 \gtrapprox 1$ (***App. A Table 2***). In simulations representative of the Lande case, we further require that most new mutations satisfy $a^2 \lessapprox 4$ (see ***Eq. S4.11***). Our choice of $a^2 \sim \Gamma(1,1)$ ($E(a^2) = V(a^2) = 1$) satisfies both requirements, with proportion 0.63 of mutations with $a^2 \leq 1$, 0.35 with $1 < a^2 \leq 4$, and 0.02 with $a^2 > 4$ (***Fig. S5.3C***). In simulations representative of the non-Lande case, we require that a substantial proportion of mutations have effect sizes $a^2 > 4$ while being much smaller than the width of the fitness function, i.e., $|a|/\sqrt{V_S} \ll 1$ (***App. A Table 2***). Our choice of $a^2 \sim \Gamma(1, 16)$ ($E(a^2) = 16$ and $V(a^2) = 256$) satisfies these requirements, with proportion 0.78 of mutations with $a^2 > 4$ and only $4 \times 10^{-6}$ with $a^2 > V_S/100$.

Lastly, we require the shift in optimum to be smaller than the width of the fitness function, i.e., $\Lambda \lessapprox \sqrt{V_S}$ (***App. A Table 2*** and ***Fig. S5.3B***). As we detail in *Section 2.2*, this requirement constrains how large the shift can be relative to the phenotypic standard deviation ($\sqrt{V_A(0)}$) in a manner that depends on the mutation rate and effect size distribution: $\Lambda/\sqrt{V_A(0)} \leq 1/\sqrt{U}/\sqrt{\int_0^\infty v(a)\, g(a)da}$ (***Eq. S2.15*** and ***Fig. S2.2***). As our largest shift, we chose $\Lambda = 4\sqrt{V_A(0)}$, which, for our highest mutation rate ($U = 0.02$), is near the upper bound given our choices of effect size distributions for the Lande and non-Lande cases (with $\Lambda/\sqrt{V_S} = 0.7$ and 1.2, respectively). As a smaller shift, we used $\Lambda = 2\sqrt{V_A(0)}$.

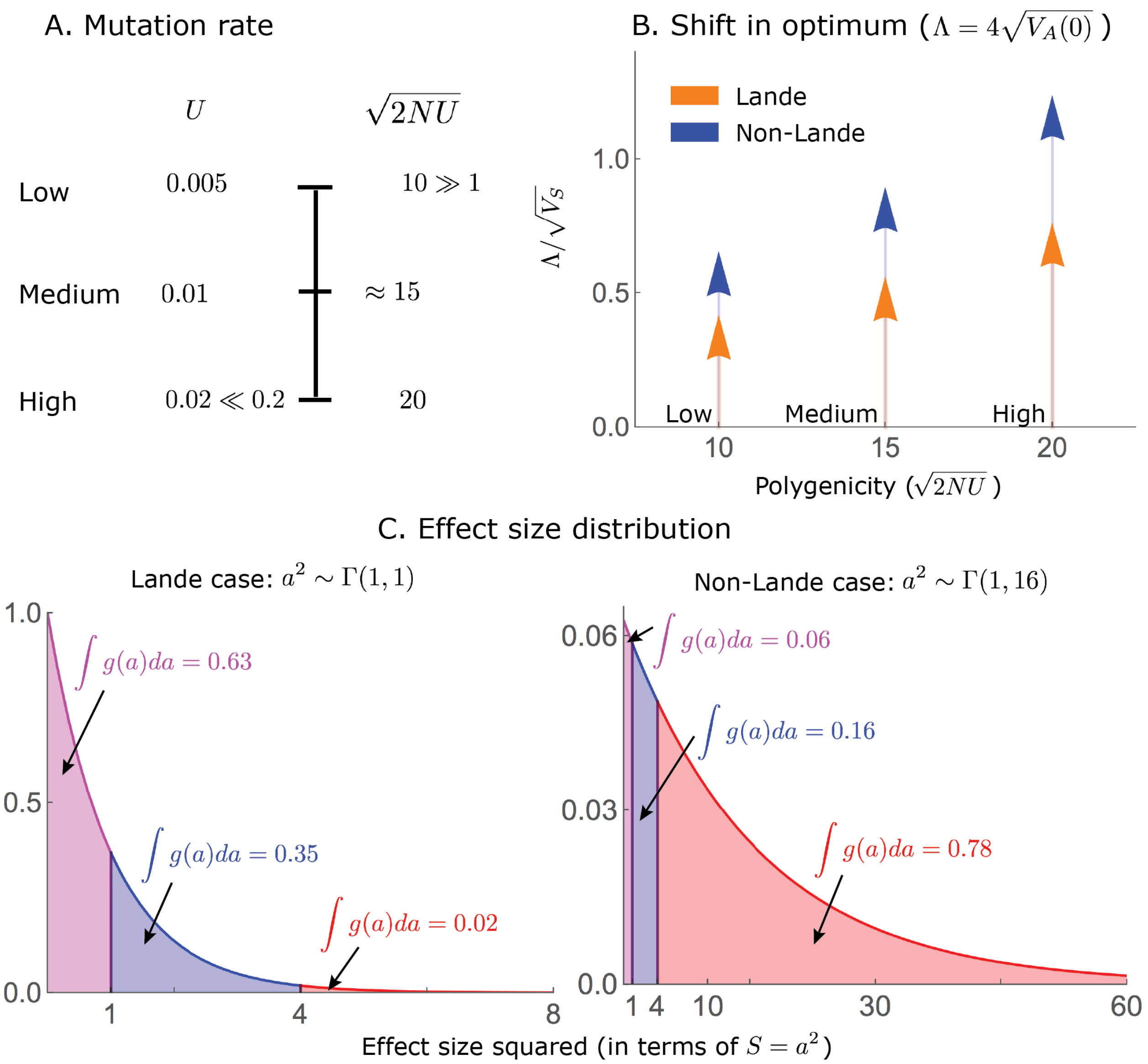

**Figure S5.3.** Our simulation parameter values are chosen to span as wide a range as possible given our conditions on them.

#### 6. Phenotypic deviations from Lande's approximation are determined by $C$

Here, we revisit the phenotypic response to selection to articulate three conjectures that extend the analysis in the main text. The phenotypic response affects the allelic dynamics primarily through the mean distance from the optimum, $D(t)$, as described in our allelic equation (***Eq. S1.1***). The allelic dynamics are insensitive to rapid stochastic fluctuations in $D(t)$, because selection effects on allele frequencies are small in a single generation, such that, in effect, alleles 'feel' the average $D(t)$ over several generations. We therefore focus on integrals of the form $\int_0^t D(\tau)/V_S d\tau$, which is how $D(t)$ features in our allelic approximations, and on the quasi-static approximation $D(t) \approx \mu_3(t)/(2V_A(t))$ for the distance during equilibration, both of which are less noisy than $D(t)$

itself. We continue to measure the trait value in units of $\delta$ but sometimes include $\delta$ in equations for clarity.

While variation in each of the model parameters affects the trajectories of the mean distance (or its integral), a few compound parameters appear most influential (***Fig. S6.1***). During the rapid phase, the distance is well approximated by Lande's solution $D_L(t)$, as illustrated for a couple of model parameter choices in ***Fig. 2B*** and for a dozen in ***Fig. S6.1B*** (where the integral over the distance scaled by the integral obtained under Lande's approximation stacks up around 1 during the rapid phase). During the equilibration phase, the distance from the optimum declines very slowly in non-Lande cases, resulting in a prolonged increase in the integral over the distance (***Fig. S6.1A*** and ***B***). Nevertheless, for a given amplification $C$, the integral over the distance scaled by the integral obtained under Lande's approximation appears to be insensitive to the values of other parameters (***Fig. S6.1B***).

The trajectories of the quasi-static approximation further support the notion that, after normalization based on Lande's approximation, $D(t)$ during equilibration depends primarily on $C$ (***Fig. S6.1C*** and ***D***). In fact, simulations support our 1$^{\text{st}}$ conjecture that

$$D(t) \approx \mu_3(t)/V_A(t) \approx (k(t)\cdot\delta)\cdot C\cdot(\Lambda\cdot\delta)/V_A(0), \qquad \text{S6.1}$$

where $(\Lambda\cdot\delta)/V_A(0) = \int_0^\infty ((D_L(\tau)\cdot\delta)/V_S)\,d\tau$ is the Lande-based normalization, and $k(t)$ is a proportionality coefficient that has no units. Simulations suggest that $k(t) \lessapprox 5$ (see, e.g., ***Fig. S6.1D***), and our 2$^{\text{nd}}$ conjecture suggests that $\int_{t_1}^\infty k(\tau)d\tau \approx 2N$ (***Eq. S6.2*** below). One important implication of ***Eq. S6.1*** is that at least under our assumptions on parameters, the distribution of effect sizes alone, and specifically $C$, determines whether the phenotypic response deviates substantially from Lande's approximation. Specifically, under our conditions on parameters $\Lambda/V_A(0) \lessapprox 1/2$, and assuming that $C \ll 1$, even large shifts are insufficient to generate appreciable deviations from Lande's approximation during equilibration.

Our 2$^{\text{nd}}$ conjecture concerns the long-term integral distance $\int_0^\infty (D(\tau)\cdot\delta/V_S)\,d\tau$. We have seen that in the Lande case, when $C \ll 1$, then $\int_0^\infty (D(\tau)\cdot\delta/V_S)\,d\tau \approx \int_0^\infty (D_L(\tau)\cdot\delta/V_S)\,d\tau = (\Lambda\cdot\delta)/V_A(0)$. So far, we have not considered $\int_0^\infty (D(\tau)\cdot\delta/V_S)\,d\tau$ in the non-Lande case. (We explained our non-Lande allelic approximations in terms of $\int_0^{t_*} (D(\tau)\cdot\delta/V_S)\,d\tau$, where $t_*$ was the effective number of generations over which alleles are subject to directional selection; e.g., ***Eq. 23***.) Our simulations support our 2$^{\text{nd}}$ conjecture, that

$$\int_0^\infty (D(\tau)\cdot\delta/V_S)\,d\tau \approx (1+C)\cdot\int_0^\infty (D_L(\tau)\cdot\delta/V_S)\,d\tau = (1+C)\cdot(\Lambda\cdot\delta)/V_A(0) \qquad \text{S6.2}$$

(***Figs. S6.1B*** and ***S6.2A***).

Our 3$^{\text{rd}}$ conjecture concerns the long-term phenotypic contribution of fixations of new mutations (i.e., that arise after the shift). The results of simulations suggest that the proportional contribution of new mutations, $1-\eta$, depends primarily on the distribution of effect sizes, and specifically on $C$. In fact, it appears that, to a good approximation, the proportional contribution of new mutations

$$1-\eta \propto C, \qquad \text{S6.3}$$

and is thus insensitive to the values of other parameters, e.g., to the extent of polygenicity and shift size (***Fig. S6.2B***). In summary, we posit that the parameter $C$, which depends only on the mutational distribution of effects sizes and arose from our analysis of the allelic response, largely determines the deviations of the phenotypic dynamics from Lande's approximation.

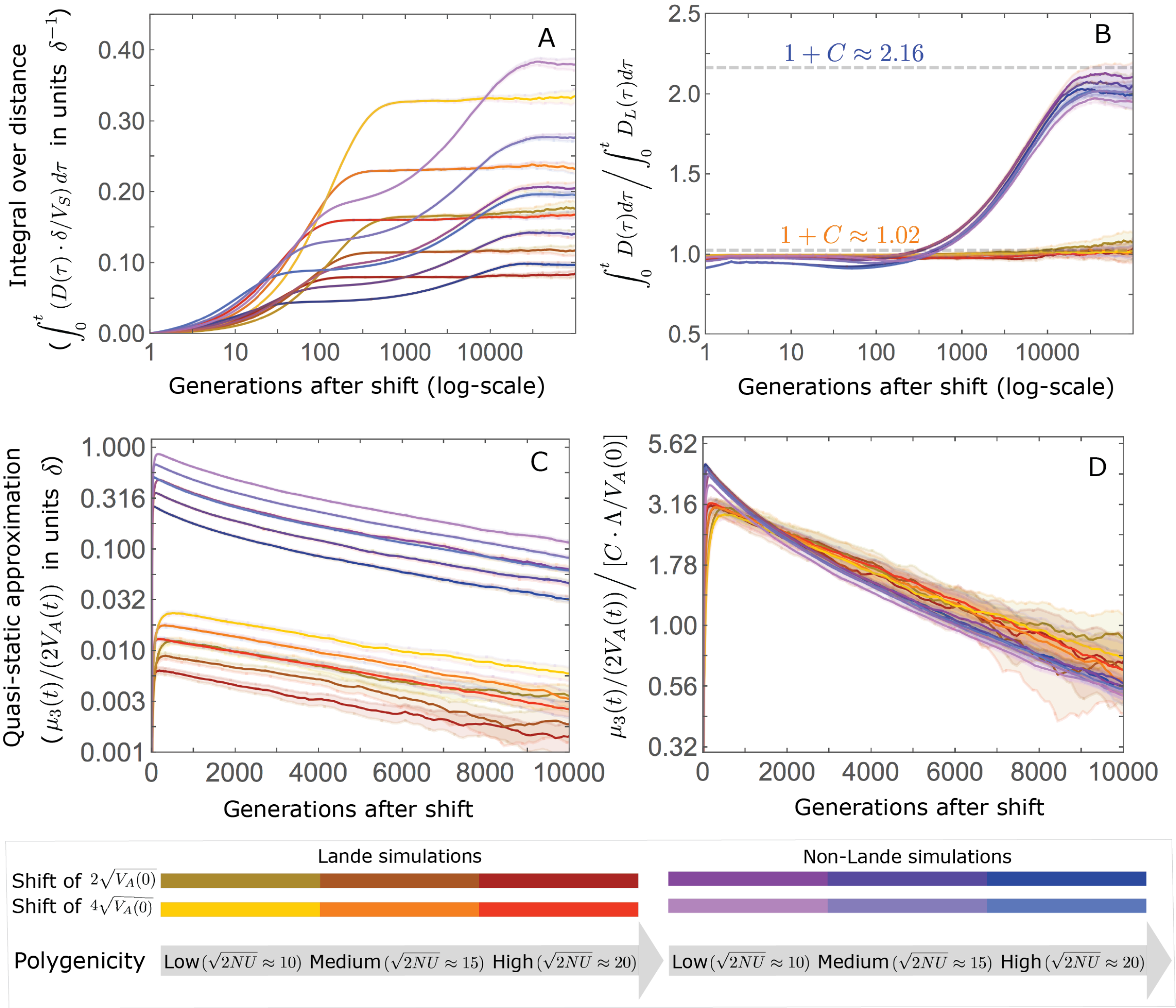

**Figure S6.1.** The trajectory of the mean integral over the phenotypic distance from the optimum is largely determined by Lande's approximation and by the amplification $C$. A) The integral distance from the optimum as a function of time after the shift. In Lande cases, the integral nears the asymptote $\int_0^\infty (D_L(\tau)\cdot\delta/V_S)\,d\tau = (\Lambda\cdot\delta)/V_A(0)$ at the end of the rapid phase, whereas in non-Lande cases, the integral continues to increase substantially during equilibration. B) The integral over the distance scaled by the integral over the distance obtained under Lande's approximation as a function of time. The phenotypic trajectories corresponding to the Lande and to the non-Lande cases with different parameter values stack up, illustrating that, to a good approximation, the impact of polygenicity and shift size are captured by Lande's approximation; the long-term impact of the effect size distribution, manifest in the non-Lande cases, is largely captured by $C$. C) The quasi-static approximation of the distance as a function of time is affected by all of the parameters. D) The quasi-static approximation of the distance scaled by $C\cdot(\Lambda\cdot\delta)/V_A(0)$ as a function of time. During the equilibration phase, the phenotypic trajectories corresponding to different parameter values stack up, illustrating that, to a good approximation, the effects of all of the parameters is captured by this compound parameter. The simulation results were averaged over 2500 runs of our AA simulations assuming $N = 10^4$, with our standard parameter values for the Lande and non-Lande cases (*Section 5.2*) and with the extent of polygenicity and shift sizes specified in the legend.

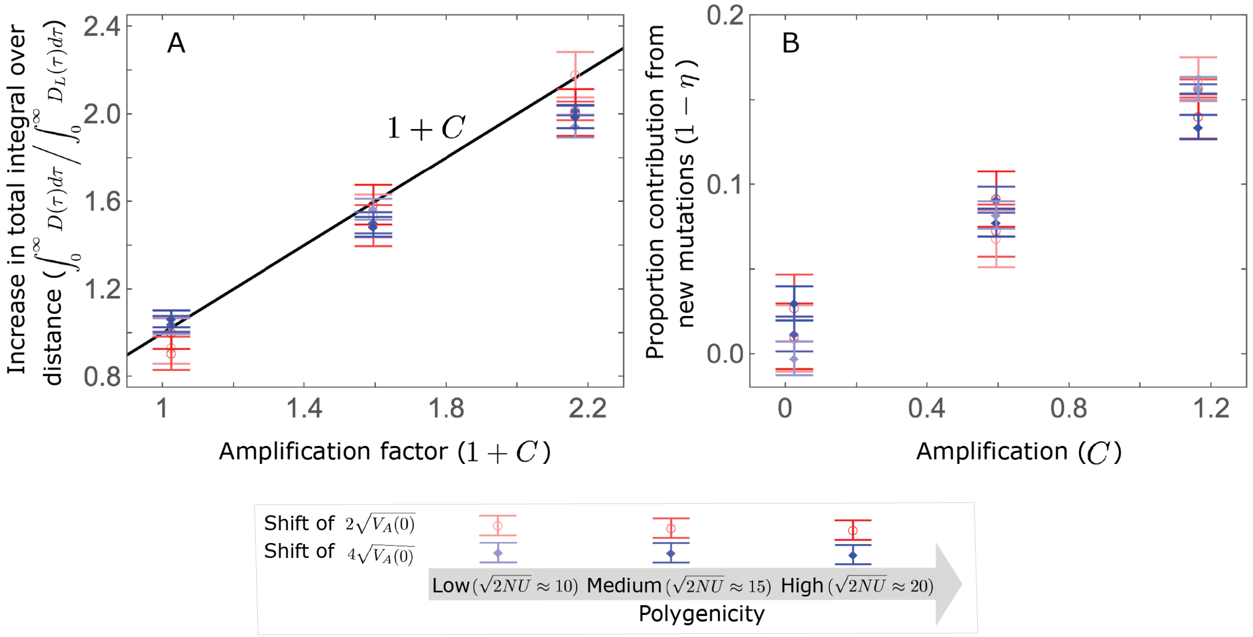

**Figure S6.2.** The amplification $C$ largely determines the increase in total integral distance, where $\int_0^\infty D(\tau)d\tau / \int_0^\infty D_L(\tau)/V_S d\tau \approx 1 + C$ (A); moreover, the proportion of long-term phenotypic change arising from new mutations $(1 - \eta)$, is approximately proportional to $C$ (B). As an illustration, we show the averages of these quantities over 500 AA simulations assuming $N = 10^4$, with the three extents of polygenicity and two shift sizes specified in the legend, and with three distributions of effect sizes: a Lande case with $a^2 \sim \Gamma(1, 1)$ corresponding to $C \approx 0.02$, a non-Lande case with $a^2 \sim \Gamma(2/9, 27)$ corresponding to $C \approx 0.59$, and a non-Lande case with $a^2 \sim \Gamma(1, 16)$ corresponding to $C \approx 1.16$.

#### 7. Additional figures

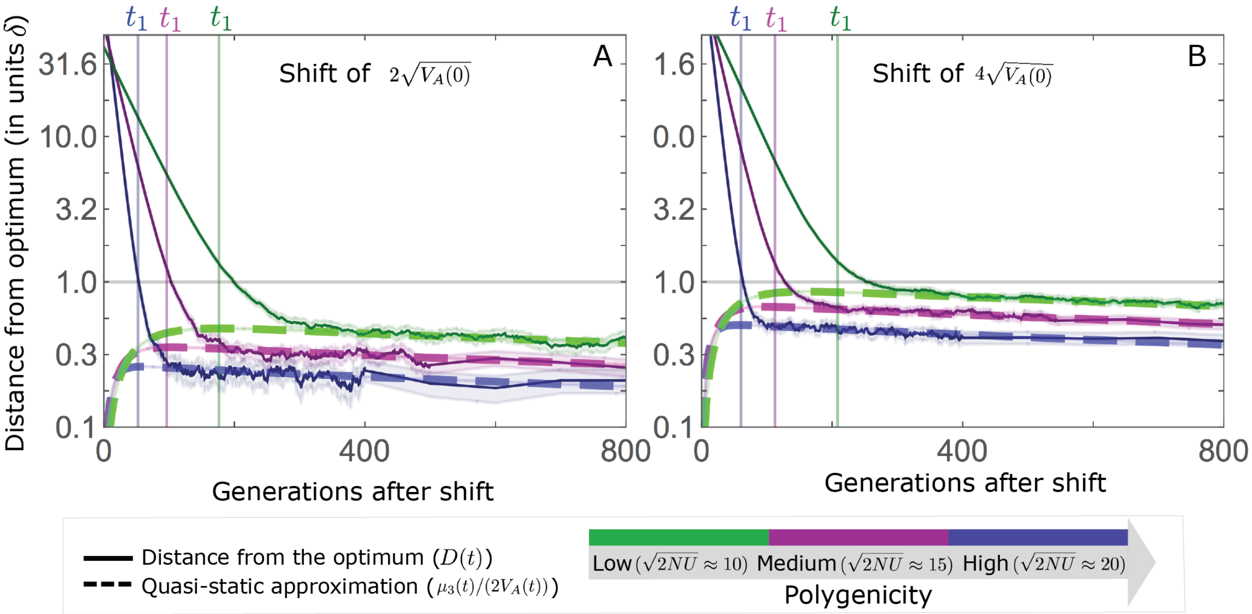

**Figure S7.1.** Our definition of the end of the rapid phase, $t_1$, roughly captures the change in phenotypic dynamic in the non-Lande case. The results shown were averaged over 2500 AA simulations, assuming $N = 10^4$, with the parameters of our standard non-Lande example (*Section 5.2*) and the specified extents of polygenicity and shift sizes. We defined $t_t$ as the time at which $D_L(t_1) = \delta$ (***Eq. 9***). Here we see that, at least in these non-Lande cases, $D_L(t_1)$ is quite close to $\delta$ as well. We also see that the quasi-static approximation (***Eq. 6***) becomes accurate shortly after $t_1$.

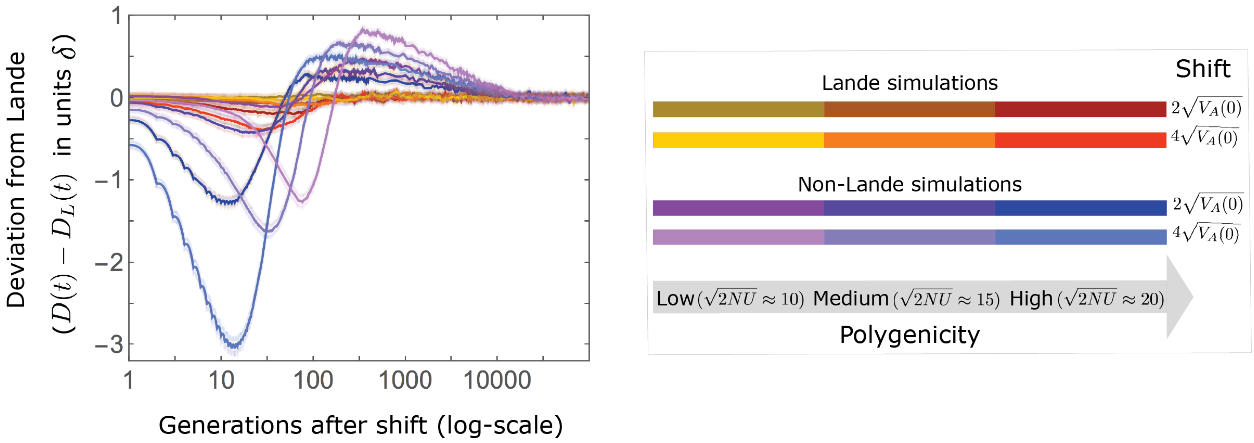

**Figure S7.2.** Deviations from Lande's approximation as a function of time after the shift. In the Lande case, i.e., when $C \ll 1$, the mean distance from the new optimum is well approximated by $D_L(t)$. Deviations increase with shift size and extent of polygenicity (as seen, e.g., in the red curve), primarily due to a moderate increase in phenotypic variance during the rapid phase. The increase in the $3^{\text{rd}}$ phenotypic moment is minimal, which is why, in this case, there are no substantial long-term deviations. In the non-Lande case, the distance from the optimum decays faster than predicted by Lande's approximation $(D(t) - D_L(t) < 0)$ during the rapid phase, because of the increase in phenotypic variance. The increase in the $3^{\text{rd}}$ phenotypic moment during the rapid phase leads to a slower decay than predicted by Lande's approximation during the equilibration phase. Even in the non-Lande case, however, we always find the distance from the optimum during equilibration to be smaller than $\delta$. The simulation results were averaged over 2500 runs of our allelic simulations (AA) with our standard parameter values for the Lande and non-Lande cases (*Section 5.2*) and with the extent of polygenicity and shift sizes specified in the legend.

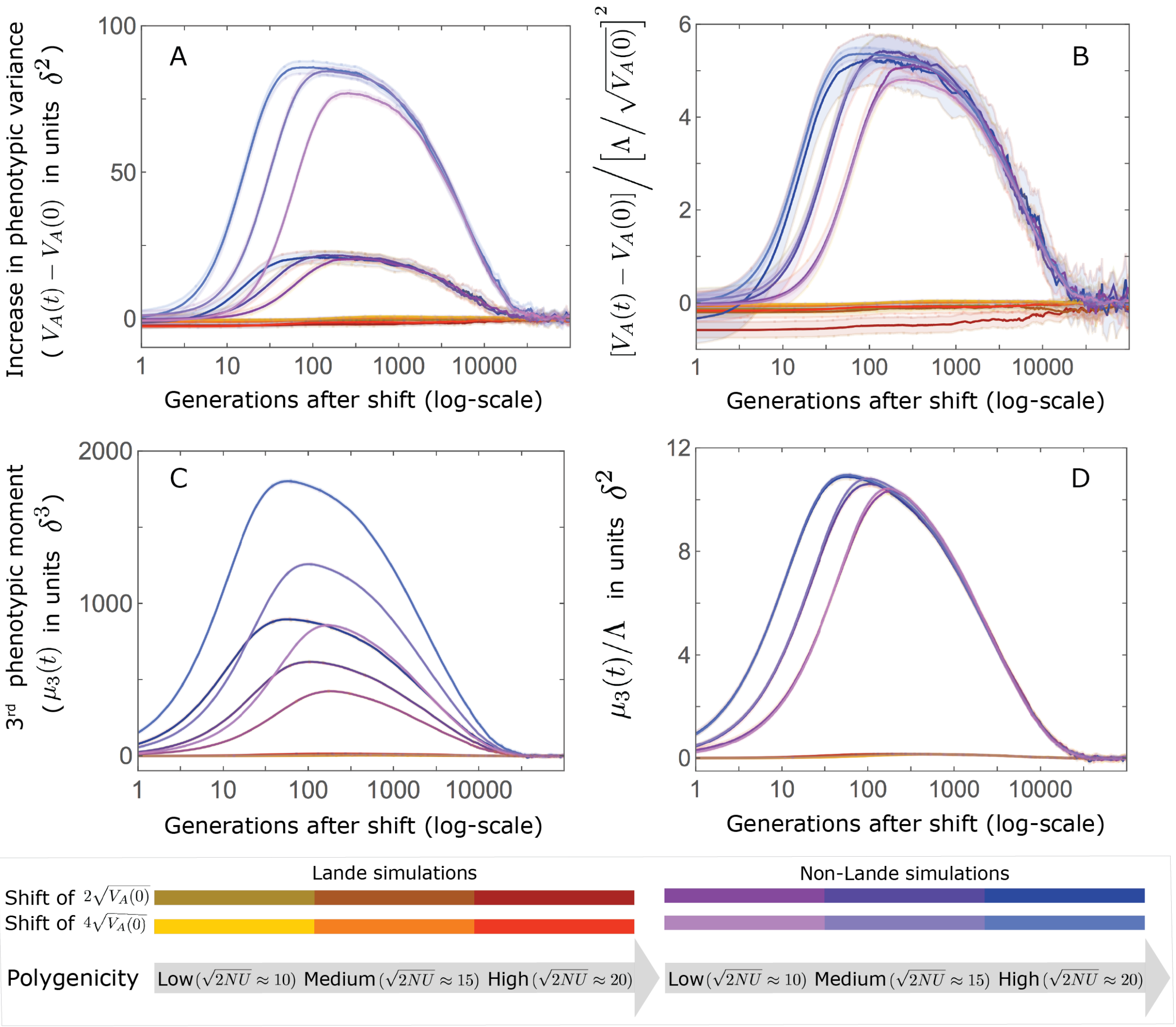

**Figure S7.3.** The increase in the 2nd and 3rd moments of the phenotypic distribution. (A and C) The increase is largely driven by alleles with large effects. The increase is largely driven by alleles with large effects. Specifically, there are substantial increases only in the non-Lande cases, i.e., those with an abundance of new mutations with $a^2 > 4$. (B and D) We conjecture that the compound parameters that largely determine the increase in moments are: $(\Lambda/\sqrt{V_A(0)})^2$ for the 2nd moment (B) and $\Lambda/\delta$ for the 3rd. Note that the slight differences among curves in the non-Lande case are short-lived (time is measured on a log-scale). The model parameters and simulations are the same as in ***Fig. S7.2***.

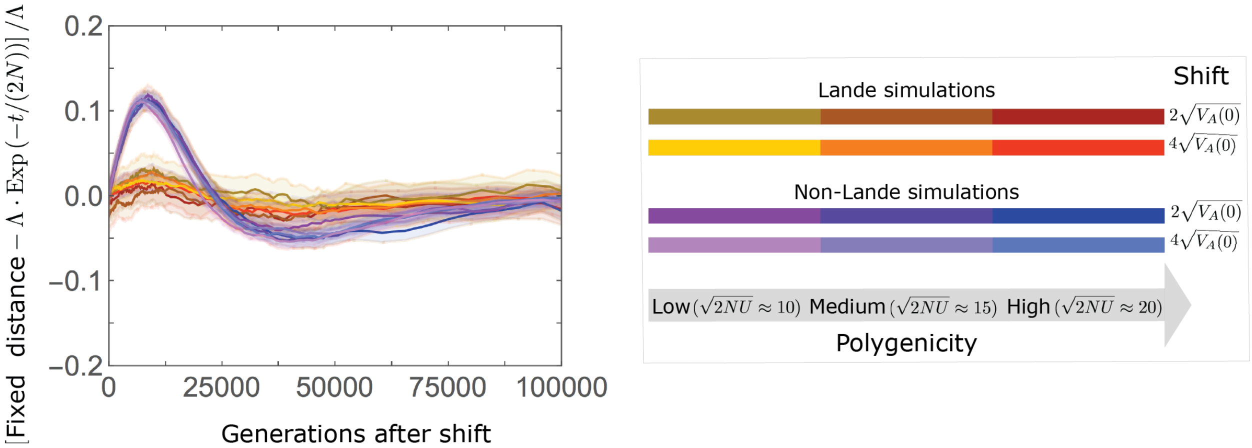

**Figure S7.4.** The approach of the fixed background to the new optimum is well approximated by $\Lambda e^{-t/(2N)}$, with deviations proportional to the shift size multiplied by a function that depends largely on $C$. We speculate that the (positive) deviations on the intermediate time-scale (e.g., around generation 1000) in the non-Lande case are primarily due to the slower accumulation of fixations from new mutations, which contribute substantially in this case, whereas the (negative) deviations on the slightly longer time-scale (e.g., around generation 4000) are due to the cumulative effect of prolonged weak directional selection, which has a substantial effect in this case. The model parameters and simulations are the same as in ***Fig. S7.2***.
